# A Type 2 Innate Lymphoid Cell-Interleukin 9 Circuit Induces Paneth Cell Metaplasia and Small Intestinal Remodeling

**DOI:** 10.1101/2022.11.02.514603

**Authors:** Cheng-Yin Yuan, Aditya Rayasam, Alison Moe, Wenwen Xu, Aniko Szabo, Michael Hayward, Clive Wells, Nita Salzman, William R. Drobyski

**Affiliations:** Department of Medicine, Blood and Marrow Transplant and Cellular Therapy Program, Medical College of Wisconsin, Milwaukee, WI; Department of Pediatrics, Medical College of Wisconsin, Milwaukee, WI; Department of Microbiology and Immunology, Medical College of Wisconsin, Milwaukee, WI; Division of Biostatistics, Institute of Health and Equity, Medical College of Wisconsin, Milwaukee, WI

**Author notes:** Address correspondence to: William R. Drobyski, M.D., Blood and Marrow Transplant and Cellular Therapy Program, Medical College of Wisconsin, 9200 West Wisconsin Avenue, Milwaukee, WI 53226.

## Abstract

Paneth cell metaplasia (PCM) typically arises in the setting of pre-existing gastrointestinal (GI) diseases; however, the mechanistic pathway that induces metaplasia and whether PCM is initiated exclusively by disorders intrinsic to the GI tract has not been delineated. Herein, we describe the development of PCM in a murine model of inducible bcr/abl oncogene-driven chronic myelogenous leukemia (CML) in which metaplasia is causally and temporally linked to the development of leukemia. Mechanistically, CML induced a proinflammatory state within the GI tract that resulted in the production of epithelial-derived IL-33. Binding of IL-33 to ST2 led to the production of IL-9 by type 2 innate lymphoid cells (ILC2s) which was directly responsible for the induction of PCM in the colon and Paneth cell hyperplasia in the ileum. In addition, IL-9 directed remodeling of the small intestines characterized by goblet and tuft cell hyperplasia along with the expansion of mucosal mast cells. Thus, we identify that an extra intestinal disease can trigger an ILC2/IL-9 immune circuit which induces PCM and regulates epithelial cell fate decisions in the GI tract.

## INTRODUCTION

Paneth cells are specialized secretory epithelial cells that are generated from leucine-rich-repeat-containing G protein coupled receptor 5 (Lgr5) positive stem cells and are located at the bottom of crypts in the small intestines (1, 2). Signaling through the wnt pathway induces the subsequent maturation of these cells within intestinal crypts (3). Paneth cells produce large quantities of antimicrobial peptides (AMPs), inflammatory mediators, and signaling molecules (4) and hence play a pivotal role in maintaining a homeostatic balance between commensal bacteria and the host immune system (5). The production of AMPs, which include α-defensins, lysozyme and secretory phospholipase A2 (sPLA2), are particularly important for the maintenance of intestinal immune homeostasis and the shaping of microbial composition within the small intestines (6). Of the AMPs produced by Paneth cells, α-defensins (also called crypt defensins or cryptdins in the mouse) are the most abundant of these secretory proteins (7). They are synthesized as inactive propeptides and then activated after cleavage by metalloproteinase 7 where they regulate the microbial ecosystem (6).

Numerous studies have shown that the disruption of Paneth cell function play a key role in many diseases such as inflammatory bowel disease and graft versus host disease (8–11). The damage to gut epithelial and Paneth cells in these disease states results in the loss of AMP production as well as a reduction in microbial diversity. Interestingly, in some disease settings, Paneth cells can also appear outside of the natural environment of the small intestines, a phenomenon which has been termed Paneth cell metaplasia (PCM). Sites of metaplasia are typically in the colon, stomach and esophagus and most often are associated with pathological conditions involving the GI tract, such as inflammatory bowel disease, Barrett’s esophagus and colorectal cancer (12–15). A hallmark of PCM is that it typically arises in the setting of chronic inflammation and pathological damage in the GI tract; however, the mechanistic pathway by which PCM develops under these conditions remains unknown.

In the current study, we report the development of PCM in a murine model of chronic myelogenous leukemia (CML) which is driven by the hematopoietic stem cell-restricted expression of the bcr/abl oncogene. In addition, we demonstrate extensive remodeling of the small intestines characterized by marked lengthening along with goblet and tuft cell hyperplasia. Mechanistically, PCM was attributable to intestinal epithelial cell-induced IL-33 production that arose in the setting of CML-induced gastrointestinal (GI) tract inflammation. This led to the activation of type 2 innate lymphoid cells (ILC2s) resulting in interleukin-9 (IL-9)-driven PCM, mucosal mast cell hyperplasia, and extensive remodeling of the small intestine characterized by marked lengthening along with goblet and tuft cell hyperplasia. Thus, intestinal inflammation can trigger an ILC2/IL-9 circuit that induces metaplasia in Paneth cells and regulates the development of specialized epithelial cell populations resident in the GI tract.

## RESULTS

### Paneth cell metaplasia emerges during bcr/abl oncogene dependent CML

We employed a murine bcr-abl oncogene-dependent chronic myelogenous leukemia (CML) model that recapitulates the development and clinical manifestations of human disease (16). Specifically, cross breeding of SCLtTA and bcr-abl mice results in double transgenic animals in which expression of the bcr-abl oncogene and the subsequent development of CML is regulated by tetracycline (Tet) administration (**Supplemental Figure 1A**). To standardize disease kinetics within a reproducibly defined temporal window, we developed a murine CML transplantation model (17) in which lethally irradiated wild type FVB mice are transplanted with bone marrow (BM) cells from SCLtTA/bcr-abl animals (henceforth referred to as CML mice) and then maintained on or off Tet. Recipients maintained off Tet exhibited weight loss (**Supplemental Figure 1B**), along with a significant temporal increase in white blood cells (**Supplemental Figure 1C**) and granulocytes (**Supplemental Figure 1D**) in the peripheral blood. These animals also had a nearly two-fold augmentation in spleen weights (**Supplemental Figure 1E**) and an increase in the frequency (**Supplemental Figure 1F**) and absolute number of CD11b^+^ Gr-1^+^ myeloid cells (**Supplemental Figure 1G**) consistent with what is observed in CML in humans (18, 19). There was also loss of normal follicular architecture in the spleen and mesenteric lymph nodes due to the expansion of myeloid cells in these sites (**Supplemental Figures 1H and 1I**). In addition, histological analysis of non-lymphoid tissues revealed accumulation of neutrophils in the liver and lung of leukemic animals (**Supplemental Figures 1J and 1K**).

Unexpectedly, during the histological examination of the colon, we observed that CML mice had a prominent number of cells with cytoplasmic eosinophilic granules that were morphologically consistent with Paneth cells (**Figure 1A**). In addition, whereas Paneth cells in the ileum of non-leukemic animals were located appropriately in the basal region of crypts, Paneth cells in the ileum of CML mice were scattered throughout the villi (see arrows **Figure 1A**). Staining of colonic tissue with phloxine tartrazine revealed characteristic Paneth cell granule affinity for phloxine (**Figure 1B**). In addition, electron microscopy also confirmed that these were Paneth cells as evidenced by the presence of spherical electron dense granules (**Figure 1C**) and tight junctions which identified these cells as epithelial and not haematopoietically-derived (**Figure 1D**). There was a significant increase in the absolute number of Paneth cells in both the colon and ileum when compared to animals maintained on Tet alone (**Figures 1E and 1F**), indicative of Paneth cell metaplasia (PCM) in the colon and hyperplasia in the ileum. Analysis of antimicrobial peptide gene expression revealed a progressive and statistically significant increase in mRNA levels of cryptdin 1 and sPLA2 in CML mice, demonstrating Paneth cell-specific gene expression (**Figures 1G and 1H**). To prove a direct and causal relationship between the development of CML and PCM, we exploited the fact that the administration of Tet in the drinking water to mice with clinically evident CML reverses the leukemia phenotype and normalizes the WBC count in the peripheral blood (see Experimental Approach **Figure 1I**). Transplant recipients in both cohorts were maintained off Tet for the first 35 days at which point the majority develop leukocytosis (**Figure 1J**). Tet administration on day 35 to mice with leukocytosis and previously documented PCM (**Figure 1E**) normalized the WBC within three weeks, whereas animals that remained off Tet had progressive leukocytosis (**Figure 1J**). Mice that had normalized their WBC counts also had resolution of PCM, whereas animals with CML had persistent metaplasia (**Figure 1K**). Thus, there was a direct and causal link between the temporal development of CML and PCM within the colon in murine transplant recipients.

**Figure 1:**
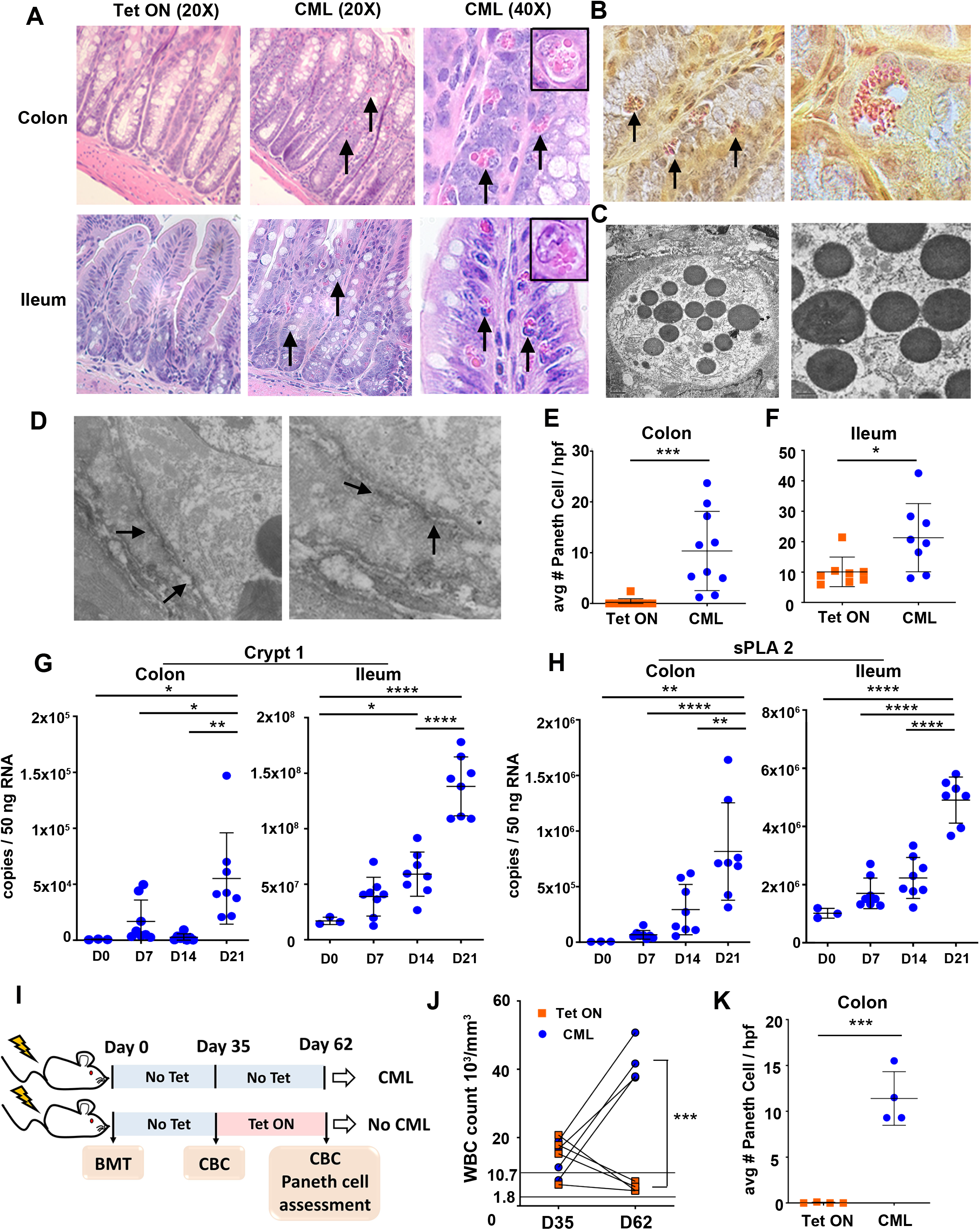
Paneth cell metaplasia develops during CML. (A-D). Lethally irradiated (1100 cGy) FVB mice were transplanted with 10 × 10^6^ BM cells from SCLtTa/bcr-abl animals, and then maintained on or off tetracycline (Tet). (A). Representative hematoxylin and eosin-stained sections of the colon and ileum from animals on or off Tet 39 days post transplantation. Original magnification is 20X and 40X. Arrows denote Paneth cells. Inset (magnification 60X) depicts individual Paneth cells in the colon and ileum. (B). Phloxine/tartrazine staining of colon from of CML animals showing characteristic phloxine staining of Paneth cell granules. Original magnification is 40X (left panel) and 60X (right panel). (C, D). Electron microscopy photomicrograph of Paneth cells in the colon of CML mice showing classical electron dense secretory granules in panel C and tight junctions (see arrows) between Paneth cell (left panel, granules visible) and adjacent cell (right panel) in panel D. Magnification is 3000X and 15,000X for panel C, and 20,000X and 35,000X for panel D. (E-G). Lethally irradiated (1100 cGy) FVB mice were transplanted with 10 × 10^6^ BM cells from SCLtTa/bcr-abl animals, and then treated with or without Tet (CML). (E, F). The average number of Paneth cells per 10 high power fields in the colon and ileum of mice 34-39 days post transplantation that were maintained on or off Tet. Results are from 2-3 experiments. (G, H). Crypt1 and sPLA2 gene expression in the colon and ileum of CML mice at specified time points post transplantation. Data are from two experiments. (I-K). Lethally irradiated FVB mice (n=4/group) were transplanted with 10 × 10^6^ BM cells from SCLtTa/bcr-abl animals and maintained off Tet for 35 days. On day 35, one cohort was placed back on Tet, while the other remained on normal drinking water. (I). Experimental scheme. (J). White blood cell counts of mice from each cohort on days 35 and 62 post transplantation. (K). The average number of Paneth cells per 10 high power fields in the colons of mice from both groups 62 days post transplantation. Results are from one of two experiments that yielded similar results. Statistics: *p<0.05, ** p<0.01, ***p<0.001, ****p<0.0001.

### IL-9 drives Paneth cell metaplasia in CML mice

To delineate a mechanistic pathway by which PCM develops in these mice, we hypothesized a role for IL-9 based on a report which had demonstrated that over expression of this cytokine in a non-disease transgenic mouse model induced PCM in the colon (20). To that end, we observed a significant increase in IL-9 gene expression in the mesenteric lymph nodes, ileum and colons of animals with CML, but not in the bone marrow or spleen (**Figure 2A**). Conversely, expression of IL-9 in non-leukemic (Tet ON) mice was essentially undetectable in all these tissue sites. IL-9 receptor mRNA levels were also augmented in the ileum and colon, but significantly reduced in the spleen of CML animals likely due to the expansion of non-IL-9R-expressing granulocyte populations (**Figure 2B**). Administration of an anti-IL-9 antibody to CML mice resulted in a significant reduction in the absolute number of Paneth cells in the colon, but not the ileum, indicating that PCM was regulated by IL-9 (**Figures 2C and 2D**). Anti-IL-9 antibody-treated mice also had a corresponding decrease in mRNA levels of cryptdin 1 and sPLA2 in the colon and ileum when compared to CML mice that received an isotype control antibody, as well as animals maintained on Tet (**Figures 2E and 2F**). Thus, blockade of IL-9 signaling effectively abrogated the development of PCM in the colon and reduced gene expression of Paneth cell AMPs in both the ileum and colon of leukemic animals.

**Figure 2:**
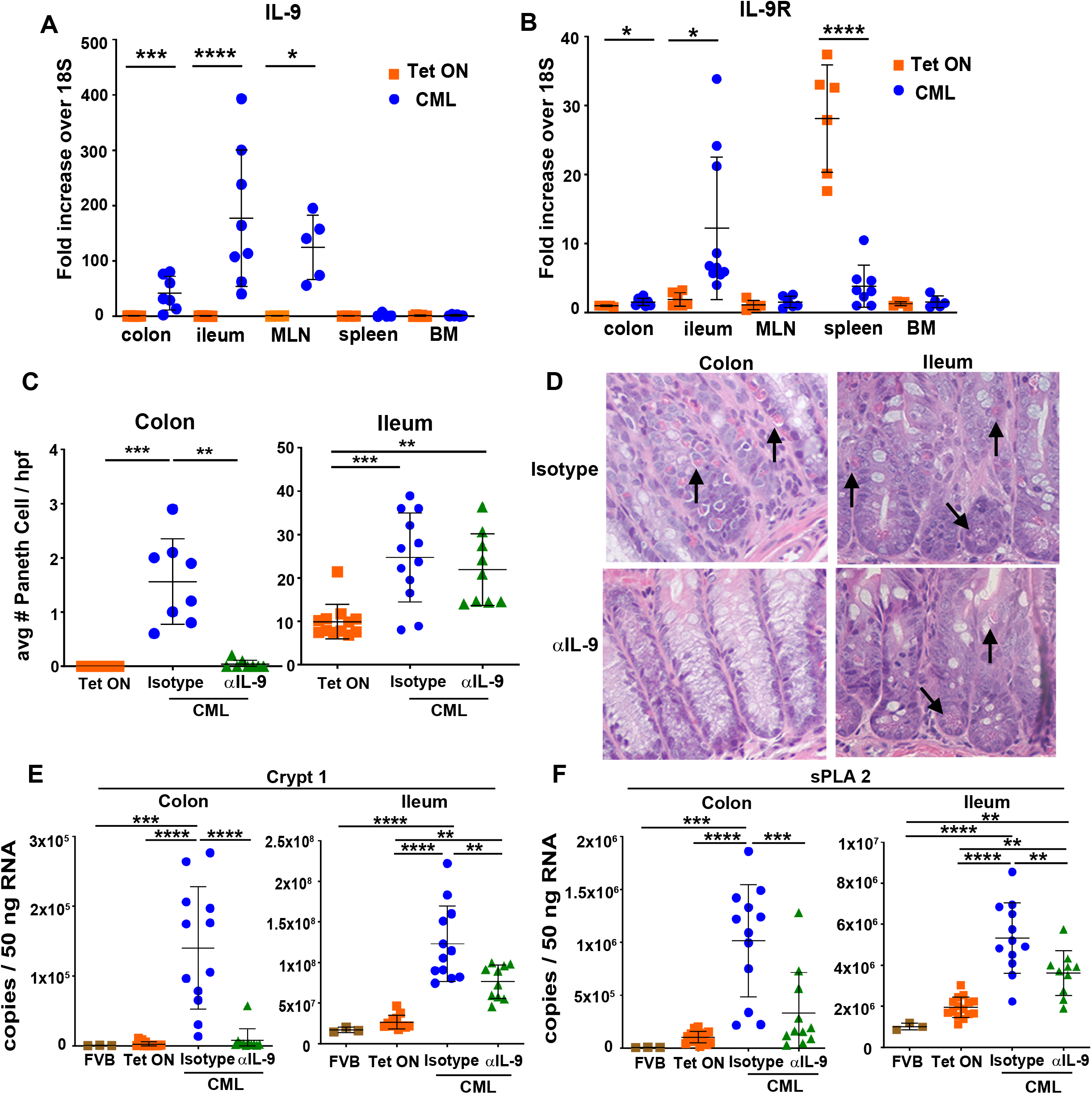
Paneth cell metaplasia is IL-9 dependent. (A-F). Lethally irradiated (1100 cGy) FVB mice were transplanted with 10 × 10^6^ BM cells from SCLtTa/bcr-abl animals. (A, B). mRNA expression of IL-9 (panel A) and IL-9 receptor (panel B) in the colon, ileum, mesenteric lymph nodes, bone marrow and spleen of animals 21-22 days post transplantation that were maintained on or off Tet (CML) is depicted. Data are expressed as fold increase over 18S. Data are from 2-4 experiments per panel. (C). The average number of Paneth cells per 10 high power fields in the colon and ileum of mice that were maintained on or off Tet and then treated three times per week with an isotype control or anti-IL-9 antibody is shown. Analysis was performed 21-39 days post transplantation. Data are from 2-3 experiments per panel. (D). Representative hematoxylin and eosin-stained sections of the colon and ileum from animals that were treated with an isotype control or anti-IL-9 antibody as in panel C. Original magnification is 40X. (E,F). Crypt1 and sPLA2 gene expression presented as copies/50 ng RNA in the colon and ileum of animals that were maintained on or off Tet and then treated with an isotype control or anti-IL-9 antibody three times per week. Normal nontransplanted FVB animals served as an additional control. Analysis was performed 21-22 days post transplantation. Data are from 2-3 experiments per panel. Statistics: *p<0.05, ** p<0.01, ***p<0.001, ****p<0.0001.

### Metabolic profiling identifies increased histamine pathway metabolites in the GI tract of CML mice

The unexpected emergence of PCM in the colon and Paneth cell hyperplasia in the ileum of CML mice led us to examine whether other cellular and metabolic alterations occurred in the GI tract of these animals. To address this question, we performed unbiased metabolic profiling and examined 973 biochemical metabolites in ileal tissue from leukemic (CML) and non-leukemic FVB recipients of syngeneic marrow grafts (FVB SYN). A total of 676 metabolites were differentially expressed (p<0.05 cutoff) between these two cohorts with 292 metabolites over expressed and 384 under expressed in CML mice (**Figure 3A**). Principal component analysis confirmed that CML mice differed substantially from non-leukemic mice with over 50% of the observed variance demonstrable along PCA1 (**Figure 3B**). Hierarchical clustering analysis demonstrated that biochemical differences were most prominently observed in amino acid and phospholipid metabolism (**Figure 3C**). A more discriminatory and statistically rigorous metabolic analysis revealed that there were 59 overexpressed and 78 under expressed metabolites between these two cohorts based on defined cutoff criteria (|log_2_(fold change) | >1.0 and p_adjusted_<0.0001 (**Figure 3D**), full list available in **Supplemental Table 1**). Examination of the top twenty most over expressed metabolites in CML mice revealed that three of the top 10 were involved in the metabolism of the aromatic amino acid histidine. Specifically, histamine which is directly downstream from histidine by way of the enzymatic action of histidine decarboxylase was the most highly expressed metabolite (144-fold increase) (**Figure 3E**). Furthermore, the downstream histamine metabolites, 1-methyl-4-imidazoleacetate and 1-methylhistamine (**Figure 3F**), were also significantly higher in CML versus non leukemic syngeneic controls (**Figure 3G**), indicating that there was activation of the histamine pathway in the GI tract that occurred concurrently with the development of PCM.

**Figure 3:**
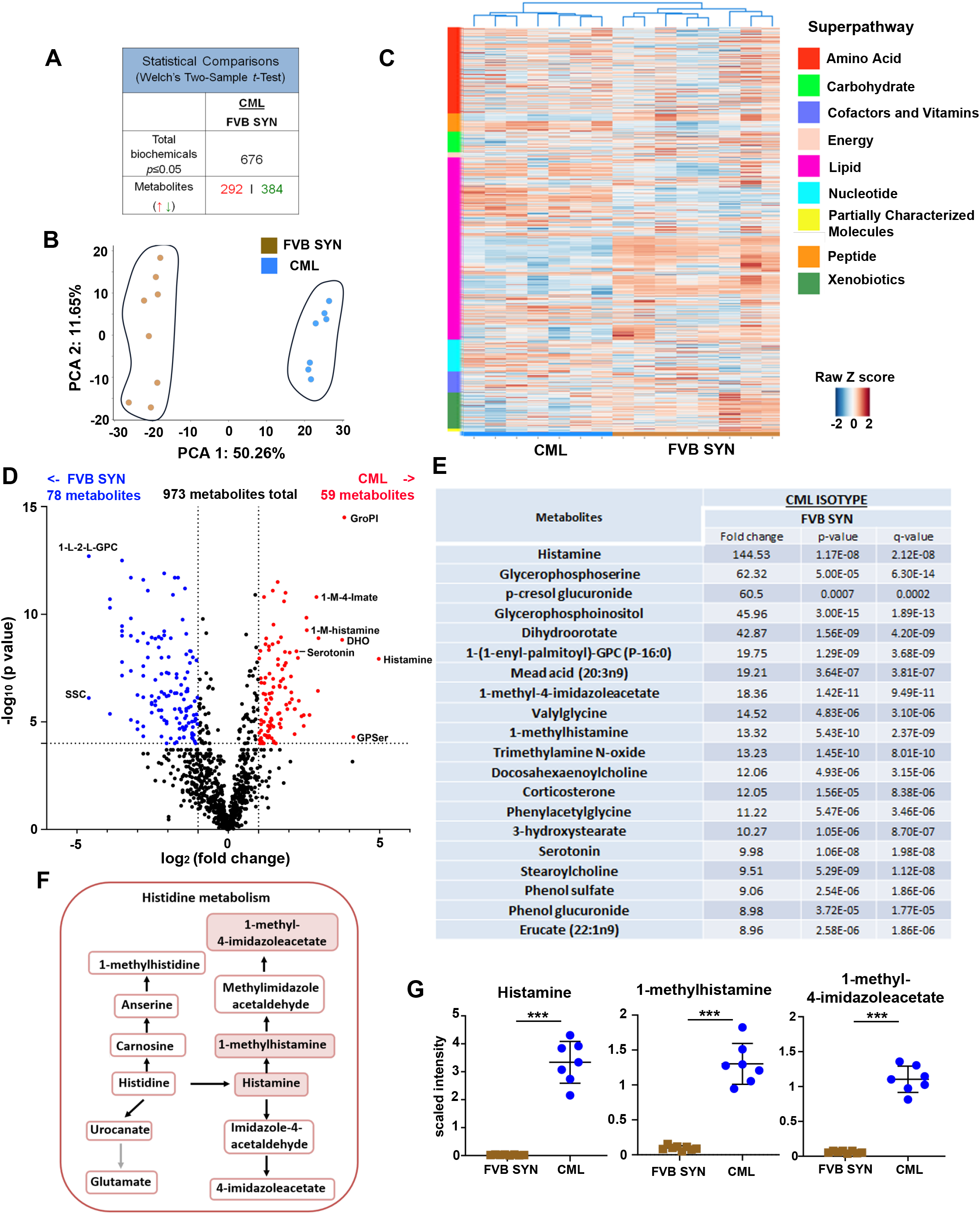
Histamine and its downstream metabolites are over expressed in the GI tract of CML mice with PCM. (A-G). Lethally irradiated FVB mice were transplanted with BM cells from normal FVB (syngeneic, FVB Syn) or SCLtTa/bcr-abl (CML) donors. Ileal tissue was harvested from animals 35 days post transplantation and processed for metabolite analysis (see Material and Methods). (A). Total number of metabolites that significantly differed (p<0.05) between FVB SYN and CML mice. (B). Principal component analysis of experimental groups. PC1 (x-axis) accounted for 50.3% of variability while PC2 (y-axis) accounted for 11.7%. (C). Heat map depicting z-scored expression of metabolites which segregate into specified pathways in each of the experimental groups. Each pathway is color-coded and shown in the far-left column, while experimental groups are depicted in the bottom horizontal color-coded lines. (D). Volcano plot demonstrating differentially expressed metabolites between syngeneic recipients and CML mice. Cutoff parameters were |log_2_ fold change| > 1.0 and p_adjusted_ <0.0001. (E). List of the top 20 metabolites that were over expressed in CML mice versus syngeneic recipients. Fold-change is depicted along with associated p-values and q-values. Q values denote the false discovery rate and were computed using the method of Storey J and Tibshirani R. (see reference 57). Abbreviations: GroPI, glycerophosphoinositol; GPSer, glycerophosphoserine; DHO, Dihydroorotate; SSC, Cysteine s-sulfate; 1-L-2-L-GPC, 1-linoleoyl-22-linoleoyl-GPC. (F). Graphical representation of the histidine metabolic pathway. (G). Scaled intensity of histamine, 1-methylhistamine, and 1-methyl-4-imidazoleacetate in each experimental group. Statistics: ***p<0.001.

### CML induces IL-9 regulated mast cell hyperplasia, intestinal remodeling, and hyperplasia of specialized intestinal epithelial cells

Histamine is found solely in basophils and mast cells in the periphery; however, only mast cells are tissue resident (21). The histamine pathway metabolite differences that we identified in CML mice therefore prompted us to examine whether mast cells were increased in the GI tract of these animals, and if their presence was similarly regulated by IL-9. This premise was supported by the observation that ileal tissue from CML animals had a ten-fold increase in serotonin which is also produced by mast cells (**Figures 3E and 4A**) (22). Gene expression of the IgE high affinity receptor (*Fcer1a*) which is present on all mast cell populations was significantly increased in the ileum of CML mice and correspondingly reduced in anti-IL-9 antibody-treated animals (**Figure 4B**). Mast cells are present in two different locations (i.e., connective tissue and the epithelial layer) which can be distinguished by specific mast cell proteases (23). To that end, we found that expression of mMCP1,2,4, that is in both mucosal and connective tissue sites, was significantly increased in leukemic animals but reduced in anti-IL-9 antibody-treated mice (**Figure 4C**), whereas there was no difference in the expression of mMCP-7 which is located only in connective tissue mast cells (**Figure 4D**). Isolation of intestinal epithelial (IECs) and lamina propria cells from the colon revealed that mMCP 1,2,4 was predominantly expressed in IECs, confirming that mast cells were confined to the mucosa (**Figures 4E**). Immunohistochemistry staining with mMCP 1, which is mucosally restricted, also demonstrated mast cell localization in the epithelial layer of the ileum in CML mice (**Figure 4F**).

**Figure 4:**
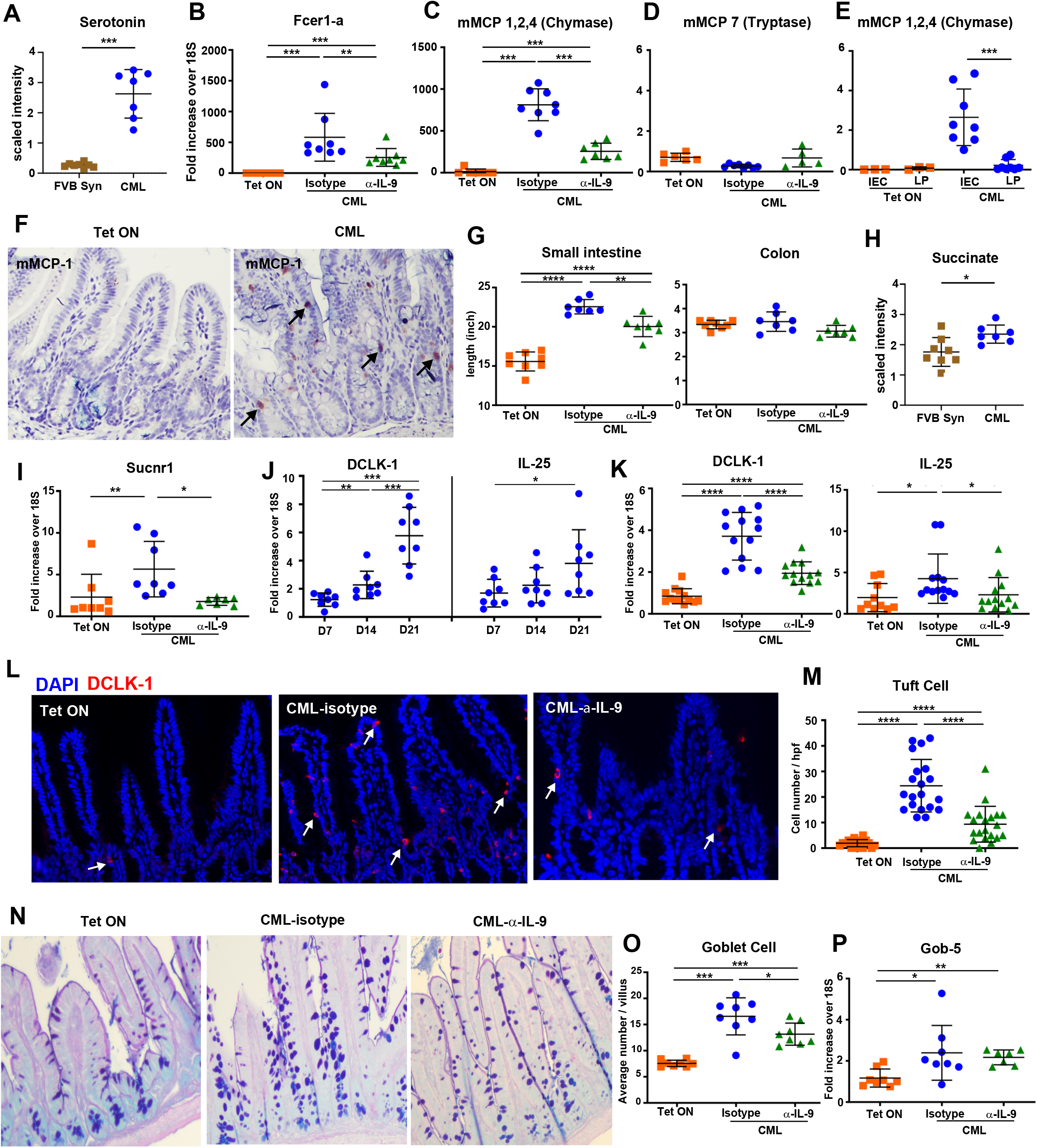
CML induces intestinal remodeling and hyperplasia of specialized epithelial cell populations. (A). Scaled intensity of serotonin in the ileum of irradiated FVB recipients transplanted with BM cells from normal FVB (FVB Syn) or SCLtTa/bcr-abl (CML) donors. (B-D). Lethally irradiated FVB mice were transplanted with 10 × 10^6^ BM cells from SCLtTa/bcr-abl mice and maintained on Tet or taken off Tet and then treated with an isotype control or anti-IL-9 antibody three times per week. mRNA expression of Fcer1-a (panel B), mMCP 1,2,4 (panel C), and mMCP 7 (panel D) in the ileum 21-22 days post transplantation. (E). mRNA expression of mMCP 1,2,4 from intestinal epithelial cells (IECs) and lamina propria cells (LP) in the colon of mice transplanted with SCLtTa/bcr-abl BM cells and maintained on or off Tet 35 days post transplantation. (F). Representative photomicrograph of mMCP 1,2,4 staining in the epithelial cell layer of the ileum 21-22 days post transplantation. (G). Small intestines and colon length of mice 21 days post transplantation that were maintained on or taken off Tet and then treated with an isotype control or anti-IL-9 antibody. (H). Scaled intensity of succinate in the ileum of recipients transplanted with BM from syngeneic or SCLtTa/bcr-abl donors. (I). mRNA expression of the succinate receptor (Sucnr1) 21-22 days post transplantation in animals that were either maintained on or taken off Tet and then treated with an isotype control or anti-IL-9 antibody. (J). mRNA expression of DCLK-1 and IL-25 in the ileum of mice who were transplanted with SCLtTa/bcr-abl BM at specified time points. (K-P). Lethally irradiated FVB mice were transplanted with BM from SCLtTa/bcr-abl mice and then maintained on or taken off Tet and then treated with an isotype control or anti-IL-9 antibody. DLCK-1 and IL-25 gene expression is shown (panel K). Representative immunofluorescence image of DLCK-1^+^ cells (panel L) and absolute number of DLCK-1^+^ tuft cells (panel M) in the ileum is depicted. Representative Alcian blue staining of goblet cells (panel N), absolute number of goblet cells (panel O), and mRNA expression of Gob-5 in the ileum (panel P) are shown. Analysis was performed 21-22 days post transplantation. Results are from two-three experiments in all panels. Statistics: *p<0.05, ** p<0.01, ***p<0.001, ****p<0.0001.

In association with mast cell alterations, we observed that there was CML-induced intestinal remodeling as evidenced by a marked increase in small intestinal, but not colonic, length that was partially attenuated by IL-9 signaling blockade (**Figure 4G**). This phenomenon has been observed in helminth infections where tuft cells respond to succinate and drive intestinal remodeling in infected hosts (23). In that regard, metabolic profiling studies revealed that there were increased succinate levels (**Figure 4H**), as well as augmented gene expression of the succinate receptor (sucnr1) in the ileum of CML mice (**Figure 4I**). Furthermore, expression of the Tuft cell marker DCLK-1 (double cortin-like kinase-1) and IL-25, which is secreted by Tuft cells and regulates type 2 immune responses (24,25), increased in a time-dependent manner (**Figure 4J**) that coincided with the temporal kinetics of leukemia progression (**Supplemental Figure 1**). Gene expression of DCKL-1 and IL-25 in the ileum of CML mice (**Figure 4K**) as well as the absolute number of DLCK-1^+^ tuft cells were also significantly reduced by IL-9 signaling blockade (**Figures 4L and 4M**). Concurrent with tuft cell hyperplasia, we observed that CML mice developed goblet cell hyperplasia that was significantly reduced in anti-IL-9 antibody-treated animals (**Figures 4N and 4O**). Expression of the goblet cell-specific gene, Gob-5, which is a calcium activated chloride gene involved in mucin production (26), was augmented in leukemic animals, although expression was not affected by IL-9 signaling blockade (**Figure 4P**). Collectively, these studies demonstrated that CML induced extensive intestinal remodeling characterized by the IL-9 regulated expansion of mast, tuft and goblet cells.

### IL-13 regulates tuft and goblet cell hyperplasia, but not PCM during CML

Binding of succinate to the succinate receptor on tuft cells leads to the secretion of IL-25 which activates ILC2s leading to release of IL-13 (27). IL-13 is then able to promote renewal of intestinal stem cells and their differentiation into tuft and goblet cells resulting in a feed forward loop (28). The observation that succinate was increased in the ileum of mice with CML therefore led us to examine whether IL-13, in addition to IL-9, co-regulated the expansion of specialized intestinal epithelial cell populations. We observed that IL-13 mRNA levels were significantly increased in the ileum, colon, mesenteric lymph nodes, and bone marrow of leukemic animals with differential expression in the ileum being the most pronounced (**Figure 5A**). In fact, the gene expression profile of IL-13 was similar to that of IL-9 in CML versus non-leukemic mice with the exception of the bone marrow (**Figure 2A**). Administration of an anti-IL-13 antibody that had previously been shown to effectively inhibit allergic disease in mice (29) had no effect on PCM or hyperplasia in either the colon or ileum, respectively (**Figure 5B**). There was, however, a significant reduction in crypt 1 (**Figure 5C**) but not sPLA2 (**Figure 5D**) gene expression in the ileum and colon of animals that received anti-IL-13 antibody. Conversely, blockade of IL-13 signaling had no effect on gene expression of the mast cell markers *Fcer1-a*, mMCP 1,2,4 or mMCP 7 (**Figures 5E-5G**), indicating that IL-9, and not IL-13, was the primary regulator of mast cell gene expression. Administration of anti-IL-13 antibody, however, did result in a significant reduction in the expression of DCLK-1 and IL-25 in tuft cells (**Figure 5H**) and Gob-5 in goblet cells (**Figure 5I**). Thus, these studies demonstrated that IL-13 regulated Paneth cell-specific gene expression along with tuft cell and goblet cell hyperplasia, but had no effect on the development of Paneth cell metaplasia or hyperplasia during CML.

**Figure 5:**
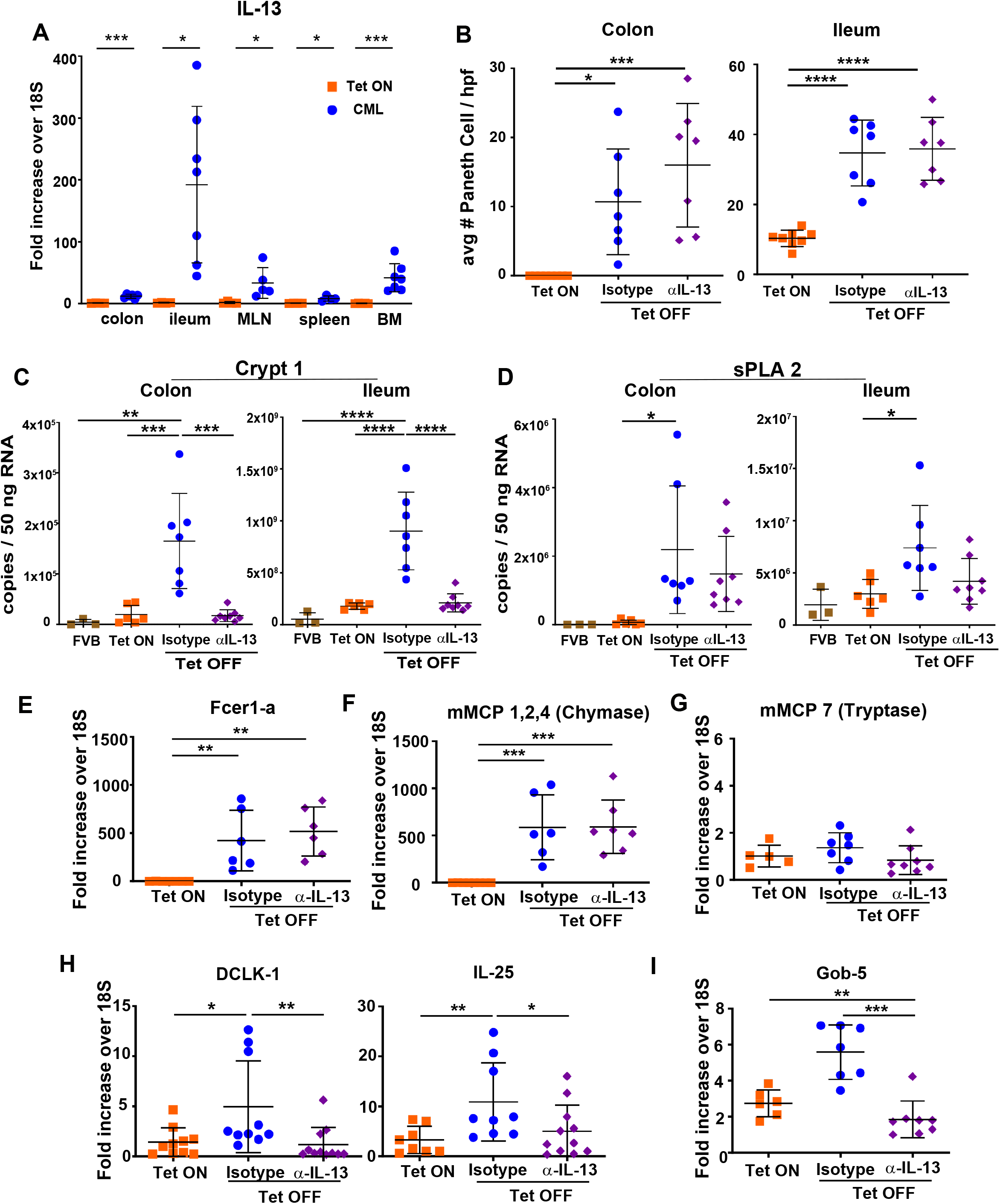
IL-13 regulates tuft and goblet cell hyperplasia but not Paneth cell metaplasia. (A). Lethally irradiated FVB mice were transplanted with BM cells from SCLtTa/bcr-abl animals. mRNA gene expression of IL-13 in the colon, ileum, mesenteric lymph nodes, spleen and bone marrow of animals that were maintained on or off Tet 21-22 days post transplantation. (B-I). Lethally irradiated FVB mice were transplanted with BM from SCLtTa/bcr-abl mice. Animals were either maintained on or taken off Tet and then treated three times per week with an isotype control or anti-IL-13 antibody. (B-D). The average number of Paneth cells per 10 high power fields (panel B), and mRNA expression of Crypt1 (panel C) and sPLA2 gene expression (panel D) in the colon and ileum 34-35 days post transplantation. (E-G). mRNA expression of Fcer1-a (panel E), mMCP 1,2,4 (panel F) and mMCP 7 (panel G) in the ileum is depicted. (H, I). mRNA expression of DCLK-1 and IL-25 (panel H) and Gob-5 (panel I) in the ileum of mice. Analysis was performed 34-35 days post transplantation. Data are from two experiments per panel. Statistics: *p<0.05, ** p<0.01, ***p<0.001, ****p<0.0001.

### Type 2 innate lymphoid cells (ILC2s) are the primary producers of IL-9

To define the cellular source of IL-9, we considered that previous studies had shown IL-9 is produced by a variety of cell types, including type 2 helper T cells (30,31), T_H_17 cells (32,33), CD4^+^ regulatory T cells (34), innate lymphoid cells (35), and mast cells (31). We therefore conducted an iterative approach in which we examined each of these cell populations to determine whether IL-9 production was detectable in these immune subsets. With respect to mast cells, while we had previously shown that mMCP 1,2,4 was predominantly expressed in the epithelial layer within the colon (**Figures 4E**), expression of IL-9 was confined to the lamina propria of CML mice (**Figure 6A**). In addition, immunofluorescence staining of colonic tissue revealed that there was no colocalization of mMCP-1 and IL-9 (**Figure 6B**), indicating that mast cells were not the source of this cytokine within the GI tract. To determine whether T cells produced IL-9, we first examined the mLN since T cells are a prominent cellular component of this tissue and prior studies had shown increased expression of IL-9 in this tissue site (**Figure 2A**). mLN cells were flow sorted into three populations (i.e., CD4^−^ TCRαβ^−^, CD4^+^ TCRαβ^+^ and CD4^−^ TCRαβ^+^ cells) and q-PCR was employed to demonstrate that IL-9 expression was significantly increased in CD4^−^ TCRαβ^−^ cells when compared to CD4^+^ TCRαβ^+^ and CD4^−^ TCRαβ^+^ T cells (**Figure 6C**). Furthermore, immunofluorescence staining demonstrated a lack of colocalization of CD3 and IL-9 in both the ileum and colon (**Figure 6D**), indicating that T cells were not the source of IL-9. As additional confirmation, we treated CML mice with an anti-CD4 antibody that depletes these T cells from tissue sites (**Supplemental Figure 2A**). We observed no difference in IL-9 expression in the ileum and an increase in IL-9 expression in the colons of anti-CD4 antibody-treated mice (**Figure 6E**). Moreover, there was no decrease in the absolute number of Paneth cells (**Supplemental Figure 2B**) or the expression of the Paneth cell genes, crypt 1 and sPLA 2, in the colon or ileum (**Supplemental Figures 2C and 2D**). In addition, there was no difference in colonic or small intestinal length (**Supplemental Figures 2E**), or expression of IL-25, DCLK-1 and mMCP 1,2,4 in the ileum (**Supplemental Figures 2F-2H**). In contrast, nearly all IL-9-expressing cells in the ileum and colon co-expressed GATA3 (**Figure 6F**), and the majority also had colocalized expression of KLGR1 (**Figure 6G**), that are markers for ILC2s (36). There was also an increased number of ILC2s in the ileum and near significance in the colon (**Figure 6H**). Furthermore, we observed increased expression of ST2 (*Il1rl1*) and IL-17RB (**Figures 6I and 6J**), as well as IL-4, IL-5 and amphiregulin (Areg) which are cytokines produced by ILC2s (37) (**Figures 6K-6M**). Collectively, these data identified type 2 innate lymphoid cells (ILC2s) as the primary source of IL-9 within the ileum and colon of CML mice.

**Figure 6:**
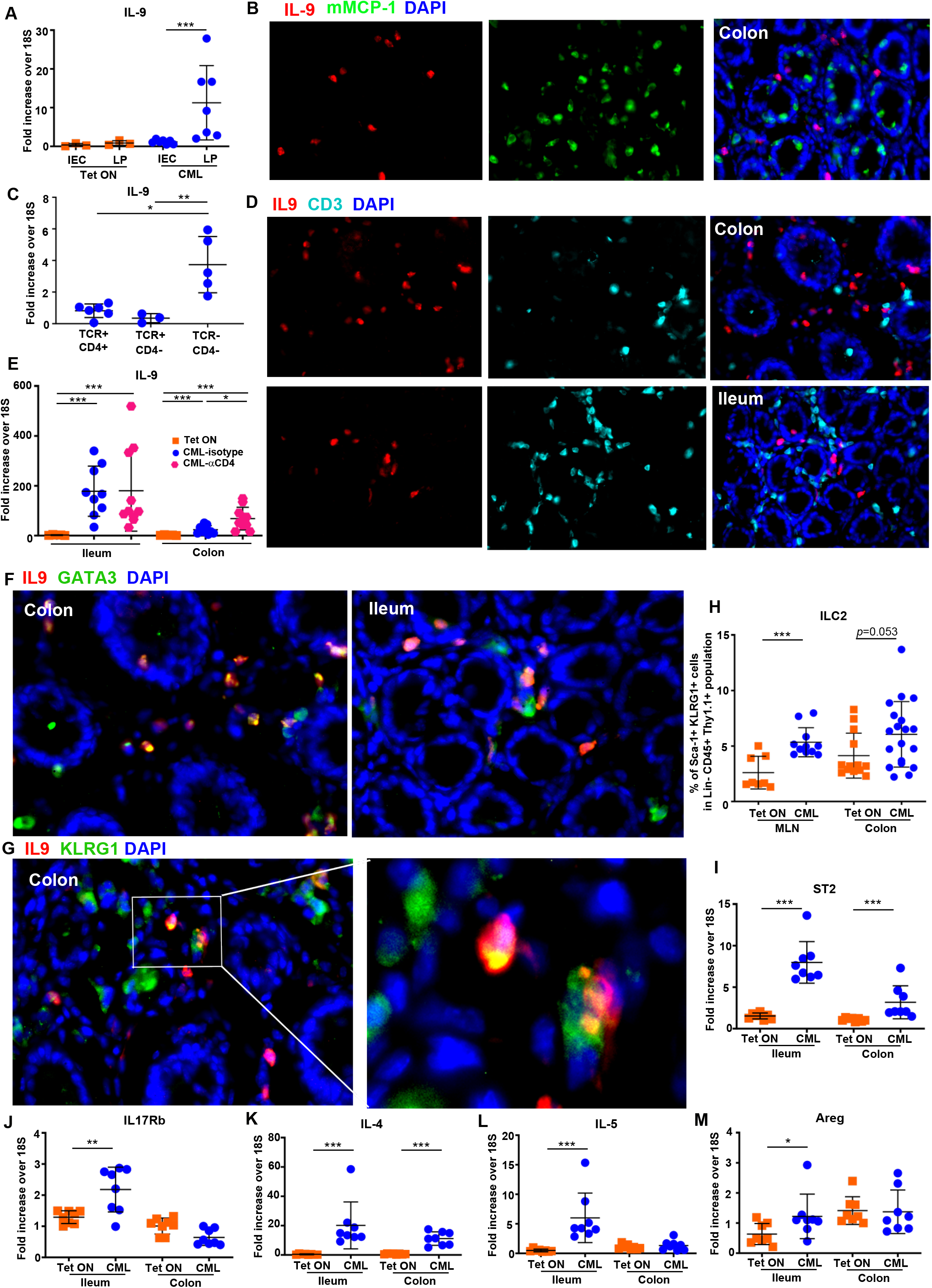
Type 2 ILCs are the primary source of IL-9. (A). Lethally irradiated FVB mice were transplanted with BM cells from SCLtTa/bcr-abl animals. Animals were then maintained on or off Tet. mRNA expression of IL-9 from isolated intestinal epithelial (IECs) and lamina propria cells (LP) 28-35 days post transplantation. (B). Immunofluorescence staining of the colon showing isolated and merged expression of mMCP-1 and IL-9 in CML mice. (C). IL-9 gene expression in sorted TCR^+^ CD4^+^, TCR^+^ CD4^−^ and TCR^−^ CD4^−^ populations from the mesenteric lymph nodes of CML mice 4-5 weeks post transplantation. (D). Immunofluorescence staining of the ileum and colon showing isolated and merged expression of CD3 and IL-9. (E). Lethally irradiated FVB mice were transplanted with BM from SCLtTa/bcr-abl mice. Animals were either maintained on or taken off Tet and then treated with an isotype control or anti-CD4 antibody twice weekly for 4 weeks. IL-9 gene expression in the ileum and colon. Lethally irradiated FVB mice were transplanted with BM cells from SCLtTa/bcr-abl animals and maintained off Tet. (F). Immunofluorescence staining in the ileum and colon showing isolated and merged expression of GATA3 and IL-9 in CML mice. (G). Immunofluorescence staining in the colon showing isolated and merged expression of KLRG1 and IL-9. (H). Frequency of ILC2s in the mLN and colon 4 weeks post transplantation. (I-M). mRNA expression of ST2 (*Il1rl1*), IL-17RB, IL-4, IL-5, and amphiregulin (Areg) in the ileum and colon 21 days post transplantation of FVB mice transplanted with SCLtTa/bcr-abl BM and then maintained on or off Tet. Statistics: *p≤0.05, ** p<0.01, ***p<0.001.

### IL-33 regulates the expression of ILC2-derived cytokines in the GI tract

We then sought to define upstream events that induced the ILC2-mediated production of IL-9 in the GI tract of CML mice. We observed that CML induced an inflammatory environment in the GI tract characterized by increased expression of IL-6, IFN-γ, TNF-α, IL-22 and GM-CSF in the ileum and colon (**Figure 7A**). Furthermore, there were augmented levels of inflammatory lipids; specifically, eicosanoids (5-HETE, 12-HETE), glycerophospholipids (glycerophosphoserine, glycerophosphoinositol) (38) and mead acid (39) in the ileum of leukemic animals (**Supplemental Table 1, Figures 3B and 7B**). Prior studies have shown that IL-33, IL-25, and/or thymic stromal lymphopoietin (TSLP) are major activators of ILC2s (40), and these cytokines function as alarmins that are produced in response to cellular injury and inflammation (41). To that end, we noted that mRNA expression of IL-33 was significantly increased in the ileum and colon (**Figure 7C**). Conversely, IL-25 which was previously shown to be augmented in the ileum where there was tuft cell hyperplasia (**Figure 4K**), was not increased in the colon (**Supplemental Figure 3A**) and TSLP was decreased in both tissue sites (**Supplemental Figure 3B**). IL-33 is localized in the nucleus prior to its secretion as a 266 amino acid protein (42) that is subsequently cleaved by proteases from neutrophils and mast cells into smaller mature forms that have increased potency (43,44). Western blot analysis confirmed increased expression of IL-33 in the ileum and colon of CML animals which was attributable to the more potent shorter 15 kDA cleavage protein (**Figures 7D and 7E, Supplemental Figures 3C, 3E, 3F and 3H**). We also observed that protein expression of chymase was significantly increased in the ileum and colon of CML animals (**Figures 7F and 7G, Supplemental Figures 3D, 3E, 3G and 3H**), providing a potential explanation for the observed smaller IL-33 cleavage product (44). IL-33 localized to the intestinal epithelial cell layer, as demonstrated by both western blot (**Figures 7H and 7I**) and immunofluorescence (**Supplemental Figures 3I and 3J**), and the predominant form in the absence of extracellular exposure to chymase was the 33kDa product. Blockade of IL-9 signaling abrogated the expression of chymase in the ileum and colon (**Figures 7J and 7K, Supplemental Figures 3L, 3M and 3O**) and was associated with a corresponding reduction in the expression of the short form of IL-33 (**Figures 7L and 7M, Supplemental Figures 3K, 3M, and 3N**), further supporting the premise that mast cell-derived chymase could mediate proteolytic cleavage of IL-33. To determine if IL-33 regulated the expression of IL-9 and other ILC2-derived cytokines, CML mice were treated with either an anti-ST2 or isotype control antibody. Inhibition of IL-33 signaling resulted in a significant decrease in gene expression of IL-9 (**Figure 7N**), along with IL-4, IL-5 and IL-13 (**Figure 7O**), but not mMCP or IL-25 (**Figure 7P**) in the ileum of leukemic animals, demonstrating that IL-33 regulated the expression of ILC2-derived cytokines.

**Figure 7:**
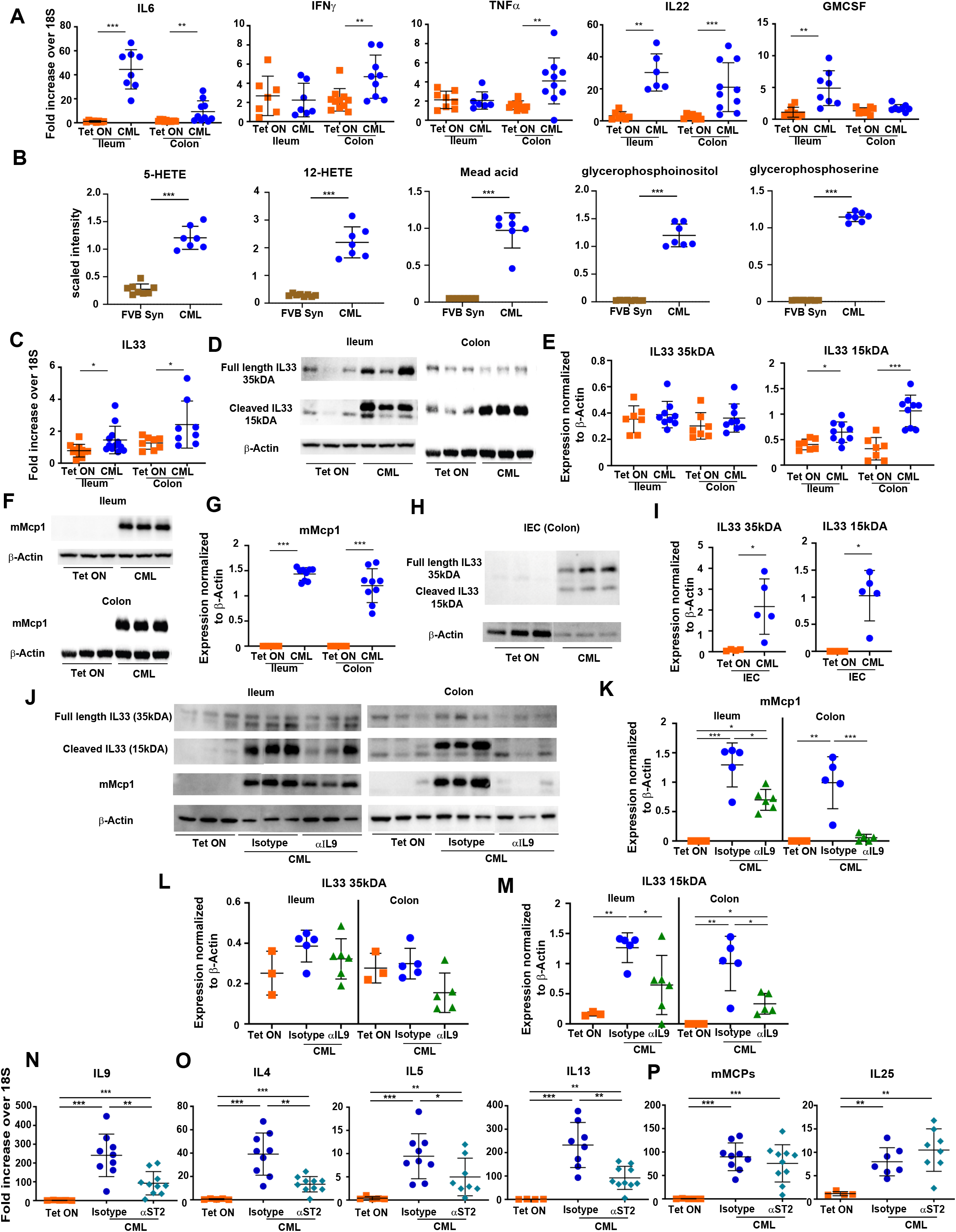
Epithelial-derived IL-33 drives IL-9 production by type 2 ILCs. (A). Lethally irradiated FVB mice were transplanted with SCLtTa/bcr-abl BM and maintained on Tet or taken off Tet. IL-6, IFN-γ, TNF-α, IL-22, and GM-CSF mRNA expression in the colon of animals 21 days post transplantation. (B). Scaled intensity of 5-HETE, 12-HETE, mead acid, glycerophosphoserine and glycerophosphoinositol in the ileum of CML versus FVB SYN mice. (C). mRNA gene expression of IL-33 in the colon and ileum of CML versus FVB SYN animals. (D, E). Immunoblot of IL-33 in the ileum and colon of CML mice demonstrating long (35 kDa) and short (15 kDa) forms (panel D) and summary data (panel E). (F, G). Immunoblot of mMCP-1 in the ileum and colon of CML animals (panel F) and summary data (panel G). (H, I). Immunoblot of IL-33 in intestinal epithelial cells (IECs) in the ileum and colon of CML mice (panel H) and summary data (panel I). (J-M). Lethally irradiated FVB mice were transplanted with 10 × 10^6^ BM cells from SCLtTa/bcr-abl mice and then treated with an isotype control or anti-IL-9 antibody three times per week. Immunoblot of mMCP-1 (chymase) and IL-33 (35 kDa and 15 kDa) in the ileum and colon (panel J), and summary data (panels L-M). (N-P). Lethally irradiated FVB mice were transplanted with SCLtTa/bcr-abl BM and treated with an isotype control or anti-ST2 antibody for two weeks beginning on day 14 post transplantation. mRNA gene expression of IL-9 (panel N), IL-4, IL-5, IL-13 (panel O), mMCP 1,2,4 and IL-25 (panel P) in the ileum. Statistics: *p≤0.05, ** p<0.01, ***p<0.001.

## DISCUSSION

Paneth cell metaplasia (PCM) invariably occurs in diseases of the GI tract, such as inflammatory bowel disease and colorectal cancer (12–15). In addition, parasitic infections within the intestines have been shown to induce metaplasia (45,46); however, extra intestinal disorders have not been associated with this phenomenon. More importantly, the mechanistic pathway by which PCM develops in the GI tract has not been clearly delineated in any disease state. In the current study, we made the unexpected observation that CML, which is a hematological malignancy that is largely confined to the bone marrow, peripheral blood, and spleen, induced the development of PCM within the colon. To validate a causative relationship between CML and PCM, we employed a tetracycline-inducible mouse model that faithfully recapitulates the clinical manifestations associated with CML where mice develop leukocytosis, granulocytic hyperplasia, splenomegaly, and bcr-abl oncogene expression, all of which are hallmarks of this disease in humans (19). We then exploited the fact that administration or withdrawal of tetracycline allowed us to temporally regulate bcr-abl oncogene expression and thereby leukemia onset and progression in a very precise manner. Consequently, we were able to demonstrate that the onset of PCM was coincident with the development of CML, and conversely, cessation of bcr-abl oncogene expression in hematopoietic stem cells by reinstitution of tetracycline resulted in the concurrent resolution of leukocytosis and metaplasia. Thus, these data provided a definitive and causal link between the temporal development of CML and PCM within the colon. In addition to PCM, we observed extensive intestinal remodeling characterized by small intestinal lengthening along with tuft cell and goblet cell hyperplasia. Intestinal remodeling is a recognized sequela of helminth and protist infections which trigger a succinate-dependent and independent tuft cell circuit that induces proliferation of specialized epithelial populations (i.e., tuft and goblet cells) and the induction of a type 2 immune response within the small intestines (24,27). Activation of ILC2s is a critical component of this immune circuit since these cells express the IL-25R which makes them responsive to tuft cell-dependent secretion of IL-25 (25). This tuft cell-ILC2 circuit-triggered event is thought to be necessary to maintain appropriate energy balance for the host to mount an anti-pathogen response (24). Paneth cell metaplasia, however, is not a recognized feature of intestinal remodeling, nor does IL-25 signaling have any reported role in Paneth cell biology. Thus, we sought to define a mechanistic pathway that would link these seemingly disparate events. To that end, we identified IL-9 as the critical cytokine responsible for driving both PCM and intestinal remodeling events in CML mice (**Figure 8**). IL-9 was specifically and differentially increased in the colon and ileum where PCM and Paneth cell hyperplasia were present. In addition, antibody-mediated blockade of this signaling pathway effectively inhibited the development of PCM as evidenced by a significant decrease in the absolute number of Paneth cells as well as mRNA levels of cryptdin 1 and sPLA2. Furthermore, inhibition of IL-9 also prevented the emergence of tuft and goblet cell hyperplasia, indicating that this cytokine mediated widespread small intestinal remodeling and altered the topography of specialized epithelial cell populations.

**Figure 8:**
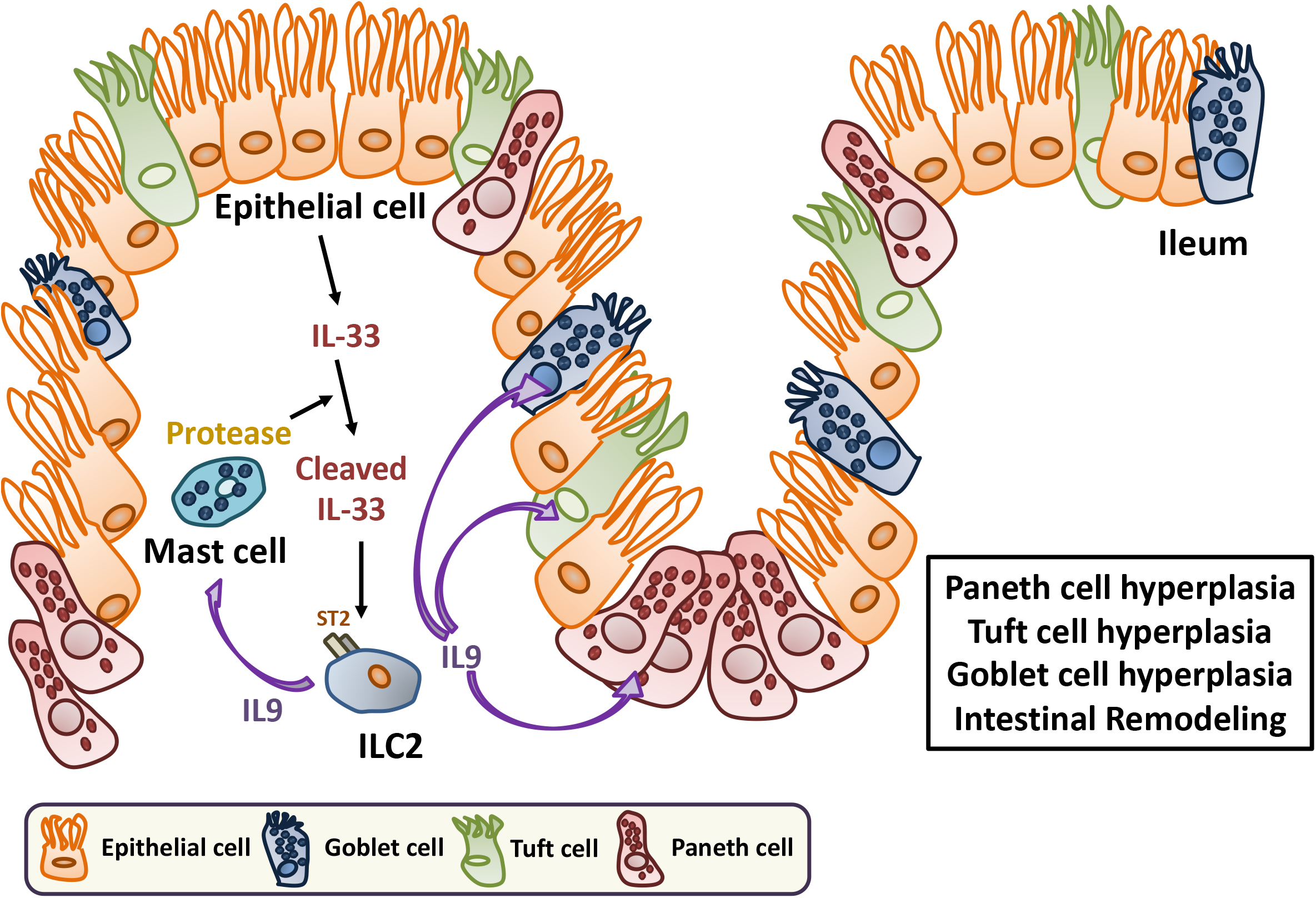
Proposed model for the development of PCM and hyperplasia of specialized epithelial cell populations by an IL-33-induced ILC2/IL-9 immune circuit. Epithelial-derived IL-33 is cleaved by mast cell protease resulting in the production of the cleaved, more potent 15 kDa form. Cleaved IL-33 binds to ST2 expressed on the surface of type 2 ILCs in the GI tract leading to the production of IL-9 which induces PCM and Paneth cell hyperplasia in the colon and ileum, respectively. IL-9 also induces hyperplasia of specialized epithelial cell populations (i.e., tuft and goblet cell hyperplasia) along with mucosal mast cell hyperplasia in the ileum.

Metabolic profiling of the small intestines also revealed significant increases in histamine and its downstream metabolites, suggesting that CML had effects on other resident intestinal cell populations that are not commonly affected during small intestinal remodeling. Mast cells are the dominant histamine-producing tissue-resident cell population and exist as two subsets; a constitutive class that is distributed in the connective tissue, and an inducible class of mucosal mast cells that are interepithelial and reside in intestinal and respiratory mucosa (23). Connective tissue mast cells in the mouse can be distinguished by expression of mMCP 4,5,6, and 7 proteases, while mucosal mast cells express mMCP 1 and 2 which function as chymases. The fact that only mMCP 1 and 2 expression was increased in the small intestines of CML animals was indicative of mucosal mast cell hyperplasia, which was further verified by immunohistochemical staining, indicating that mast cells were the likely source of histamine in leukemic animals. With respect to IL-9, this cytokine has also been shown to enhance the survival and proliferation of mast cells (31) as these cells express the IL-9 receptor on the cell surface (47). In fact, transgenic over expression of IL-9 results in intestinal mastocytosis during nematode infections (48). Notably, mast cells have also been implicated in the pathophysiology of inflammatory bowel diseases (IBD) (49,50) but have not been shown to have an etiological role in the emergence of PCM which is associated with IBD. In CML mice, we observed that mast cell hyperplasia was similarly regulated by IL-9, as anti-IL-9 antibody administration effectively abrogated hyperplasia in the ileum as confirmed by mast cell quantitation and gene expression of mast cell proteases. Thus, IL-9 coordinately regulated both PCM and mast cell hyperplasia.

The unexpected effect of CML on intestinal remodeling and epithelial cell fate decisions led us to further examine the underlying regulatory network downstream of IL-9 that helped to facilitate this remodeling event. Metabolic profiling studies revealed an increased level of succinate along with augmented expression of the succinate receptor in CML mice. The succinate receptor is known to be expressed at highest levels in tuft cells (27), and the increase in IL-25 and DLCK-1 expression along with marked small intestinal lengthening were all indicative of tuft cell hyperplasia which was confirmed by immunofluorescence staining in the small intestines. Tuft cell hyperplasia has been shown to be dependent upon the production of IL-13 by type 2 ILCs which also induces goblet cell metaplasia (51). Blockade of this pathway in CML mice resulted in a reduction in the number of tuft and goblet cells, as well as a significant decrease in Paneth cell-specific gene expression (30). However, whereas there was a reduction in crypt 1 gene expression in anti-IL-13 antibody-treated animals, there were no quantitative alterations in the absolute number of Paneth cells or other Paneth cell-specific peptides when compared to isotype antibody-treated control mice. Furthermore, IL-13 had no impact on the development of mast cell hyperplasia. Thus, IL-13, unlike IL-9, was more restricted to the regulation of tuft and goblet cell development in CML mice and had no effect on the emergence of PCM or mast cell hyperplasia.

Given the critical role of IL-9 in promoting PCM and intestinal remodeling, we sought to define the cell population that was responsible for the production of this cytokine in the GI tract. IL-9 has been shown to be produced by type 2 helper T cells (30,31), T_H_17 cells (32,33), regulatory T cells (34), and innate lymphoid cells (35), and has effects on both hematopoietic and non-hematopoietic cell populations (32). Therefore, we pursued an iterative approach to identify IL-9-expressing populations in the GI tract. Immunofluorescence staining demonstrated that there was a lack of co-localization of IL-9 and the mast cell marker mMCP-1, and IL-9 transcripts were found only in the lamina propria which was not the resident site for mucosal mast cells. In addition, IL-9 was not detectable in CD3^+^ T cells in the ileum or colon, and IL-9 transcripts were predominantly expressed in a non-T cell population in the mesenteric lymph nodes. Depletion of CD4^+^ T cells also had no effect on gene expression of IL-9, Paneth, tuft or mast cell markers, indicating that CD4^+^ T cells were not the source of IL-9. Conversely, we observed that IL-9 prominently co-localized with GATA3 and KLRG-1 which are markers for ILC2 cells (40). In addition, there was significantly increased gene expression of IL-4, IL-5, ST2, IL-17Rb, and amphiregulin (Areg) in the small intestines that are characteristic of type 2 ILCs (37). Thus, these data indicated that ILC2s were the dominant source of IL-9 in the GI tract.

To determine the upstream signaling events that led to the activation of ILC2s, we noted that CML induced an inflammatory environment within the GI tract characterized by increased expression of an array of proinflammatory cytokines (e.g., GM-CSF, TNF-α, IL-6, IFN-γ, and IL-22) as well as lipids that have been implicated in inflammation. The ability of CML to induce systemic inflammation has been reported in mice and newly diagnosed CML patients where increased levels of inflammatory cytokines and chemokines have been documented in the plasma and hematopoietic tissues (52–54). We observed that IL-33, which functions as an alarmin (42), along with its receptor ST2, were augmented in the ileum and colon of CML animals and localized to the epithelial cell layer. IL-33 in mice is secreted as a long form 266 amino acid protein that is initially localized in the nucleus of epithelial barrier tissues along with endothelial and epithelial cells in blood vessels (41). Inflammatory proteases from neutrophils (i.e., elastase and cathepsin G) (43) and mast cells (i.e., chymase and tryptase) (44) can cleave this protein into smaller mature forms of this protein which have significantly enhanced (i.e., ~10-30-fold) biological activity. In fact, we observed that there were smaller forms of IL-33 present within the GI tract along with increased protein expression of chymase which is stored as a secretory granule in mast cells. The association of mast cells in close proximity to ILC2s has been demonstrated in environmentally facing tissue sites (37). We therefore speculate that the large number of mast cells in the mucosal layer was a source of chymase leading to the enzymatic cleavage of IL-33 into more potent forms that resulted in the activation of IL-9-producing ILC2s, thereby linking mast cells to the development of PCM (**Figure 8**). This conclusion was further supported by the fact that blockade of IL-9 signaling completely abrogated expression of chymase and this was associated with disappearance of the short form of IL-33. Finally, inhibition of IL-33/ST2 signaling with an ST2 monoclonal antibody significantly reduced the expression of IL-9, along with other ILC2-derived cytokines, indicating that IL-33 was a critical upstream regulator of type 2 ILC-mediated IL-9 production.

In summary, our studies demonstrate that an extra intestinal disease can induce an inflammatory environment within the GI tract that leads to the production of IL-33 from stressed epithelial cell populations. Binding of IL-33 to ST2 induces type 2 ILC-mediated production of IL-9 which is the proximate cytokine responsible for the development of PCM as well as the emergence of extensive small intestinal remodeling characterized by hyperplasia of specialized epithelial cell populations. Thus, an IL-33-regulated ILC2/IL-9 immune circuit resident in the GI tract establishes a mechanistic pathway for the development of PCM which may also have pathophysiological relevance for the emergence of PCM arising in GI intrinsic diseases such as colorectal cancer and inflammatory bowel diseases.

## MATERIAL AND METHODS

### Mice

FVB (H-2^q^) mice were bred in the Animal Resource Center at the Medical College of Wisconsin (MCW) or purchased from Jackson Laboratories (Bar Harbor, ME). Transgenic mice in which a tetracycline-controlled transactivator was placed under the control of the murine stem cell leukemia gene 3’ enhancer [SCLtTA mice (FVB background)] were crossbred to transgenic TRE-BCR-ABL mice (FVB background) which expressed the bcr/abl oncogene to create double transgenic SCLtTA x BCR-ABL mice (i.e., CML mice) (16). Screening of SCL, bcr/abl, and double transgenic mice was conducted by PCR. Induction of bcr/abl expression by withdrawal of tetracycline (0.5 g/l) from the drinking water results in activation of the transgene and subsequent leukemogenesis, as previously described (16). All animals were housed in the American Association for Laboratory Animal Care (AALAC)-accredited Animal Resource Center of the Medical College of Wisconsin. Experiments were all carried out under protocols approved by the MCW Institutional Animal Care and Use Committee. Mice received regular mouse chow and acidified tap water ad libitum.

### Tetracycline-dependent CML transplantation model

Bone marrow (BM) was flushed from donor femurs and tibias with Dulbecco’s modified media (DMEM, Gibco-BRL, Life Technologies, Grand Island, NY) and passed through sterile mesh filters to obtain single cell suspensions. A transplant model (17) was used in this study to achieve more uniform disease onset kinetics since CML arising de novo in mice can have significant variability in disease onset (16). To conduct these studies, bone marrow was taken from CML donors and passed through sterile filters to obtain single cell suspensions. Host mice were conditioned with total body irradiation administered as a single exposure at a dose rate of 60 cGy using a Shepherd Mark I Cesium Irradiator (J.L. Shepherd and Associates, San Fernando, CA). Irradiated recipients received a single intravenous injection in the lateral tail vein of BM (10 × 10^6^) in a total volume of 0.4 ml.

### Reagents

Anti-IL-9 (MM9C1) is a previously described mouse IgG2a antibody (36) that was administered intraperitoneally at a dose of 200 μg three times per week. Mouse IgG2a (C140SF9) was used as an isotype control and administered at the same dose and schedule. Both anti-IL-9 and the isotype antibody were purified from the culture supernatant of hybridoma cells using the Proteus protein purification spin kit (Bio-Rad, Hercules, CA). Anti-IL-13 is an IgG1 antibody (55) that was administered at a dose of 10 mg/kg three times a week by intraperitoneal injection. Mouse IgG1 was used as an isotype control antibody (BioXCell, West Lebanon, NH). Anti-CD4 (clone GK1.5, rat IgG2b) (BioXCell) and rat IgG isotype antibody (Jackson ImmunoResearch, West Grove, PA) were administered at a dose of 250 μg twice per week. Anti-ST2 (R&D Systems, Minneapolis, MN, clone 245707, rat IgG2b) and rat IgG isotype antibody (Jackson ImmunoResearch) were administered at a dose of 50 or 100 μg three times per week starting at day 14 post transplantation.

### Blood collection and complete blood count (CBC)

Blood samples were obtained by venipuncture of the facial (sub-mandibular) vein to monitor the progression of CML. Complete blood counts were performed using a scil Vet ABC ™ Hematology Analyzer (Scil Animal Care Company, Gurnee, Ill).

### Phloxine-Tartrazine staining for Paneth cells

Formalin-fixed paraffin embedded (FFPE) tissue samples were cut into 5-μm sections and dewaxed with xylene and ethanol. The rehydrated sections were stained with Harris modified hematoxylin solution (Thermo, Carlsbad, CA) for visualization of nuclei in tissue sections. The sections were then immediately stained with 0.5% phloxine (Sigma, St. Louis, MO) in 0.5% CaCl2 solution, and differentiated with a saturated tartrazine (Sigma) solution. Paneth cell granules were identified under light microscopy as red-appearing granules.

### Histological analysis and quantification of specialized intestinal epithelial cells

Distal colons and small intestines were fixed in 10% neutral-buffered formalin and embedded in paraffin. Formalin-fixed paraffin embedded (FFPE) tissue samples were then cut into 5-μm sections and stained with hematoxylin and eosin. Paneth cells were quantitated by selecting 10 randomly picked high power fields and averaging the number of Paneth cells per 10 high power fields for each experimental group. The average number of Goblet cells in 10 randomly selected villi in the ileum per sample were quantitated. Tuft cells were enumerated by randomly selecting 5 high-power fields per sample with each data point representing the cell count per one high power field. All slides were coded and read in a blinded fashion. Light microscopy images were visualized with a Nikon ECLIPSE E400 microscope (Tokyo, Japan). Image acquisition was performed with a Nikon DS-Fi3 colorimetric digital camera and software package. Immunofluorescence images were visualized with a Nikon ECLIPSE Ti microscope and camera along with NES elements software package.

### Immunohistochemistry

Fresh frozen paraffin-embedded tissue samples were de-waxed with CitriSolv (DeconLabs, King of Prussia, PA) and rehydrated before performing immunohistochemistry staining using Utra-Sensitive ABC Peroxidase Staining Kit (Thermo Fisher Scientific). Samples were immersed in EDTA Antigen Retrieval Buffer (pH 8.5, Sigma) on a hot plate and simmered for 25 minutes. Endogenous peroxidase activity was quenched with Peroxidase Suppressor Buffer (Thermo Fisher Scientific) for 30 minutes at room temperature. Samples were blocked and stained with primary antibody overnight at 4°C. After fully washing with BupH Tris Buffered Saline (TBS, Thermo Fisher Scientific), samples were incubated with biotinylated secondary antibody and developed with HRP and substrate using a DAB substrate kit (Thermo Fisher Scientific). Primary and secondary antibodies used in this study are listed in Supplemental Table 2. Images were visualized with a Nikon ECLIPSE E400 microscope. Image acquisition was performed with a Nikon DS-Fi3 colorimetric camera.

### Immunofluorescence

Distal colons and small intestines were fixed in 4% formaldehyde and embedded in O.C.T. compound (Sakura, Torrance, CA) for cryosections. Tissue samples were then cut into 12-micron sections for immunofluorescence staining using a Leica CM1520 cryostat (Leica Biosystems, Deer Park, IL). Samples were first fixed in methanol in −20°C then treated with Antigen Retrieval Buffer (pH 6, Sigma). 0.3% Triton X-100 (Sigma) was used to permeabilize the sample sections. Samples were blocked and incubated with primary antibody overnight and incubated with fluorochrome-conjugated secondary antibody on the next day. Primary and secondary antibodies used in this study are listed in Supplemental Table 2. Images were visualized with a Nikon ECLIPSE TE2000-U microscope. Image acquisition was performed with NES elements software package.

### Western Blot Analysis

Tissues harvested from mice were homogenized using a homogenizer (PowerGen 125, Thermo Fisher) in RIPA buffer (Thermo Fisher Cat. 89900) with the appropriate dilution of protease and phosphatase inhibitors (Thermo Fisher Cat. A32959). The homogenized tissue was then stored at −80 °C. Protein levels were normalized using a BCA Assay. Primary antibodies used for western blot were used as follows: rat anti-IL-33-unconjugated (1:1000; R&D Systems MAB3626), rat anti-MCP1/Mcpt1 (1:1000; R&D Systems MAB5416) and mouse anti-ß-actin (1:5000; Cell Signaling; E4D9Z). The appropriate HRP-conjugated secondary antibodies were used at 1:5000 (1:5000; R&D Systems). For a detailed list of reagents, refer to Supplementary Table 2. Relative expression of proteins was assessed using BioRad chemiluminescence imaging system.

### Electron Microscopy

Paraffin-embedded colonic samples were dewaxed in 3 × 24hr changes of xylene and rehydrated through decreasing concentrations of ethanol in distilled water. After rehydration, the sample was washed in 0.1M sodium cacodylate buffer (2 × 10min) then fixed in 2% glutaraldehyde in 0.1M cacodylate buffer for one hour. After three 10 minute washes in buffer, the sample was post fixed in 1% aqueous osmium tetroxide for one hour on ice, washed in distilled water, dehydrated through graded methanol and embedded in epoxy resin (EMBed812). Ultrathin sections (60 nm) were stained with uranyl acetate and lead citrate and viewed in a JEOL2100 transmission electron microscope (Japanese Electron Optics LtD, Tokyo, Japan) and images recorded using a Gatan Ultrascan CCD camera (Pleasanton, CA).

### Splenocyte Isolation

Splenocytes were obtained by grinding the tissues though a mesh screen with a syringe plunger. Red blood cells in the cell suspension were lysed with Ammonium Chloride Tris (ACT) lysis buffer, prepared with ammonium chloride solution and Tris-HCl solution, pH 7.2. The cell suspension was subsequently filtered through a cell strainer and prepared for further analysis.

### Isolation of Intestinal Epithelial Cells and Lymphocyte Populations in the Colon

Epithelial cells were isolated from colon samples in the pre-digestion buffer using the Lamina Propria Dissociation Kit (Miltenyi Biotec, Auburn, CA) according to the manufacturer’s instructions. To isolate lymphocytes from the lamina propria, colon samples were first washed in DMEM medium with DTT and EDTA. Samples were then digested with 10μg/ml liberase TL (Roche, Basel, Switzerland) and 0.05% DNase (QIAGEN, Hilden, Germany), and processed using the gentleMACS Dissociator (Miltenyi). The resulting cell suspension was then layered on a 44%/67% Percoll gradient (Sigma).

### Flow Cytometry

Cells were re-suspended in Fluorescence Activated Cell Sorting buffer (FACS buffer, 2% FBS in PBS) and pre-stained with LIVE/DEAD Fixable Aqua (Thermo Fisher Scientific) according to the manufacturer’s instructions to exclude dead cells. Cells were then labeled with anti-mouse fluorescently conjugated antibodies as listed in Supplemental Table 2. Flow cytometry was used to identify CD4 T cells (CD4^+^ TCRβ^+^), granulocytes (Gr1^+^ CD11b^+^), and ILC2 (CD45^+^ Lin^−^ Thy1^+^ KLRG1^+^ Sca-1^+^) with surface markers. Cells were analyzed on an LSR-II or Fortessa X-20 flow cytometer with FACSDiva software (BD). Data were analyzed using FlowJo software (BD).

### RNA extraction and precipitation

Tissues harvested from mice were homogenized using a homogenizer (PowerGen 125, Thermo Fisher) in TRIzol RNA Isolation Reagent (Thermo Fisher Scientific). The homogenizer was previously cleaned with diethyl pyrocarbonate (DEPC) water to inactive RNase. Chloroform and isopropyl alcohol were added to the TRIzol tissue lysate to precipitate RNA. For isolated cells, RNA was fixed by treating with RNAlater (Thermo Fisher Scientific) at 4°C prior to processing extracted RNA. RNA was extracted using the RNeasy Mini Kit (QIAGEN).

### Quantitative real-time PCR analysis for anti-microbial peptides and cytokine genes

To evaluate the expression of mouse AMPs, an absolute quantification method using standard curves was employed. Plasmid DNA of sPLA2 and Crypt1 was prepared using the QIAprep Spin Miniprep Kit (Qiagen) and used to generate standard curves (56). RNA was isolated from homogenized colonic and ileal tissues in trizol. cDNA was then reverse-transcribed using the QIAGEN RT kit (QIAGEN) according to the manufacturer’s instructions and subjected to RT-PCR using the QIAGEN QuantiTect SYBR Green PCR Master Mix in triplicate. Samples were run in a CFX C1000 Real-time Thermal Cycler (Bio-Rad, Hercules, CA). Standard curves were generated using Bio-Rad software, and the numbers of genomes per sample were extrapolated from the Ct values. Data are presented as copy number of AMPs genomes per 50ng of mouse RNA.

To determine gene expression values of cytokines, RNA and cDNA were prepared and run as described above with the following modifications. An 18S reference gene was amplified using the QuantiTect Primer Assay Kit (Qiagen). The primers were purchased from Integrated DNA Technologies (Coralville, IA) and are listed in Supplemental Table 3. Specificity for all q-PCR reactions was verified by melting curve analysis. Cytokine values in different cohorts were normalized to an internal control 18S. To calculate fold-change in gene expression, the average ΔΔCt values from triplicate wells were combined from separate experiments.

### Metabolic Profiling

Metabolic profiling of ileal samples from CML and non-leukemic control animals was conducted by Metabolon (Morrisville, NC). Samples were prepared using the automated MicroLab STAR® (Hamilton Company, Reno, NV). Several recovery standards were added prior to the first step in the extraction process for QC purposes. To remove protein, dissociate small molecules bound to protein or trapped in the precipitated protein matrix, and to recover chemically diverse metabolites, proteins were precipitated with methanol under vigorous shaking for 2 min (Glen Mills GenoGrinder 2000, Clifton, NJ) followed by centrifugation. The resulting extract was divided into five fractions: two for analysis by two separate reverse phase (RP)/UPLC-MS/MS methods with positive ion mode electrospray ionization (ESI), one for analysis by RP/UPLC-MS/MS with negative ion mode ESI, one for analysis by HILIC/UPLC-MS/MS with negative ion mode ESI, and one sample was reserved for backup. Samples were placed briefly on a TurboVap® (Zymark, Hopkinton, MA) to remove the organic solvent. The sample extracts were stored overnight under nitrogen before preparation for analysis.

Several types of controls were analyzed in concert with the experimental samples: a pooled matrix sample generated by taking a small volume of each experimental sample (or alternatively, use of a pool of well-characterized human plasma) served as a technical replicate throughout the data set; extracted water samples served as process blanks; and a cocktail of quality control (QC) standards that were carefully chosen not to interfere with the measurement of endogenous compounds were spiked into every analyzed sample, allowed instrument performance monitoring and aided chromatographic alignment. Instrument variability was determined by calculating the median relative standard deviation (RSD) for the standards that were added to each sample prior to injection into the mass spectrometers. Overall process variability was determined by calculating the median RSD for all endogenous metabolites (i.e., non-instrument standards) present in 100% of the pooled matrix samples. Experimental samples were randomized across the platform run with QC samples spaced evenly among the injections.

All methods utilized a Waters ACQUITY ultra-performance liquid chromatography (UPLC) and a Thermo Scientific Q-Exactive high resolution/accurate mass spectrometer interfaced with a heated electrospray ionization (HESI-II) source and Orbitrap mass analyzer operated at 35,000 mass resolution. The sample extract was dried then reconstituted in solvents compatible to each of the four methods. Each reconstitution solvent contained a series of standards at fixed concentrations to ensure injection and chromatographic consistency. One aliquot was analyzed using acidic positive ion conditions, chromatographically optimized for more hydrophilic compounds. In this method, the extract was gradient eluted from a C18 column (Waters UPLC BEH C18-2.1×100 mm, 1.7 μm) using water and methanol, containing 0.05% perfluoropentanoic acid (PFPA) and 0.1% formic acid (FA). Another aliquot was also analyzed using acidic positive ion conditions; however, it was chromatographically optimized for more hydrophobic compounds. In this method, the extract was gradient eluted from the same afore mentioned C18 column using methanol, acetonitrile, water, 0.05% PFPA and 0.01% FA and was operated at an overall higher organic content. Another aliquot was analyzed using basic negative ion optimized conditions using a separate dedicated C18 column. The basic extracts were gradient eluted from the column using methanol and water, however with 6.5mM Ammonium Bicarbonate at pH 8. The fourth aliquot was analyzed via negative ionization following elution from a HILIC column (Waters UPLC BEH Amide 2.1×150 mm, 1.7 μm) using a gradient consisting of water and acetonitrile with 10mM Ammonium Formate, pH 10.8. The MS analysis alternated between MS and data-dependent MS^n^ scans using dynamic exclusion. The scan range varied slighted between methods but covered 70-1000 m/z. Raw data files are archived and extracted as described below.

Raw data was extracted, peak-identified and QC processed using Metabolon’s hardware and software. These systems are built on a web-service platform utilizing Microsoft’s .NET technologies, which run on high-performance application servers and fiber-channel storage arrays in clusters to provide active failover and load-balancing. Compounds were identified by comparison to library entries of purified standards or recurrent unknown entities. Metabolon maintains a library based on authenticated standards that contains the retention time/index (RI), mass to charge ratio (*m/z*), and chromatographic data (including MS/MS spectral data) on all molecules present in the library. Furthermore, biochemical identifications are based on three criteria: retention index within a narrow RI window of the proposed identification, accurate mass match to the library +/- 10 ppm, and the MS/MS forward and reverse scores between the experimental data and authentic standards. The MS/MS scores are based on a comparison of the ions present in the experimental spectrum to the ions present in the library spectrum. While there may be similarities between these molecules based on one of these factors, the use of all three data points can be utilized to distinguish and differentiate biochemicals. More than 3300 commercially available purified standard compounds have been acquired and registered into LIMS for analysis on all platforms for determination of their analytical characteristics. Additional mass spectral entries have been created for structurally unnamed biochemicals, which have been identified by virtue of their recurrent nature (both chromatographic and mass spectral). These compounds have the potential to be identified by future acquisition of a matching purified standard or by classical structural analysis.

A variety of curation procedures were carried out to ensure that a high-quality data set was made available for statistical analysis and data interpretation. The quality control and curation processes were designed to ensure accurate and consistent identification of true chemical entities and to remove those representing system artifacts, misassignments and background noise. Metabolon data analysts used proprietary visualization and interpretation software to confirm the consistency of peak identification among the various samples. Library matches for each compound were checked for each sample and corrected if necessary.

### Statistical analysis

The nonparametric 2-tailed Mann-Whitney U test was used to determine significant differences in pair wise group comparisons. For more than two comparisons within groups, results were compared using the Kruskal-Wallis test with a post hoc Bonferroni correction. Welch’s two sample t test was performed for pair wise comparisons of metabolites derived from the ileum of individual animals. Statistical analysis was done using Prism software (GraphPad). For comparison of weight curves, a mixed effect model was fitted to the observations with fixed day and group effects with their interaction to model the mean, and a random subject-specific intercept and first-order continuous auto-regressive correlation to model the dependence structure. Based on the fitted model the change from day 0 to each following measurement day was compared between the three treatment groups using appropriate contrasts. Holm’s method was used for multiple testing adjustment. All the analyses were performed in R version 3.4.1 (2017-06-30), using the nlme 3.1.131 package for fitting the mixed model, and multcomp 1.4.6 for multiple testing. A p value of ≤ 0.05 was deemed to be significant in all experiments.

## AUTHOR CONTRIBUTIONS

C.-Y.Y. conducted experimental design, performed animal studies, flow cytometric analysis, wrote and edited the manuscript. A.R., A.M., W.X., and M.H. performed research and analyzed data. A.S. assisted with the biostatistical analysis. C.W. performed electron microscopy. N.S. assisted with design of the study, analyzed data and edited the manuscript. W.R.D. developed the overall concept, designed experiments, supervised the study and wrote the manuscript.

## ACKNOWLEDGMENTS

This research was supported by grants from the National Institutes of Health (HL R01 064603 and HL R01 126166) (W.R.D) and GM R01 099526 (N.S.). We thank Drs. Kate Dixon and Roger Palframan from UCB Pharma, Cambridge, MA for provision of the anti-IL-13 antibody.

