## Supplemental Data for "A Type 2 Innate Lymphoid Cell-Interleukin 9 Circuit Induces Paneth Cell Metaplasia and Small Intestinal Remodeling"

### Supplemental Figure 1

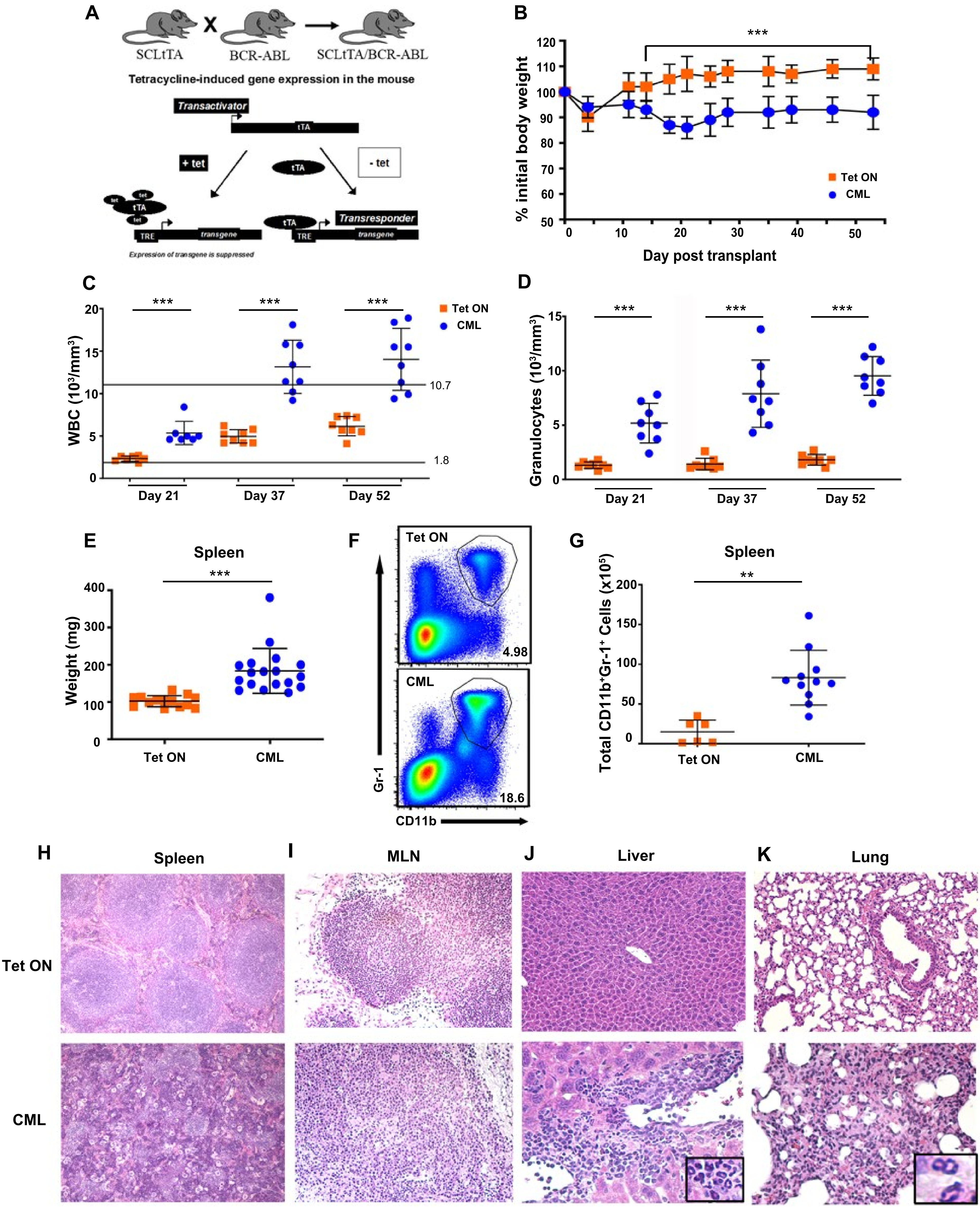

#### Supplemental Figure 2

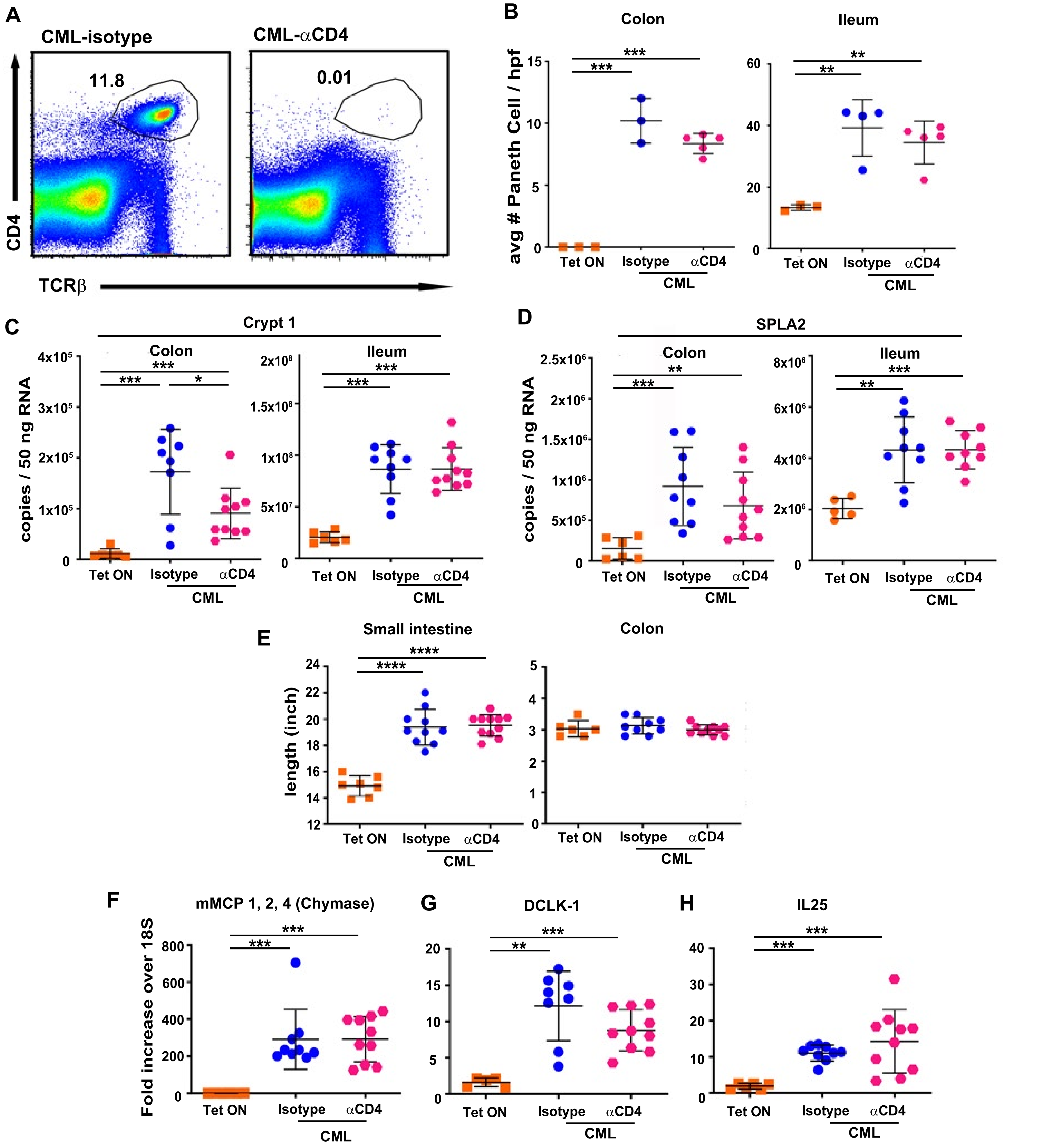

### Supplemental Figure 3

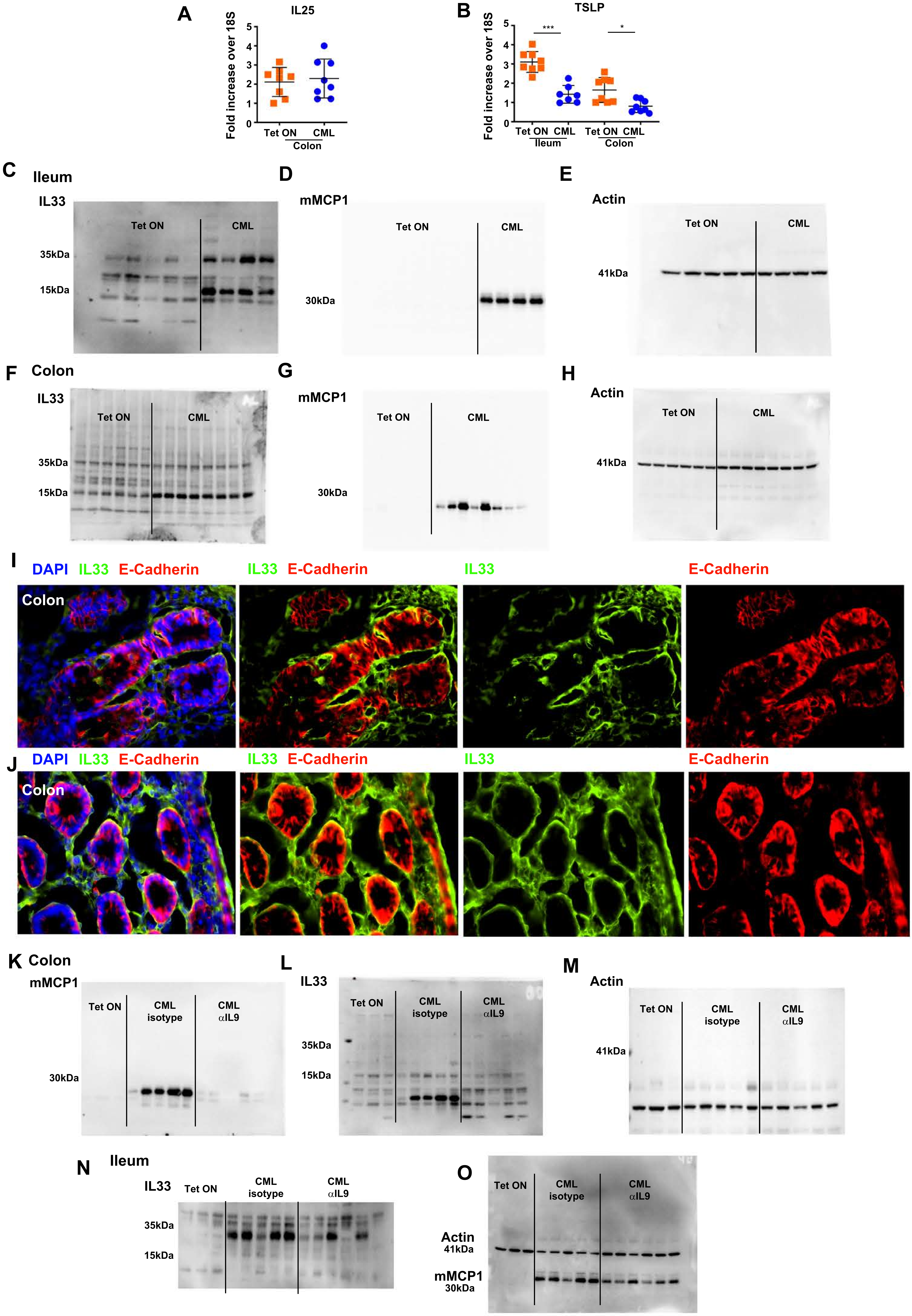

#### SUPPLEMENTAL FIGURE LEGENDS

Figure 1: **Tetracycline-off inducible murine transplantation model of CML.** (A) Breeding schematic and mechanism by which crossing of SCLtTa and bcr-abl mice creates mice that develop CML after withdrawal of tetracycline (Tet) from the drinking water. (B-K). Lethally irradiated (1100 cGy) FVB mice were transplanted with 10 x $10^6$ BM cells from SCLtTa / bcr-abl animals, and then maintained on or off Tet in the drinking water. (B). Serial weight curves of mice that were maintained on (n=8) or off Tet (n=8). (C, D). Serial total white blood cell and granulocyte counts in the peripheral blood at days 21, 37, and 52 after transplantation. Horizontal lines in panel C denote the lower and upper limits of normal for white blood cell counts. Data are from two-six experiments. (E). Spleen weight 28-35 days post transplantation. (F). Representative dot plot depicting CD11b and Gr-1 expression on spleen cells. (G). Absolute number of CD11b$^+$ Gr-1$^+$ cells in the spleen 28-35 days post transplantation. Data are from 2-4 experiments per panel. (H-K). Representative photomicrographs of spleen, mesenteric lymph node, liver and lung from mice with and without CML 35-45 days post transplantation. Insets in panels J and K depict collections of neutrophils in liver and lung, respectively. Statistics: ** $p<0.01$, ***$p<0.001$.

Figure 2: **Depletion of CD4$^+$ T cells has no effect on PCM or intestinal remodeling.** (A-H). Lethally irradiated FVB mice were transplanted with BM from SCLtTa/bcr-abl mice. Animals were either maintained on or taken off Tet and then treated two times per week with an isotype control or anti-CD4 antibody for 28 days. (A). Dot plot demonstrating CD4$^+$ TCRβ$^+$ T cells in the spleen of mice treated with an isotype control or anti-CD4 antibody. (B). Absolute number of Paneth cell in the colon and ileum. (C, D). Crypt1 and sPLA2 gene expression in the colon and ileum. (E). Small intestines and colon length. (F-H). mRNA expression of Fcer1-a (panel F), DLCK-1 (panel G), and IL-25 (panel H) in the ileum. Data are from two experiments. Statistics: *$p<0.05$, ** $p<0.01$, ***$p<0.001$, ****$p<0.0001$.

Figure 3: **IL-33 localizes to the intestinal epithelium in CML mice.** (A-J). Lethally irradiated FVB mice (n=8/group) were transplanted with SCLtTa/bcr-abl BM (10 x $10^6$) and maintained on Tet or taken off Tet. Colon and ileum samples were harvested on day 35 post-transplantation. (A, B). mRNA gene expression of IL-25 in the colon (panel E), and TSLP in the colon and ileum (panel F) of CML animals. (C-H). Immunoblots of IL-33 (long and short forms), mMCP1 and actin in the colon (panels C-E) and ileum (panels F-H) of CML mice. (I, J) Immunofluorescence staining in the colon showing merged expression of IL-33, E-cadherin and DAPI. Longitudinal (panel I) and cross sectional (panel J) tissue images are depicted. (K-O). Lethally irradiated FVB mice (n=8/group) were transplanted with SCLtTa/bcr-abl BM and maintained on Tet or taken off Tet. Animals taken off Tet were treated with anti-IL-9 antibody or an isotype control three times per week for three weeks. Colon and ileum tissue extracts were collected 21 days post transplantation. Immunoblots of IL-33, mMCP-1 and actin in the colon (panels K-M) and ileum (panels N, O). Statistics: *$p<0.05$, ***$p<0.001$.

#### SUPPLEMENTAL TABLE 1: FOLD CHANGE METABOLITES IN THE ILEUM OF CML MICE

| Super Pathway | Sub Pathway | Biochemical Name | CML_ISOTYPE FVB_SYN Fold change | p-value | q-value |
|---|---|---|---|---|---|
| Amino Acid | Histidine Metabolism | histamine | 144.53 | 1.17E-08 | 2.12E-08 |
| Lipid | Phospholipid Metabolism | glycerophosphoserine* | 62.32 | 5.00E-05 | 6.3E-14 |
| Amino Acid | Tyrosine Metabolism | p-cresol glucuronide* | 60.50 | 0.0007 | 0.0002 |
| Lipid | Phospholipid Metabolism | glycerophosphoinositol* | 45.96 | 3.00E-15 | 1.89E-13 |
| Nucleotide | Pyrimidine Metabolism, Orotate containing | dihydroorotate | 42.87 | 1.56E-09 | 4.2E-09 |
| Lipid | Lysoplasmalogen | 1-(1-enyl-palmitoyl)-GPC (P-16:0)* | 19.75 | 1.29E-09 | 3.68E-09 |
| Lipid | Long Chain Polyunsaturated Fatty Acid (n3 a | mead acid (20:3n9) | 19.21 | 3.64E-07 | 3.81E-07 |
| Amino Acid | Histidine Metabolism | 1-methyl-4-imidazoleacetate | 18.36 | 1.42E-11 | 9.49E-11 |
| Peptide | Dipeptide | valylglycine | 14.52 | 4.83E-06 | 3.1E-06 |
| Amino Acid | Histidine Metabolism | 1-methylhistamine | 13.32 | 5.43E-10 | 2.37E-09 |
| Lipid | Phospholipid Metabolism | trimethylamine N-oxide | 13.23 | 1.45E-10 | 8.01E-10 |
| Lipid | Fatty Acid Metabolism (Acyl Choline) | docosahexaenoylcholine | 12.06 | 4.93E-06 | 3.15E-06 |
| Lipid | Corticosteroids | corticosterone | 12.05 | 1.56E-05 | 8.38E-06 |
| Peptide | Acetylated Peptides | phenylacetylglycine | 11.22 | 5.47E-06 | 3.46E-06 |
| Lipid | Fatty Acid, Monohydroxy | 3-hydroxystearate | 10.27 | 1.05E-06 | 8.7E-07 |
| Amino Acid | Tryptophan Metabolism | serotonin | 9.98 | 1.06E-08 | 1.98E-08 |
| Lipid | Fatty Acid Metabolism (Acyl Choline) | stearoylcholine* | 9.51 | 5.29E-09 | 1.12E-08 |
| Amino Acid | Tyrosine Metabolism | phenol sulfate | 9.06 | 2.54E-06 | 1.86E-06 |
| Amino Acid | Tyrosine Metabolism | phenol glucuronide | 8.98 | 3.72E-05 | 1.77E-05 |
| Lipid | Long Chain Monounsaturated Fatty Acid | erucate (22:1n9) | 8.96 | 2.58E-06 | 1.86E-06 |
| Lipid | Fatty Acid Metabolism (Acyl Carnitine, Polyu | linolenoylcarnitine (C18:3)* | 8.15 | 8.75E-07 | 7.44E-07 |
| Lipid | Long Chain Polyunsaturated Fatty Acid (n3 a | docosadienoate (22:2n6) | 8.01 | 4.22E-07 | 4.24E-07 |
| Peptide | Dipeptide | isoleucylglycine | 7.90 | 7.72E-08 | 1.01E-07 |
| Peptide | Acetylated Peptides | phenylacetylcarnitine | 7.80 | 0.0007 | 0.0002 |
| Lipid | Eicosanoid | 12-HETE | 7.62 | 6.09E-09 | 1.24E-08 |
| Lipid | Lysoplasmalogen | 1-(1-enyl-oleoyl)-GPE (P-18:1)* | 7.61 | 1.29E-06 | 1.04E-06 |
| Lipid | Lysoplasmalogen | 1-(1-enyl-stearoyl)-GPE (P-18:0)* | 7.03 | 2.58E-06 | 1.86E-06 |
| Lipid | Lysoplasmalogen | 1-(1-enyl-palmitoyl)-GPE (P-16:0)* | 6.77 | 1.60E-06 | 1.25E-06 |
| Lipid | Fatty Acid Metabolism (Acyl Carnitine, Polyu | docosapentaenoylcarnitine (C22:5n3)* | 6.70 | 1.82E-08 | 3.04E-08 |
| Lipid | Sphingomyelins | sphingomyelin (d18:2/24:2)* | 6.61 | 8.95E-12 | 7.56E-11 |
| Nucleotide | Pyrimidine Metabolism, Uracil containing | uridine-2',3'-cyclic monophosphate | 6.58 | 2.56E-07 | 2.77E-07 |
| Lipid | Fatty Acid Metabolism (Acyl Choline) | palmitoylcholine | 6.48 | 5.63E-09 | 1.17E-08 |
| Peptide | Dipeptide | valylglutamine | 6.44 | 2.37E-08 | 3.85E-08 |
| Cofactors and Vitam | Tetrahydrobiopterin Metabolism | dihydrobiopterin | 6.32 | 2.68E-11 | 1.62E-10 |
| Xenobiotics | Benzoate Metabolism | p-cresol sulfate | 6.23 | 0.0069 | 0.0017 |
| Lipid | Fatty Acid Metabolism (Acyl Carnitine, Polyu | linoleoylcarnitine (C18:2)* | 6.11 | 9.90E-08 | 1.22E-07 |
| Lipid | Hexosylceramides (HCER) | glycosyl-N-nervonoyl-sphingosine (d18:1/2 | 5.91 | 3.65E-08 | 5.71E-08 |
| Lipid | Sphingomyelins | sphingomyelin (d18:2/14:0, d18:1/14:1)* | 5.72 | 7.56E-06 | 4.58E-06 |
| Lipid | Docosanoid | 14-HDoHE/17-HDoHE | 5.67 | 1.17E-09 | 3.68E-09 |
| Peptide | Dipeptide | tyrosylglycine | 5.63 | 1.29E-07 | 1.52E-07 |
| Amino Acid | Tryptophan Metabolism | 3-indoxyl sulfate | 5.62 | 1.83E-05 | 9.54E-06 |
| Lipid | Fatty Acid Metabolism (Acyl Carnitine, Mono | myristoleoylcarnitine (C14:1)* | 5.46 | 2.33E-07 | 2.57E-07 |
| Carbohydrate | Glycogen Metabolism | maltose | 5.46 | 0.0002 | 6.82E-05 |
| Lipid | Fatty Acid Metabolism (Acyl Carnitine, Mono | palmitoleoylcarnitine (C16:1)* | 5.44 | 1.23E-05 | 6.8E-06 |
| Lipid | Fatty Acid Metabolism (Acyl Carnitine, Mono | oleoylcarnitine (C18:1) | 5.44 | 2.09E-07 | 2.34E-07 |
| Peptide | Dipeptide | phenylalanylglycine | 5.42 | 3.86E-06 | 2.63E-06 |
| Lipid | Diacylglycerol | linoleoyl-docosahexaenoyl-glycerol (18:2/2 | 5.32 | 0.0014 | 0.0004 |
| Amino Acid | Urea cycle; Arginine and Proline Metabolism | N-methylproline | 5.27 | 4.12E-06 | 2.77E-06 |
| Lipid | Fatty Acid Metabolism (Acyl Carnitine, Polyu | dihomo-linoleoylcarnitine (C20:2)* | 5.23 | 2.00E-09 | 5.17E-09 |
| Peptide | Dipeptide | leucylglycine | 5.16 | 2.24E-08 | 3.68E-08 |
| Lipid | Fatty Acid Metabolism (Acyl Carnitine, Mono | erucoylcarnitine (C22:1)* | 5.16 | 1.22E-08 | 2.18E-08 |
| Amino Acid | Polyamine Metabolism | putrescine | 5.15 | 4.23E-06 | 2.83E-06 |
| Lipid | Phospholipid Metabolism | glycerophosphoethanolamine | 5.12 | 2.86E-12 | 3.29E-11 |
| Nucleotide | Pyrimidine Metabolism, Cytidine containing | cytidine 2',3'-cyclic monophosphate | 4.79 | 0.0001 | 4.99E-05 |

| | | | | | |
|---|---|---|---|---|---|
| Nucleotide | Dinucleotide | (3'-5')-uridylylcytidine* | 4.54 | 9.34E-05 | 3.97E-05 |
| Lipid | Fatty Acid, Dicarboxylate | 2-hydroxyadipate | 4.50 | 7.92E-05 | 3.42E-05 |
| Lipid | Long Chain Saturated Fatty Acid | arachidate (20:0) | 4.44 | 1.65E-05 | 8.69E-06 |
| Lipid | Eicosanoid | 5-HETE | 4.43 | 2.39E-07 | 2.61E-07 |
| Cofactors and Vitam | Tetrahydrobiopterin Metabolism | biopterin | 4.42 | 3.02E-10 | 1.41E-09 |
| Lipid | Fatty Acid Metabolism (Acyl Carnitine, Medi | laurylcarnitine (C12) | 4.39 | 9.92E-08 | 1.22E-07 |
| Lipid | Sphingomyelins | sphingomyelin (d18:1/22:2, d18:2/22:1, d1 | 4.36 | 8.00E-12 | 7.23E-11 |
| Peptide | Dipeptide | leucylglutamine* | 4.32 | 2.48E-09 | 6.03E-09 |
| Lipid | Fatty Acid Metabolism (Acyl Carnitine, Polyu | adrenoylcarnitine (C22:4)* | 4.21 | 4.46E-07 | 4.44E-07 |
| Lipid | Fatty Acid Metabolism (Acyl Carnitine, Mono | cis-4-decenoylcarnitine (C10:1) | 4.18 | 5.77E-05 | 2.59E-05 |
| Peptide | Dipeptide | threonylphenylalanine | 4.16 | 8.06E-06 | 4.8E-06 |
| Peptide | Dipeptide | tryptophylglycine | 4.15 | 6.96E-06 | 4.28E-06 |
| Lipid | Fatty Acid Metabolism (Acyl Carnitine, Polyu | docosadienoylcarnitine (C22:2)* | 4.13 | 8.47E-07 | 7.24E-07 |
| Peptide | Dipeptide | valylleucine | 4.12 | 1.42E-05 | 7.77E-06 |
| Lipid | Sterol | lanosterol | 4.08 | 0.0002 | 9.27E-05 |
| Lipid | Fatty Acid, Monohydroxy | 2-hydroxystearate | 4.05 | 7.83E-07 | 6.8E-07 |
| Amino Acid | Methionine, Cysteine, SAM and Taurine Met | methionine sulfone | 3.98 | 9.90E-06 | 5.64E-06 |
| Nucleotide | Pyrimidine Metabolism, Uracil containing | 3-ureidopropionate | 3.94 | 2.02E-05 | 1.04E-05 |
| Peptide | Acetylated Peptides | phenylacetyltaurine | 3.92 | 0.0021 | 0.0006 |
| Lipid | Fatty Acid Metabolism (Acyl Carnitine, Polyu | docosatrienoylcarnitine (C22:3)* | 3.91 | 4.84E-07 | 4.68E-07 |
| Amino Acid | Methionine, Cysteine, SAM and Taurine Met | cystathionine | 3.91 | 5.79E-08 | 8.14E-08 |
| Nucleotide | Pyrimidine Metabolism, Thymine containing | thymidine 5'-monophosphate | 3.86 | 4.42E-05 | 2.06E-05 |
| Xenobiotics | Food Component/Plant | 3-hydroxyindolin-2-one | 3.84 | 0.004 | 0.001 |
| Lipid | Fatty Acid Metabolism (Acyl Carnitine, Medi | decanoylcarnitine (C10) | 3.83 | 1.40E-06 | 1.12E-06 |
| Lipid | Fatty Acid Metabolism (Acyl Carnitine, Polyu | dihomo-linolenoylcarnitine (C20:3n3 or 6)* | 3.82 | 3.10E-09 | 7.4E-09 |
| Lipid | Glycerolipid Metabolism | glycerophosphoglycerol | 3.81 | 4.57E-09 | 1.03E-08 |
| Lipid | Fatty Acid Metabolism (Acyl Carnitine, Mono | undecenoylcarnitine (C11:1) | 3.75 | 0.01 | 0.0023 |
| Amino Acid | Phenylalanine Metabolism | phenyllactate (PLA) | 3.74 | 0.0986 | 0.017 |
| Lipid | Fatty Acid Metabolism (Acyl Choline) | arachidonoylcholine | 3.73 | 5.32E-05 | 2.43E-05 |
| Lipid | Eicosanoid | thromboxane B2 | 3.72 | 1.52E-05 | 8.24E-06 |
| Lipid | Long Chain Polyunsaturated Fatty Acid (n3 a | dihomo-linoleate (20:2n6) | 3.71 | 2.72E-05 | 1.35E-05 |
| Nucleotide | Dinucleotide | (3'-5')-uridylyluridine | 3.62 | 5.19E-05 | 2.38E-05 |
| Lipid | Dihydrosphingomyelins | myristoyl dihydrosphingomyelin (d18:0/14 | 3.61 | 2.68E-06 | 1.9E-06 |
| Lipid | Diacylglycerol | linoleoyl-arachidonoyl-glycerol (18:2/20:4) | 3.57 | 0.0357 | 0.007 |
| Lipid | Fatty Acid Metabolism (Acyl Carnitine, Mono | 5-dodecenoylcarnitine (C12:1) | 3.53 | 4.32E-05 | 2.02E-05 |
| Xenobiotics | Chemical | ectoine | 3.50 | 0.0056 | 0.0014 |
| Lipid | Eicosanoid | 15-HETE | 3.49 | 9.27E-07 | 7.77E-07 |
| Lipid | Phospholipid Metabolism | glycerophosphorylcholine (GPC) | 3.48 | 1.31E-09 | 3.68E-09 |
| Peptide | Dipeptide | alanylleucine | 3.46 | 2.19E-06 | 1.64E-06 |
| Amino Acid | Polyamine Metabolism | N-acetylputrescine | 3.46 | 4.28E-06 | 2.84E-06 |
| Lipid | Fatty Acid Metabolism (Acyl Carnitine, Mono | eicosenoylcarnitine (C20:1)* | 3.42 | 2.44E-09 | 6.03E-09 |
| Amino Acid | Histidine Metabolism | carnosine | 3.42 | 8.93E-08 | 1.15E-07 |
| Xenobiotics | Chemical | O-sulfo-L-tyrosine | 3.39 | 5.25E-07 | 4.96E-07 |
| Lipid | Hexosylceramides (HCER) | glycosyl ceramide (d18:2/24:1, d18:1/24:2 | 3.31 | 4.03E-05 | 1.9E-05 |
| Lipid | Fatty Acid Metabolism (Acyl Carnitine, Long | myristoylcarnitine (C14) | 3.28 | 1.58E-07 | 1.82E-07 |
| Lipid | Fatty Acid Metabolism (Acyl Carnitine, Medi | octanoylcarnitine (C8) | 3.27 | 0.0002 | 7.92E-05 |
| Lipid | Sphingomyelins | sphingomyelin (d18:1/20:1, d18:2/20:0)* | 3.24 | 1.43E-11 | 9.49E-11 |
| Lipid | Fatty Acid Metabolism (Acyl Carnitine, Medi | hexanoylcarnitine (C6) | 3.23 | 0.0004 | 0.0001 |
| Lipid | Long Chain Polyunsaturated Fatty Acid (n3 a | nisinate (24:6n3) | 3.22 | 1.00E-05 | 5.68E-06 |
| Nucleotide | Purine Metabolism, Adenine containing | adenosine-2',3'-cyclic monophosphate | 3.22 | 0.0001 | 5.09E-05 |
| Lipid | Sphingosines | eicosanoylsphingosine (d20:1)* | 3.18 | 4.94E-07 | 4.74E-07 |
| Lipid | Galactosyl Glycerolipids | 1-linoleoyl-digalactosylglycerol (18:2)* | 3.08 | 0.0317 | 0.0063 |
| Lipid | Sphingolipid Synthesis | sphinganine-1-phosphate | 3.08 | 0.0058 | 0.0014 |
| Peptide | Dipeptide Derivative | isoleucylhydroxyproline* | 3.05 | 0.0001 | 4.5E-05 |
| Lipid | Long Chain Monounsaturated Fatty Acid | eicosenoate (20:1) | 3.03 | 8.31E-05 | 3.57E-05 |
| Peptide | Dipeptide | phenylalanylalanine | 2.98 | 1.51E-07 | 1.75E-07 |
| Cofactors and Vitam | Nicotinate and Nicotinamide Metabolism | trigonelline (N'-methylnicotinate) | 2.96 | 0.0002 | 8.7E-05 |
| Peptide | Dipeptide | histidylalanine | 2.93 | 8.27E-06 | 4.85E-06 |

| | | | | | |
|---|---|---|---|---|---|
| Lipid | Eicosanoid | 15-deoxy delta-12,14-prostaglandin J2 | 2.93 | 2.54E-05 | 1.26E-05 |
| Amino Acid | Histidine Metabolism | 1-ribosyl-imidazoleacetate* | 2.93 | 5.61E-05 | 2.54E-05 |
| Amino Acid | Histidine Metabolism | 4-imidazoleacetate | 2.93 | 1.29E-05 | 7.04E-06 |
| Peptide | Dipeptide | glycylleucine | 2.91 | 7.65E-05 | 3.33E-05 |
| Lipid | Long Chain Polyunsaturated Fatty Acid (n3 a | eicosapentaenoate (EPA; 20:5n3) | 2.91 | 7.00E-05 | 3.05E-05 |
| Lipid | Sphingomyelins | sphingomyelin (d18:2/18:1)* | 2.91 | 9.32E-08 | 1.18E-07 |
| Amino Acid | Tyrosine Metabolism | O-methyltyrosine | 2.90 | 5.07E-09 | 1.11E-08 |
| Lipid | Long Chain Polyunsaturated Fatty Acid (n3 a | docosapentaenoate (n3 DPA; 22:5n3) | 2.86 | 0.0001 | 5.89E-05 |
| Lipid | Fatty Acid Metabolism (also BCAA Metabolis | propionylcarnitine (C3) | 2.85 | 0.0012 | 0.0004 |
| Nucleotide | Dinucleotide | (3'-5')-guanylylcytidine | 2.81 | 0.0085 | 0.002 |
| Lipid | Long Chain Polyunsaturated Fatty Acid (n3 a | adrenate (22:4n6) | 2.79 | 6.89E-05 | 3.02E-05 |
| Carbohydrate | Aminosugar Metabolism | N-acetylneuraminate | 2.78 | 1.12E-08 | 2.06E-08 |
| Peptide | Dipeptide | leucylalanine | 2.77 | 3.26E-05 | 1.57E-05 |
| Lipid | Fatty Acid Metabolism (also BCAA Metabolis | butyrylcarnitine (C4) | 2.73 | 0.0038 | 0.001 |
| Cofactors and Vitam | Pantothenate and CoA Metabolism | coenzyme A | 2.70 | 0.0036 | 0.0009 |
| Lipid | Eicosanoid | 12-HHTrE | 2.67 | 1.07E-07 | 1.29E-07 |
| Lipid | Long Chain Polyunsaturated Fatty Acid (n3 a | arachidonate (20:4n6) | 2.64 | 2.78E-06 | 1.96E-06 |
| Lipid | Sphingolipid Synthesis | sphinganine | 2.61 | 0.0014 | 0.0004 |
| Nucleotide | Pyrimidine Metabolism, Orotate containing | N-carbamoylaspartate | 2.58 | 7.84E-07 | 6.8E-07 |
| Lipid | Mevalonate Metabolism | 3-hydroxy-3-methylglutarate | 2.58 | 1.72E-08 | 2.94E-08 |
| Lipid | Fatty Acid Metabolism (Acyl Carnitine, Medi | nonanoylcarnitine (C9) | 2.55 | 0.005 | 0.0013 |
| Lipid | Fatty Acid Metabolism (Acyl Carnitine, Polyu | arachidonoylcarnitine (C20:4) | 2.54 | 9.67E-07 | 8.06E-07 |
| Nucleotide | Pyrimidine Metabolism, Uracil containing | pseudouridine | 2.54 | 6.04E-07 | 5.54E-07 |
| Amino Acid | Tryptophan Metabolism | indoxyl glucuronide | 2.53 | 0.0042 | 0.0011 |
| Energy | TCA Cycle | 2-methylcitrate/homocitrate | 2.51 | 0.033 | 0.0065 |
| Lipid | Fatty Acid Metabolism (Acyl Choline) | oleoylcholine | 2.50 | 0.0002 | 7.19E-05 |
| Lipid | Ceramides | ceramide (d18:2/24:1, d18:1/24:2)* | 2.50 | 3.47E-09 | 8E-09 |
| Cofactors and Vitam | Tocopherol Metabolism | gamma-tocotrienol | 2.49 | 0.0872 | 0.0154 |
| Lipid | Long Chain Saturated Fatty Acid | stearate (18:0) | 2.47 | 7.60E-06 | 4.58E-06 |
| Lipid | Fatty Acid Metabolism (Acyl Carnitine, Medi | cis-3,4-methyleneheptanoylcarnitine | 2.45 | 6.57E-05 | 2.89E-05 |
| Lipid | Sphingomyelins | sphingomyelin (d18:1/18:1, d18:2/18:0) | 2.44 | 1.25E-11 | 9.49E-11 |
| Xenobiotics | Drug - Topical Agents | salicylate | 2.44 | 0.0003 | 9.4E-05 |
| Amino Acid | Alanine and Aspartate Metabolism | N,N-dimethylalanine | 2.44 | 0.0033 | 0.0009 |
| Nucleotide | Pyrimidine Metabolism, Uracil containing | 5,6-dihydrouridine | 2.42 | 3.32E-08 | 5.26E-08 |
| Lipid | Fatty Acid Metabolism (Acyl Carnitine, Hydro | 3-hydroxyoleoylcarnitine | 2.41 | 4.62E-05 | 2.15E-05 |
| Nucleotide | Purine Metabolism, Adenine containing | adenosine 5'-diphosphate (ADP) | 2.40 | 0.0112 | 0.0025 |
| Peptide | Dipeptide | lysylleucine | 2.39 | 0.004 | 0.001 |
| Amino Acid | Glutathione Metabolism | 2-hydroxybutyrate/2-hydroxyisobutyrate | 2.39 | 4.69E-07 | 4.6E-07 |
| Cofactors and Vitam | Thiamine Metabolism | thiamin diphosphate | 2.37 | 0.0187 | 0.004 |
| Amino Acid | Tryptophan Metabolism | kynurenine | 2.34 | 0.0002 | 7.59E-05 |
| Amino Acid | Urea cycle; Arginine and Proline Metabolism | homocitrulline | 2.32 | 9.39E-06 | 5.45E-06 |
| Lipid | Fatty Acid Metabolism (Acyl Carnitine, Long | palmitoylcarnitine (C16) | 2.31 | 3.30E-06 | 2.28E-06 |
| Amino Acid | Leucine, Isoleucine and Valine Metabolism | alpha-hydroxyisovalerate | 2.30 | 2.17E-06 | 1.64E-06 |
| Nucleotide | Dinucleotide | (3'-5')-guanylyluridine | 2.27 | 0.0022 | 0.0006 |
| Nucleotide | Pyrimidine Metabolism, Orotate containing | orotidine | 2.25 | 7.21E-08 | 9.6E-08 |
| Nucleotide | Pyrimidine Metabolism, Uracil containing | 5-methyluridine (ribothymidine) | 2.24 | 0.0005 | 0.0002 |
| Amino Acid | Glutathione Metabolism | glutathione, reduced (GSH) | 2.21 | 0.0023 | 0.0006 |
| Carbohydrate | Pentose Phosphate Pathway | ribose 1-phosphate | 2.18 | 0.0006 | 0.0002 |
| Lipid | Fatty Acid Metabolism (Acyl Carnitine, Hydro | 3-hydroxydecanoylcarnitine | 2.15 | 2.86E-05 | 1.4E-05 |
| Amino Acid | Lysine Metabolism | 6-oxopiperidine-2-carboxylate | 2.15 | 0.0438 | 0.0084 |
| Xenobiotics | Food Component/Plant | stachydrine | 2.15 | 0.0138 | 0.0031 |
| Lipid | Fatty Acid Metabolism (Acyl Carnitine, Hydro | (R)-3-hydroxybutyrylcarnitine | 2.14 | 0.0015 | 0.0004 |
| Amino Acid | Alanine and Aspartate Metabolism | N-acetylaspartate (NAA) | 2.13 | 9.53E-06 | 5.51E-06 |
| Lipid | Diacylglycerol | linoleoyl-arachidonoyl-glycerol (18:2/20:4) | 2.12 | 0.0112 | 0.0025 |
| Peptide | Dipeptide | glycylvaline | 2.10 | 0.0039 | 0.001 |
| Lipid | Fatty Acid, Monohydroxy | 2-hydroxypalmitate | 2.10 | 0.0015 | 0.0004 |
| Amino Acid | Urea cycle; Arginine and Proline Metabolism | N-delta-acetylornithine | 2.10 | 4.39E-08 | 6.57E-08 |
| Lipid | Diacylglycerol | linoleoyl-docosahexaenoyl-glycerol (18:2/2 | 2.09 | 0.0422 | 0.0081 |

| | | | | | |
|---|---|---|---|---|---|
| Lipid | Sphingomyelins | sphingomyelin (d18:2/16:0, d18:1/16:1)* | 2.08 | 4.28E-08 | 6.52E-08 |
| Nucleotide | Purine Metabolism, Guanine containing | guanosine 2'-monophosphate (2'-GMP)* | 2.08 | 0.0281 | 0.0057 |
| Amino Acid | Lysine Metabolism | 5-(galactosylhydroxy)-L-lysine | 2.03 | 4.41E-08 | 6.57E-08 |
| Lipid | Sphingomyelins | sphingomyelin (d18:2/24:1, d18:1/24:2)* | 1.99 | 7.28E-08 | 9.6E-08 |
| Lipid | Fatty Acid Metabolism (Acyl Carnitine, Monc | nervonoylcarnitine (C24:1)* | 1.98 | 3.04E-05 | 1.48E-05 |
| Lipid | Long Chain Saturated Fatty Acid | margarate (17:0) | 1.98 | 0.0027 | 0.0007 |
| Amino Acid | Methionine, Cysteine, SAM and Taurine Met | N-acetylmethionine | 1.96 | 5.83E-05 | 2.61E-05 |
| Amino Acid | Leucine, Isoleucine and Valine Metabolism | 2-hydroxy-3-methylvalerate | 1.96 | 0.0035 | 0.0009 |
| Lipid | Inositol Metabolism | chiro-inositol | 1.96 | 0.0104 | 0.0024 |
| Amino Acid | Glutamate Metabolism | N-acetylglutamate | 1.95 | 7.63E-06 | 4.58E-06 |
| Amino Acid | Glutathione Metabolism | 4-hydroxy-nonenal-glutathione | 1.94 | 0.0192 | 0.0041 |
| Lipid | Fatty Acid, Monohydroxy | 3-hydroxyoleate* | 1.93 | 0.0004 | 0.0001 |
| Nucleotide | Dinucleotide | (3'-5')-adenylylcytidine | 1.92 | 0.0043 | 0.0011 |
| Lipid | Fatty Acid Metabolism (Acyl Carnitine, Short | acetylcarnitine (C2) | 1.91 | 0.0002 | 6.77E-05 |
| Nucleotide | Purine Metabolism, Adenine containing | adenosine 3'-monophosphate (3'-AMP) | 1.91 | 0.0067 | 0.0016 |
| Carbohydrate | Pentose Phosphate Pathway | 6-phosphogluconate | 1.90 | 0.0132 | 0.003 |
| Amino Acid | Polyamine Metabolism | N('1)-acetylspermidine | 1.89 | 9.57E-05 | 4.03E-05 |
| Lipid | Fatty Acid Metabolism (Acyl Carnitine, Long | arachidoylcarnitine (C20)* | 1.87 | 0.0003 | 9.38E-05 |
| Lipid | Docosanoid | 4-HDoHE | 1.87 | 0.0005 | 0.0002 |
| Nucleotide | Purine Metabolism, Adenine containing | adenosine 5'-monophosphate (AMP) | 1.86 | 6.13E-06 | 3.83E-06 |
| Nucleotide | Pyrimidine Metabolism, Uracil containing | uridine 5'-monophosphate (UMP) | 1.86 | 2.09E-07 | 2.34E-07 |
| Peptide | Dipeptide | glutaminylleucine | 1.85 | 0.0549 | 0.0102 |
| Lipid | Fatty Acid Metabolism (Acyl Carnitine, Polyu | docosahexaenoylcarnitine (C22:6)* | 1.85 | 0.0003 | 0.0001 |
| Amino Acid | Glycine, Serine and Threonine Metabolism | dimethylglycine | 1.85 | 4.51E-06 | 2.94E-06 |
| Amino Acid | Leucine, Isoleucine and Valine Metabolism | beta-hydroxyisovalerate | 1.83 | 0.0465 | 0.0088 |
| Lipid | Phospholipid Metabolism | cytidine-5'-diphosphoethanolamine | 1.80 | 0.0056 | 0.0014 |
| Lipid | Sphingomyelins | palmitoyl sphingomyelin (d18:1/16:0) | 1.80 | 8.63E-10 | 3.12E-09 |
| Amino Acid | Leucine, Isoleucine and Valine Metabolism | methylsuccinate | 1.80 | 0.1114 | 0.019 |
| Amino Acid | Glutamate Metabolism | glutamine | 1.79 | 0.0031 | 0.0008 |
| Cofactors and Vitam | Folate Metabolism | 5-methyltetrahydrofolate (5MeTHF) | 1.79 | 0.0024 | 0.0007 |
| Lipid | Sphingolipid Synthesis | sphingadienine | 1.79 | 0.0064 | 0.0016 |
| Amino Acid | Tryptophan Metabolism | N-formylkynurenine | 1.79 | 0.0137 | 0.0031 |
| Amino Acid | Histidine Metabolism | histidine methyl ester | 1.78 | 0.0006 | 0.0002 |
| Peptide | Dipeptide Derivative | leucylhydroxyproline* | 1.78 | 0.0448 | 0.0085 |
| Amino Acid | Polyamine Metabolism | diacetylspermidine* | 1.78 | 0.0025 | 0.0007 |
| Amino Acid | Tryptophan Metabolism | C-glycosyltryptophan | 1.78 | 4.83E-06 | 3.1E-06 |
| Lipid | Long Chain Polyunsaturated Fatty Acid (n3 a | heneicosapentaenoate (21:5n3) | 1.78 | 0.0008 | 0.0003 |
| Lipid | Fatty Acid Metabolism (Acyl Carnitine, Long | pentadecanoylcarnitine (C15)* | 1.76 | 0.0001 | 4.43E-05 |
| Nucleotide | Dinucleotide | (3'-5')-adenylyluridine | 1.76 | 0.0109 | 0.0025 |
| Amino Acid | Histidine Metabolism | imidazole propionate | 1.76 | 0.4717 | 0.0679 |
| Amino Acid | Glutathione Metabolism | CoA-glutathione* | 1.75 | 0.0584 | 0.0108 |
| Lipid | Glycerolipid Metabolism | glycerol 3-phosphate | 1.75 | 0.0191 | 0.0041 |
| Lipid | Fatty Acid Metabolism (Acyl Choline) | dihomo-linolenoyl-choline | 1.74 | 0.0025 | 0.0007 |
| Cofactors and Vitam | Nicotinate and Nicotinamide Metabolism | N1-Methyl-4-pyridone-3-carboxamide | 1.74 | 0.0042 | 0.0011 |
| Peptide | Dipeptide | glycylisoleucine | 1.72 | 0.0174 | 0.0038 |
| Amino Acid | Methionine, Cysteine, SAM and Taurine Met | N-acetylmethionine sulfoxide | 1.72 | 0.0553 | 0.0102 |
| Lipid | Fatty Acid Metabolism (Acyl Choline) | palmitoloelycholine | 1.72 | 0.0006 | 0.0002 |
| Xenobiotics | Food Component/Plant | pyrraline | 1.72 | 0.1116 | 0.019 |
| Amino Acid | Methionine, Cysteine, SAM and Taurine Met | taurocyamine | 1.72 | 0.0001 | 5.18E-05 |
| Xenobiotics | Chemical | perfluorooctanesulfonate (PFOS) | 1.71 | 0.0009 | 0.0003 |
| Amino Acid | Methionine, Cysteine, SAM and Taurine Met | 2,3-dihydroxy-5-methylthio-4-pentenoate | 1.71 | 6.02E-06 | 3.79E-06 |
| Amino Acid | Histidine Metabolism | 3-methylhistidine | 1.70 | 0.0005 | 0.0002 |
| Lipid | Fatty Acid, Dicarboxylate | octadecadienedioate (C18:2-DC)* | 1.69 | 0.0033 | 0.0009 |
| Lipid | Eicosanoid | 6-keto prostaglandin F1alpha | 1.69 | 0.0004 | 0.0002 |
| Lipid | Long Chain Polyunsaturated Fatty Acid (n3 a | docosapentaenoate (n6 DPA; 22:5n6) | 1.69 | 0.0032 | 0.0009 |
| Cofactors and Vitam | Vitamin A Metabolism | retinol (Vitamin A) | 1.68 | 0.0138 | 0.0031 |
| Lipid | Sphingomyelins | stearoyl sphingomyelin (d18:1/18:0) | 1.68 | 1.60E-05 | 8.5E-06 |
| Nucleotide | Purine Metabolism, Adenine containing | adenosine 2'-monophosphate (2'-AMP) | 1.68 | 0.0284 | 0.0058 |

| | | | | | |
|---|---|---|---|---|---|
| Cofactors and Vitam | Nicotinate and Nicotinamide Metabolism | N1-Methyl-2-pyridone-5-carboxamide | 1.68 | 0.0106 | 0.0024 |
| Cofactors and Vitam | Nicotinate and Nicotinamide Metabolism | quinolinate | 1.68 | 0.019 | 0.004 |
| Nucleotide | Pyrimidine Metabolism, Cytidine containing | 2'-O-methylcytidine | 1.67 | 0.0019 | 0.0005 |
| Nucleotide | Purine Metabolism, Guanine containing | guanosine 5'-monophosphate (5'-GMP) | 1.67 | 0.0002 | 7.6E-05 |
| Xenobiotics | Food Component/Plant | tartronate (hydroxymalonate) | 1.66 | 0.0196 | 0.0041 |
| Lipid | Ceramides | N-behenoyl-sphingadienine (d18:2/22:0)* | 1.64 | 0.0003 | 9.76E-05 |
| Lipid | Hexosylceramides (HCER) | glycosyl-N-palmitoyl-sphingosine (d18:1/1 | 1.64 | 0.0015 | 0.0004 |
| Amino Acid | Histidine Metabolism | 1-methyl-5-imidazolelactate | 1.64 | 0.0377 | 0.0073 |
| Amino Acid | Glutamate Metabolism | glutamate, gamma-methyl ester | 1.64 | 7.30E-06 | 4.44E-06 |
| Carbohydrate | Aminosugar Metabolism | N-glycolylneuraminate | 1.64 | 0.0088 | 0.0021 |
| Lipid | Eicosanoid | leukotriene B4 | 1.63 | 0.0189 | 0.004 |
| Amino Acid | Tyrosine Metabolism | 3-(4-hydroxyphenyl)lactate | 1.62 | 0.0136 | 0.003 |
| Carbohydrate | Glycolysis, Gluconeogenesis, and Pyruvate M | fructose 1,6-bisphosphate | 1.62 | 0.0447 | 0.0085 |
| Lipid | Long Chain Saturated Fatty Acid | nonadecanoate (19:0) | 1.60 | 0.0186 | 0.004 |
| Amino Acid | Urea cycle; Arginine and Proline Metabolism | citrulline | 1.60 | 0.0005 | 0.0002 |
| Nucleotide | Purine Metabolism, Adenine containing | 2'-deoxyadenosine 5'-monophosphate | 1.60 | 0.045 | 0.0086 |
| Nucleotide | Pyrimidine Metabolism, Uracil containing | 2'-deoxyuridine | 1.59 | 0.0097 | 0.0023 |
| Amino Acid | Leucine, Isoleucine and Valine Metabolism | 3-hydroxyisobutyrate | 1.59 | 0.0301 | 0.006 |
| Lipid | Sterol | desmosterol | 1.58 | 0.0611 | 0.0112 |
| Amino Acid | Polyamine Metabolism | 4-acetamidobutanoate | 1.58 | 0.013 | 0.0029 |
| Amino Acid | Lysine Metabolism | N6,N6,N6-trimethyllysine | 1.57 | 0.0103 | 0.0024 |
| Lipid | Dihydrosphingomyelins | palmitoyl dihydrosphingomyelin (d18:0/16 | 1.57 | 0.0002 | 7.51E-05 |
| Amino Acid | Lysine Metabolism | N6-methyllysine | 1.57 | 1.47E-06 | 1.16E-06 |
| Lipid | Sphingomyelins | sphingomyelin (d18:1/24:1, d18:2/24:0)* | 1.56 | 0.0005 | 0.0002 |
| Nucleotide | Pyrimidine Metabolism, Cytidine containing | cytosine | 1.56 | 0.6351 | 0.0878 |
| Amino Acid | Methionine, Cysteine, SAM and Taurine Met | 2-hydroxy-4-(methylthio)butanoic acid | 1.56 | 0.0981 | 0.017 |
| Lipid | Long Chain Polyunsaturated Fatty Acid (n3 a | tetradecadienoate (14:2)* | 1.56 | 0.0002 | 7.41E-05 |
| Amino Acid | Histidine Metabolism | anserine | 1.55 | 0.0058 | 0.0014 |
| Carbohydrate | Pentose Metabolism | ribitol | 1.55 | 0.0036 | 0.001 |
| Amino Acid | Glycine, Serine and Threonine Metabolism | N-acetylserine | 1.55 | 0.0023 | 0.0006 |
| Xenobiotics | Food Component/Plant | methyl glucopyranoside (alpha + beta) | 1.55 | 0.0013 | 0.0004 |
| Amino Acid | Glutamate Metabolism | N-methyl-GABA | 1.55 | 0.0004 | 0.0001 |
| Lipid | Fatty Acid, Dicarboxylate | 3-hydroxyadipate | 1.55 | 0.0558 | 0.0103 |
| Carbohydrate | Glycolysis, Gluconeogenesis, and Pyruvate M | glucose 6-phosphate | 1.53 | 0.0318 | 0.0063 |
| Lipid | Fatty Acid Metabolism (Acyl Choline) | linoleoylcholine* | 1.52 | 0.032 | 0.0064 |
| Nucleotide | Pyrimidine Metabolism, Thymine containing | thymine | 1.52 | 0.0292 | 0.0059 |
| Xenobiotics | Benzoate Metabolism | hippurate | 1.51 | 0.0994 | 0.0171 |
| Amino Acid | Lysine Metabolism | saccharopine | 1.51 | 0.1442 | 0.0238 |
| Nucleotide | Pyrimidine Metabolism, Orotate containing | orotate | 1.50 | 0.0245 | 0.0051 |
| Lipid | Eicosanoid | prostaglandin A2 | 1.49 | 0.0102 | 0.0024 |
| Lipid | Fatty Acid Metabolism (Acyl Carnitine, Long | stearoylcarnitine (C18) | 1.48 | 0.0142 | 0.0031 |
| Lipid | Sphingomyelins | sphingomyelin (d18:1/22:1, d18:2/22:0, d1 | 1.48 | 2.21E-05 | 1.12E-05 |
| Amino Acid | Alanine and Aspartate Metabolism | N-acetylalanine | 1.48 | 0.0094 | 0.0022 |
| Xenobiotics | Food Component/Plant | dihydroferulate | 1.48 | 0.2331 | 0.0365 |
| Amino Acid | Lysine Metabolism | 5-hydroxylysine | 1.47 | 0.0055 | 0.0014 |
| Lipid | Phospholipid Metabolism | cytidine 5'-diphosphocholine | 1.47 | 0.0251 | 0.0052 |
| Nucleotide | Pyrimidine Metabolism, Cytidine containing | cytidine 5'-monophosphate (5'-CMP) | 1.47 | 0.0004 | 0.0001 |
| Nucleotide | Purine Metabolism, (Hypo)Xanthine/Inosine | inosine 5'-monophosphate (IMP) | 1.46 | 0.1073 | 0.0184 |
| Lipid | Fatty Acid, Monohydroxy | 3-hydroxymyristate | 1.46 | 0.0006 | 0.0002 |
| Nucleotide | Pyrimidine Metabolism, Uracil containing | uridine 5'-diphosphate (UDP) | 1.46 | 0.047 | 0.0088 |
| Cofactors and Vitam | Ascorbate and Aldarate Metabolism | 2-O-methylascorbic acid | 1.46 | 2.86E-05 | 1.4E-05 |
| Lipid | Hexosylceramides (HCER) | glycosyl-N-stearoyl-sphingosine (d18:1/18: | 1.45 | 0.1896 | 0.0304 |
| Nucleotide | Purine Metabolism, Adenine containing | adenosine 3',5'-diphosphate | 1.45 | 0.0527 | 0.0098 |
| Xenobiotics | Drug - Topical Agents | 2,6-dihydroxybenzoic acid | 1.45 | 0.2782 | 0.0428 |
| Nucleotide | Pyrimidine Metabolism, Thymine containing | thymidine | 1.43 | 2.48E-05 | 1.24E-05 |
| Lipid | Long Chain Polyunsaturated Fatty Acid (n3 a | dihomo-linolenate (20:3n3 or n6) | 1.43 | 0.0129 | 0.0029 |
| Lipid | Fatty Acid, Monohydroxy | 3-hydroxyhexanoate | 1.42 | 0.2073 | 0.0328 |
| Cofactors and Vitam | Hemoglobin and Porphyrin Metabolism | bilirubin (Z,Z) | 1.42 | 0.7291 | 0.0988 |

| | | | | | |
|---|---|---|---|---|---|
| Xenobiotics | Chemical | dimethyl sulfone | 1.41 | 0.0399 | 0.0077 |
| Amino Acid | Lysine Metabolism | pipecolate | 1.41 | 6.09E-05 | 2.69E-05 |
| Lipid | Sphingomyelins | hydroxypalmitoyl sphingomyelin (d18:1/16 | 1.40 | 0.0029 | 0.0008 |
| Energy | Oxidative Phosphorylation | acetylphosphate | 1.40 | 0.0195 | 0.0041 |
| Lipid | Endocannabinoid | N-oleoyltaurine | 1.39 | 0.3359 | 0.0505 |
| Amino Acid | Methionine, Cysteine, SAM and Taurine Met | N-formylmethionine | 1.39 | 0.021 | 0.0044 |
| Amino Acid | Leucine, Isoleucine and Valine Metabolism | alpha-hydroxyisocaproate | 1.39 | 0.641 | 0.0884 |
| Lipid | Fatty Acid Metabolism (Acyl Carnitine, Long | margaroylcarnitine (C17)* | 1.38 | 0.0075 | 0.0018 |
| Amino Acid | Leucine, Isoleucine and Valine Metabolism | isovalerylcarnitine (C5) | 1.38 | 0.2156 | 0.034 |
| Amino Acid | Leucine, Isoleucine and Valine Metabolism | 3-methylglutaconate | 1.36 | 0.1263 | 0.0212 |
| Cofactors and Vitam | Ascorbate and Aldarate Metabolism | dehydroascorbate | 1.35 | 0.0169 | 0.0037 |
| Amino Acid | Glutathione Metabolism | ophthalmate | 1.35 | 0.0958 | 0.0166 |
| Lipid | Fatty Acid, Amino | 2-aminoheptanoate | 1.35 | 0.0312 | 0.0062 |
| Lipid | Fatty Acid, Monohydroxy | 3-hydroxyoctanoate | 1.35 | 0.114 | 0.0193 |
| Lipid | Mevalonate Metabolism | mevalonate | 1.34 | 0.2181 | 0.0343 |
| Amino Acid | Glutamate Metabolism | gamma-aminobutyrate (GABA) | 1.34 | 0.0048 | 0.0012 |
| Carbohydrate | Aminosugar Metabolism | N-acetyl-glucosamine 1-phosphate | 1.34 | 0.0011 | 0.0003 |
| Cofactors and Vitam | Tocopherol Metabolism | gamma-tocopherol/beta-tocopherol | 1.33 | 0.5262 | 0.0748 |
| Lipid | Long Chain Monounsaturated Fatty Acid | 10-nonadecenoate (19:1n9) | 1.33 | 0.146 | 0.024 |
| Energy | TCA Cycle | succinate | 1.33 | 0.0152 | 0.0033 |
| Lipid | Short Chain Fatty Acid | butyrate/isobutyrate (4:0) | 1.32 | 0.0721 | 0.0131 |
| Amino Acid | Glutamate Metabolism | N-acetylglutamine | 1.32 | 0.0776 | 0.0139 |
| Lipid | Long Chain Monounsaturated Fatty Acid | oleate/vaccenate (18:1) | 1.31 | 0.0205 | 0.0043 |
| Lipid | Ketone Bodies | 3-hydroxybutyrate (BHBA) | 1.31 | 0.6722 | 0.092 |
| Cofactors and Vitam | Pantothenate and CoA Metabolism | 3'-dephosphocoenzyme A | 1.29 | 0.095 | 0.0165 |
| Amino Acid | Polyamine Metabolism | spermidine | 1.28 | 0.0207 | 0.0043 |
| Lipid | Fatty Acid, Monohydroxy | 3-hydroxylaurate | 1.28 | 0.0765 | 0.0138 |
| Amino Acid | Glycine, Serine and Threonine Metabolism | N-acetylglycine | 1.27 | 0.0539 | 0.01 |
| Amino Acid | Lysine Metabolism | N,N,N-trimethyl-5-aminovalerate | 1.27 | 0.0147 | 0.0032 |
| Lipid | Long Chain Saturated Fatty Acid | palmitate (16:0) | 1.27 | 0.0345 | 0.0068 |
| Lipid | Sphingosines | sphingosine | 1.27 | 0.0719 | 0.013 |
| Energy | TCA Cycle | itaconate | 1.27 | 0.2067 | 0.0327 |
| Lipid | Lysophospholipid | 1-stearoyl-GPI (18:0) | 1.26 | 0.1602 | 0.0261 |
| Amino Acid | Glycine, Serine and Threonine Metabolism | N-acetylthreonine | 1.26 | 0.1433 | 0.0237 |
| Amino Acid | Urea cycle; Arginine and Proline Metabolism | ornithine | 1.26 | 0.008 | 0.0019 |
| Lipid | Fatty Acid, Monohydroxy | 2-hydroxydecanoate | 1.26 | 0.1075 | 0.0184 |
| Nucleotide | Purine Metabolism, (Hypo)Xanthine/Inosine | 2'-deoxyinosine | 1.25 | 0.0978 | 0.0169 |
| Lipid | Short Chain Fatty Acid | valerate (5:0) | 1.25 | 0.281 | 0.0432 |
| Xenobiotics | Food Component/Plant | 3-indoleglyoxylic acid | 1.25 | 0.1514 | 0.0248 |
| Lipid | Long Chain Polyunsaturated Fatty Acid (n3 a | stearidonate (18:4n3) | 1.25 | 0.1511 | 0.0248 |
| Amino Acid | Urea cycle; Arginine and Proline Metabolism | 3-amino-2-piperidone | 1.24 | 0.2669 | 0.0413 |
| Carbohydrate | Nucleotide Sugar | UDP-glucose | 1.24 | 0.034 | 0.0067 |
| Lipid | Lactosylceramides (LCER) | lactosyl-N-palmitoyl-sphingosine (d18:1/16 | 1.24 | 0.0154 | 0.0034 |
| Energy | TCA Cycle | aconitate [cis or trans] | 1.24 | 0.0265 | 0.0054 |
| Lipid | Fatty Acid Metabolism (Acyl Carnitine, Hydro | 3-hydroxypalmitoylcarnitine | 1.23 | 0.134 | 0.0223 |
| Lipid | Primary Bile Acid Metabolism | glycocholate | 1.23 | 0.5467 | 0.0772 |
| Lipid | Sphingomyelins | sphingomyelin (d18:1/14:0, d16:1/16:0)* | 1.22 | 0.0043 | 0.0011 |
| Lipid | Phospholipid Metabolism | phosphoethanolamine | 1.22 | 0.0034 | 0.0009 |
| Amino Acid | Alanine and Aspartate Metabolism | asparagine | 1.22 | 0.1454 | 0.024 |
| Nucleotide | Purine Metabolism, (Hypo)Xanthine/Inosine | allantoin | 1.22 | 0.0124 | 0.0028 |
| Nucleotide | Purine Metabolism, Adenine containing | 2'-deoxyadenosine | 1.21 | 0.0838 | 0.0149 |
| Amino Acid | Methionine, Cysteine, SAM and Taurine Met | lanthionine | 1.21 | 0.464 | 0.0671 |
| Amino Acid | Leucine, Isoleucine and Valine Metabolism | beta-hydroxyisovaleroylcarnitine | 1.20 | 0.0341 | 0.0067 |
| Lipid | Fatty Acid, Dihydroxy | 2,4-dihydroxybutyrate | 1.20 | 0.0389 | 0.0075 |
| Cofactors and Vitam | Pantothenate and CoA Metabolism | pantothenate | 1.20 | 0.0304 | 0.0061 |
| Xenobiotics | Benzoate Metabolism | 4-hydroxyhippurate | 1.20 | 0.5775 | 0.0811 |
| Amino Acid | Leucine, Isoleucine and Valine Metabolism | tiglylcarnitine (C5:1-DC) | 1.19 | 0.2465 | 0.0383 |
| Lipid | Plasmalogen | 1-(1-enyl-palmitoyl)-2-arachidonoyl-GPC (P | 1.19 | 0.0617 | 0.0113 |

| | | | | | |
|---|---|---|---|---|---|
| Lipid | Long Chain Polyunsaturated Fatty Acid (n3 a | docosahexaenoate (DHA; 22:6n3) | 1.19 | 0.117 | 0.0197 |
| Carbohydrate | Fructose, Mannose and Galactose Metabolis | galactose 1-phosphate | 1.19 | 0.2429 | 0.0379 |
| Carbohydrate | Glycolysis, Gluconeogenesis, and Pyruvate M | 3-phosphoglycerate | 1.18 | 0.155 | 0.0253 |
| Xenobiotics | Chemical | glycerol 2-phosphate | 1.18 | 0.9275 | 0.1217 |
| Lipid | Fatty Acid, Monohydroxy | 2-hydroxyoctanoate | **1.18** | 0.0932 | 0.0163 |
| Nucleotide | Pyrimidine Metabolism, Cytidine containing | 2'-deoxycytidine | 1.17 | 0.3746 | 0.0556 |
| Lipid | Phosphatidylcholine (PC) | 1,2-dipalmitoyl-GPC (16:0/16:0) | **1.17** | 0.0937 | 0.0163 |
| Amino Acid | Leucine, Isoleucine and Valine Metabolism | isovalerate (i5:0) | 1.17 | 0.2933 | 0.0448 |
| Nucleotide | Pyrimidine Metabolism, Uracil containing | 2'-O-methyluridine | 1.17 | 0.2043 | 0.0324 |
| Amino Acid | Methionine, Cysteine, SAM and Taurine Met | N-acetyltaurine | **1.17** | 0.0797 | 0.0142 |
| Xenobiotics | Food Component/Plant | 2-aminophenol sulfate | 1.17 | 0.4231 | 0.0619 |
| Lipid | Dihydrosphingomyelins | behenoyl dihydrosphingomyelin (d18:0/22 | 1.16 | 0.69 | 0.0942 |
| Lipid | Fatty Acid, Dicarboxylate | 2-hydroxyglutarate | 1.16 | 0.2003 | 0.032 |
| Lipid | Plasmalogen | 1-(1-enyl-stearoyl)-2-arachidonoyl-GPE (P- | 1.15 | 0.1154 | 0.0195 |
| Amino Acid | Lysine Metabolism | 2-aminoadipate | **1.15** | 0.0382 | 0.0074 |
| Nucleotide | Purine Metabolism, Adenine containing | adenosine | 1.15 | 0.2802 | 0.0431 |
| Nucleotide | Pyrimidine Metabolism, Thymine containing | 3-aminoisobutyrate | 1.15 | 0.2767 | 0.0427 |
| Lipid | Fatty Acid, Monohydroxy | 2-hydroxyheptanoate* | 1.15 | 0.5365 | 0.076 |
| Lipid | Monoacylglycerol | 1-meadoylglycerol (20:3n9)* | 1.14 | 0.8147 | 0.1089 |
| Amino Acid | Methionine, Cysteine, SAM and Taurine Met | S-adenosylmethionine (SAM) | **1.14** | 0.0293 | 0.0059 |
| Lipid | Fatty Acid, Dicarboxylate | 3-methylglutarate/2-methylglutarate | 1.14 | 0.5833 | 0.0816 |
| Lipid | Neurotransmitter | acetylcholine | 1.13 | 0.228 | 0.0358 |
| Carbohydrate | Pentose Phosphate Pathway | sedoheptulose-7-phosphate | 1.13 | 0.3219 | 0.0486 |
| Cofactors and Vitam | Nicotinate and Nicotinamide Metabolism | 1-methylnicotinamide | 1.13 | 0.3657 | 0.0544 |
| Partially Characteriz | Partially Characterized Molecules | glutamine_degradant* | 1.13 | 0.3923 | 0.0579 |
| Peptide | Polypeptide | val-val-ala | 1.12 | 0.3429 | 0.0514 |
| Amino Acid | Glycine, Serine and Threonine Metabolism | threonine | 1.12 | 0.1832 | 0.0295 |
| Xenobiotics | Food Component/Plant | erythritol | 1.12 | 0.5511 | 0.0777 |
| Nucleotide | Pyrimidine Metabolism, Cytidine containing | cytidine | 1.11 | 0.3472 | 0.0518 |
| Xenobiotics | Food Component/Plant | 2,3-dihydroxyisovalerate | 1.11 | 0.7232 | 0.0983 |
| Amino Acid | Methionine, Cysteine, SAM and Taurine Met | hypotaurine | 1.11 | 0.2437 | 0.0379 |
| Amino Acid | Glycine, Serine and Threonine Metabolism | betaine | 1.10 | 0.167 | 0.027 |
| Amino Acid | Tyrosine Metabolism | tyrosine | 1.10 | 0.3157 | 0.0477 |
| Lipid | Fatty Acid, Monohydroxy | 3-hydroxydecanoate | 1.10 | 0.3534 | 0.0526 |
| Amino Acid | Glutamate Metabolism | glutamate | **1.10** | 0.0554 | 0.0102 |
| Xenobiotics | Benzoate Metabolism | guaiacol sulfate | 1.10 | 0.5744 | 0.0807 |
| Cofactors and Vitam | Nicotinate and Nicotinamide Metabolism | adenosine 5'-diphosphoribose (ADP-ribose | 1.09 | 0.5791 | 0.0812 |
| Amino Acid | Histidine Metabolism | 1-methylhistidine | 1.09 | 0.7254 | 0.0985 |
| Cofactors and Vitam | Ascorbate and Aldarate Metabolism | ascorbate (Vitamin C) | 1.08 | 0.4031 | 0.0593 |
| Nucleotide | Pyrimidine Metabolism, Uracil containing | 3-(3-amino-3-carboxypropyl)uridine* | 1.08 | 0.6506 | 0.0896 |
| Lipid | Medium Chain Fatty Acid | heptanoate (7:0) | 1.08 | 0.5947 | 0.0831 |
| Amino Acid | Histidine Metabolism | N-acetylhistamine | 1.07 | 0.3227 | 0.0487 |
| Carbohydrate | Nucleotide Sugar | UDP-glucuronate | 1.07 | 0.3985 | 0.0588 |
| Amino Acid | Methionine, Cysteine, SAM and Taurine Met | S-adenosylhomocysteine (SAH) | 1.07 | 0.1313 | 0.0219 |
| Nucleotide | Pyrimidine Metabolism, Cytidine containing | 2'-deoxycytidine 5'-monophosphate | 1.07 | 0.4492 | 0.0651 |
| Amino Acid | Glycine, Serine and Threonine Metabolism | serine | 1.06 | 0.4436 | 0.0645 |
| Lipid | Long Chain Monounsaturated Fatty Acid | 10-heptadecenoate (17:1n7) | 1.06 | 0.5402 | 0.0764 |
| Amino Acid | Urea cycle; Arginine and Proline Metabolism | proline | 1.06 | 0.479 | 0.0689 |
| Energy | Oxidative Phosphorylation | phosphate | 1.06 | 0.2383 | 0.0372 |
| Nucleotide | Purine Metabolism, (Hypo)Xanthine/Inosine | urate | 1.06 | 0.4451 | 0.0647 |
| Amino Acid | Creatine Metabolism | guanidinoacetate | 1.06 | 0.5809 | 0.0813 |
| Lipid | Medium Chain Fatty Acid | caprylate (8:0) | 1.06 | 0.6678 | 0.0915 |
| Xenobiotics | Benzoate Metabolism | 3-phenylpropionate (hydrocinnamate) | 1.05 | 0.4025 | 0.0593 |
| Amino Acid | Creatine Metabolism | creatine | **1.05** | 0.0834 | 0.0148 |
| Carbohydrate | Aminosugar Metabolism | erythronate* | 1.05 | 0.4905 | 0.0703 |
| Carbohydrate | Aminosugar Metabolism | glucosamine-6-phosphate | 1.05 | 0.5286 | 0.0749 |
| Lipid | Fatty Acid Metabolism (Acyl Carnitine, Long | behenoylcarnitine (C22)* | 1.04 | 0.756 | 0.102 |
| Lipid | Fatty Acid Metabolism (Acyl Carnitine, Hydro | (S)-3-hydroxybutyrylcarnitine | 1.04 | 0.9246 | 0.1215 |

| | | | | | |
|---|---|---|---|---|---|
| Nucleotide | Dinucleotide | (3'-5')-cytidylyluridine* | 1.04 | 0.6603 | 0.0907 |
| Xenobiotics | Food Component/Plant | homostachydrine* | 1.04 | 0.8942 | 0.1182 |
| Cofactors and Vitam | Vitamin B6 Metabolism | pyridoxamine phosphate | 1.04 | 0.6041 | 0.0843 |
| Cofactors and Vitam | Vitamin D Metabolism | vitamin D3 sulfate | 1.03 | 0.889 | 0.1177 |
| Cofactors and Vitam | Tocopherol Metabolism | alpha-tocopherol | 1.03 | 0.6644 | 0.0911 |
| Amino Acid | Leucine, Isoleucine and Valine Metabolism | ethylmalonate | 1.03 | 0.8138 | 0.1089 |
| Lipid | Diacylglycerol | oleoyl-arachidonoyl-glycerol (18:1/20:4) [2 | 1.03 | 0.8905 | 0.1178 |
| Amino Acid | Glutathione Metabolism | glutathione, oxidized (GSSG) | 1.02 | 0.696 | 0.0947 |
| Lipid | Fatty Acid, Dihydroxy | 2S,3R-dihydroxybutyrate | 1.02 | 0.9538 | 0.125 |
| Amino Acid | Tryptophan Metabolism | N-acetylkynurenine (2) | 1.02 | 0.9256 | 0.1215 |
| Amino Acid | Glycine, Serine and Threonine Metabolism | sarcosine | 1.01 | 0.9165 | 0.1206 |
| Lipid | Fatty Acid, Monohydroxy | alpha-hydroxycaproate | 1.01 | 0.8751 | 0.1162 |
| Lipid | Lysophospholipid | 1-oleoyl-GPG (18:1)* | 1.01 | 0.4247 | 0.0621 |
| Lipid | Sterol | cholesterol | 1.01 | 0.8441 | 0.1125 |
| Cofactors and Vitam | Vitamin B6 Metabolism | pyridoxal | 1.01 | 0.7265 | 0.0986 |
| Nucleotide | Pyrimidine Metabolism, Uracil containing | uracil | 1.01 | 0.7397 | 0.1 |
| Lipid | Long Chain Polyunsaturated Fatty Acid (n3 a | linoleate (18:2n6) | 1.01 | 0.8475 | 0.1127 |
| Lipid | Long Chain Polyunsaturated Fatty Acid (n3 a | linolenate [alpha or gamma; (18:3n3 or 6)] | 1.01 | 0.73 | 0.0988 |
| Amino Acid | Lysine Metabolism | 5-aminovalerate | 1.01 | 0.8965 | 0.1183 |
| Xenobiotics | Food Component/Plant | cinnamoylglycine | 1.00 | | |
| Nucleotide | Purine Metabolism, Adenine containing | adenylosuccinate | 1.00 | 0.2338 | 0.0366 |
| Carbohydrate | Nucleotide Sugar | UDP-N-acetylglucosamine/galactosamine | 1.00 | 0.996 | 0.1298 |
| Nucleotide | Purine Metabolism, Guanine containing | 2'-deoxyguanosine | 0.99 | 0.8768 | 0.1164 |
| Amino Acid | Lysine Metabolism | N-acetyl-cadaverine | 0.99 | 0.6931 | 0.0944 |
| Nucleotide | Pyrimidine Metabolism, Uracil containing | uridine | 0.99 | 0.968 | 0.1266 |
| Amino Acid | Glycine, Serine and Threonine Metabolism | glycine | 0.99 | 0.903 | 0.1191 |
| Amino Acid | Histidine Metabolism | histidine | 0.99 | 0.997 | 0.1299 |
| Nucleotide | Dinucleotide | (3'-5')-guanylyladenosine* | 0.99 | 0.9821 | 0.1283 |
| Xenobiotics | Food Component/Plant | formononetin | 0.99 | 0.4692 | 0.0677 |
| Carbohydrate | Nucleotide Sugar | UDP-galactose | 0.98 | 0.7793 | 0.1048 |
| Nucleotide | Pyrimidine Metabolism, Uracil containing | beta-alanine | 0.98 | 0.7523 | 0.1016 |
| Lipid | Phospholipid Metabolism | choline | 0.98 | 0.6893 | 0.0942 |
| Cofactors and Vitam | Vitamin B6 Metabolism | pyridoxamine | 0.97 | 0.9941 | 0.1297 |
| Lipid | Carnitine Metabolism | carnitine | 0.96 | 0.6298 | 0.0873 |
| Amino Acid | Phenylalanine Metabolism | phenylalanine | 0.96 | 0.8128 | 0.1089 |
| Nucleotide | Purine Metabolism, Adenine containing | adenosine 3',5'-cyclic monophosphate (cAI | 0.96 | 0.4433 | 0.0645 |
| Xenobiotics | Chemical | 4-chlorobenzoic acid | 0.96 | 0.2682 | 0.0414 |
| Amino Acid | Tryptophan Metabolism | indoleacetate | 0.95 | 0.9664 | 0.1265 |
| Amino Acid | Histidine Metabolism | 1-methyl-5-imidazoleacetate | 0.95 | 0.9161 | 0.1206 |
| Lipid | Medium Chain Fatty Acid | 10-undecenoate (11:1n1) | 0.95 | 0.6353 | 0.0878 |
| Carbohydrate | Glycolysis, Gluconeogenesis, and Pyruvate M | lactate | 0.95 | 0.5721 | 0.0805 |
| Xenobiotics | Benzoate Metabolism | benzoate | 0.95 | 0.3108 | 0.0472 |
| Amino Acid | Methionine, Cysteine, SAM and Taurine Met | taurine | 0.95 | 0.2945 | 0.0449 |
| Nucleotide | Purine Metabolism, (Hypo)Xanthine/Inosine | N1-methylinosine | 0.94 | 0.8024 | 0.1078 |
| Cofactors and Vitam | Nicotinate and Nicotinamide Metabolism | nicotinamide | 0.94 | 0.4835 | 0.0694 |
| Lipid | Lysophospholipid | 1-linolenoyl-GPG (18:3)* | 0.94 | 0.4641 | 0.0671 |
| Lipid | Sterol | 4-cholesten-3-one | 0.93 | 0.4026 | 0.0593 |
| Lipid | Ceramides | N-stearoyl-sphingadienine (d18:2/18:0)* | 0.93 | 0.4294 | 0.0627 |
| Nucleotide | Purine Metabolism, (Hypo)Xanthine/Inosine | inosine | 0.93 | 0.2026 | 0.0322 |
| Cofactors and Vitam | Ascorbate and Aldarate Metabolism | oxalate (ethanedioate) | 0.93 | 0.2033 | 0.0323 |
| Amino Acid | Lysine Metabolism | N6-acetyllysine | 0.93 | 0.6509 | 0.0896 |
| Carbohydrate | Nucleotide Sugar | guanosine 5'-diphospho-fucose | 0.93 | 0.344 | 0.0514 |
| Lipid | Dihydrosphingomyelins | sphingomyelin (d18:0/20:0, d16:0/22:0)* | 0.93 | 0.5141 | 0.0734 |
| Amino Acid | Methionine, Cysteine, SAM and Taurine Met | 3-sulfo-L-alanine | 0.93 | 0.6044 | 0.0843 |
| Lipid | Fatty Acid Metabolism (Acyl Carnitine, Long | lignoceroylcarnitine (C24)* | 0.92 | 0.4929 | 0.0706 |
| Amino Acid | Methionine, Cysteine, SAM and Taurine Met | 5-methylthioribose** | 0.92 | 0.4706 | 0.0679 |
| Amino Acid | Urea cycle; Arginine and Proline Metabolism | trans-4-hydroxyproline | 0.92 | 0.8794 | 0.1166 |
| Amino Acid | Lysine Metabolism | lysine | 0.92 | 0.328 | 0.0494 |

| | | | | | |
|---|---|---|---|---|---|
| Amino Acid | Methionine, Cysteine, SAM and Taurine Met | methionine sulfoxide | 0.92 | 0.6927 | 0.0944 |
| Amino Acid | Lysine Metabolism | N6,N6-dimethyllysine | 0.92 | 0.5184 | 0.0739 |
| Nucleotide | Purine and Pyrimidine Metabolism | methylphosphate | 0.91 | 0.6094 | 0.0849 |
| Lipid | Lysophospholipid | 1-oleoyl-GPI (18:1) | 0.90 | 0.7669 | 0.1034 |
| Lipid | Lysophospholipid | 1-linoleoyl-GPG (18:2)* | 0.90 | 0.4147 | 0.0608 |
| Nucleotide | Purine Metabolism, (Hypo)Xanthine/Inosine | xanthosine | 0.90 | 0.2923 | 0.0448 |
| Energy | TCA Cycle | citraconate/glutaconate | 0.90 | 0.8474 | 0.1127 |
| Xenobiotics | Food Component/Plant | pheophorbide A | 0.90 | 0.463 | 0.0671 |
| Amino Acid | Leucine, Isoleucine and Valine Metabolism | N-acetylvaline | 0.89 | 0.5014 | 0.0716 |
| Lipid | Phosphatidylethanolamine (PE) | 1-palmitoyl-2-stearoyl-GPE (16:0/18:0)* | 0.89 | 0.773 | 0.1041 |
| Amino Acid | Tryptophan Metabolism | indoleacetylglycine | 0.89 | 0.4299 | 0.0627 |
| Amino Acid | Tryptophan Metabolism | indolelactate | 0.89 | 0.3502 | 0.0522 |
| Carbohydrate | Aminosugar Metabolism | N-acetylglucosamine/N-acetylgalactosamir | 0.88 | 0.3156 | 0.0477 |
| Cofactors and Vitam | Ascorbate and Aldarate Metabolism | threonate | 0.88 | 0.1867 | 0.03 |
| Amino Acid | Leucine, Isoleucine and Valine Metabolism | leucine | 0.88 | 0.1425 | 0.0236 |
| Lipid | Ceramide PEs | palmitoyl-sphingosine-phosphoethanolam | 0.88 | 0.4049 | 0.0595 |
| Amino Acid | Urea cycle; Arginine and Proline Metabolism | N-monomethylarginine | 0.88 | 0.5484 | 0.0774 |
| Amino Acid | Urea cycle; Arginine and Proline Metabolism | dimethylarginine (SDMA + ADMA) | 0.88 | 0.6173 | 0.0858 |
| Xenobiotics | Benzoate Metabolism | 3-(3-hydroxyphenyl)propionate | 0.88 | 0.807 | 0.1082 |
| Amino Acid | Guanidino and Acetamido Metabolism | 1-methylguanidine | 0.88 | 0.3156 | 0.0477 |
| Xenobiotics | Chemical | sulfate* | 0.87 | 0.1921 | 0.0308 |
| Lipid | Plasmalogen | 1-(1-enyl-palmitoyl)-2-palmitoleoyl-GPC (P | 0.87 | 0.2924 | 0.0448 |
| Amino Acid | Alanine and Aspartate Metabolism | alanine | 0.87 | 0.1204 | 0.0202 |
| Energy | TCA Cycle | citrate | **0.87** | 0.0093 | 0.0022 |
| Lipid | Fatty Acid, Dihydroxy | 3,4-dihydroxybutyrate | 0.87 | 0.3843 | 0.0568 |
| Lipid | Sphingomyelins | sphingomyelin (d18:2/23:1)* | 0.87 | 0.2408 | 0.0376 |
| Xenobiotics | Food Component/Plant | N-carboxymethylalanine | 0.87 | 0.344 | 0.0514 |
| Lipid | Fatty Acid, Dihydroxy | 2R,3R-dihydroxybutyrate | **0.87** | 0.0875 | 0.0155 |
| Lipid | Endocannabinoid | N-stearoyltaurine | 0.87 | 0.3431 | 0.0514 |
| Amino Acid | Urea cycle; Arginine and Proline Metabolism | pro-hydroxy-pro | 0.86 | 0.8067 | 0.1082 |
| Amino Acid | Methionine, Cysteine, SAM and Taurine Met | methionine | 0.86 | 0.2931 | 0.0448 |
| Cofactors and Vitam | Riboflavin Metabolism | flavin adenine dinucleotide (FAD) | **0.86** | 0.0007 | 0.0002 |
| Peptide | Modified Peptides | N,N-dimethyl-pro-pro | **0.86** | 0.0512 | 0.0096 |
| Lipid | Fatty Acid, Dicarboxylate | adipate (C6-DC) | 0.86 | 0.3784 | 0.056 |
| Amino Acid | Urea cycle; Arginine and Proline Metabolism | arginine | 0.85 | 0.1181 | 0.0199 |
| Amino Acid | Glutathione Metabolism | 3'-dephospho-CoA-glutathione* | 0.85 | 0.2318 | 0.0364 |
| Amino Acid | Alanine and Aspartate Metabolism | N-acetylasparagine | 0.85 | 0.6536 | 0.0898 |
| Lipid | Dihydroceramides | N-palmitoyl-sphinganine (d18:0/16:0) | 0.85 | 0.2445 | 0.038 |
| Amino Acid | Leucine, Isoleucine and Valine Metabolism | isobutyrylglycine | 0.84 | 0.5242 | 0.0746 |
| Amino Acid | Polyamine Metabolism | 5-methylthioadenosine (MTA) | **0.84** | 0.0597 | 0.011 |
| Carbohydrate | Aminosugar Metabolism | N-acetylglucosaminylasparagine | **0.84** | 0.0664 | 0.0121 |
| Lipid | Fatty Acid, Monohydroxy | 8-hydroxyoctanoate | 0.84 | 0.3145 | 0.0477 |
| Lipid | Fatty Acid Synthesis | malonate | 0.83 | 0.6281 | 0.0872 |
| Lipid | Lysophospholipid | 2-palmitoleoyl-GPC (16:1)* | 0.83 | 0.6137 | 0.0854 |
| Amino Acid | Methionine, Cysteine, SAM and Taurine Met | S-methylcysteine | 0.83 | 0.1382 | 0.023 |
| Carbohydrate | Pentose Metabolism | ribonate | 0.83 | 0.8209 | 0.1096 |
| Amino Acid | Leucine, Isoleucine and Valine Metabolism | isobutyrylcarnitine (C4) | 0.82 | 0.2081 | 0.0329 |
| Amino Acid | Tryptophan Metabolism | kynurenate | 0.82 | 0.4802 | 0.069 |
| Nucleotide | Purine Metabolism, (Hypo)Xanthine/Inosine | xanthine | **0.82** | 0.0236 | 0.0049 |
| Cofactors and Vitam | Nicotinate and Nicotinamide Metabolism | nicotinamide adenine dinucleotide (NAD+) | **0.82** | 0.0921 | 0.0162 |
| Xenobiotics | Benzoate Metabolism | 4-hydroxybenzoate | 0.82 | 0.6337 | 0.0878 |
| Amino Acid | Glutathione Metabolism | S-methylglutathione | 0.81 | 0.202 | 0.0322 |
| Lipid | Fatty Acid, Branched | (14 or 15)-methylpalmitate (a17:0 or i17:0 | 0.81 | 0.3748 | 0.0556 |
| Nucleotide | Purine Metabolism, Adenine containing | N6-carbamoylthreonyladenosine | **0.81** | 0.0179 | 0.0039 |
| Amino Acid | Leucine, Isoleucine and Valine Metabolism | isoleucine | **0.81** | 0.0443 | 0.0085 |
| Lipid | Sphingomyelins | lignoceroyl sphingomyelin (d18:1/24:0) | 0.81 | 0.1089 | 0.0186 |
| Amino Acid | Histidine Metabolism | imidazole lactate | **0.81** | 0.0102 | 0.0024 |
| Lipid | Plasmalogen | 1-(1-enyl-palmitoyl)-2-palmitoyl-GPC (P-16 | **0.80** | 0.0874 | 0.0155 |

| | | | | | |
|---|---|---|---|---|---|
| Lipid | Plasmalogen | 1-(1-enyl-palmitoyl)-2-oleoyl-GPC (P-16:0/1 | 0.80 | 0.0147 | 0.0032 |
| Lipid | Sphingolipid Synthesis | phytosphingosine | 0.80 | 0.1984 | 0.0317 |
| Lipid | Phosphatidylcholine (PC) | 1-palmitoyl-2-stearoyl-GPC (16:0/18:0) | 0.80 | 0.1166 | 0.0197 |
| Lipid | Phospholipid Metabolism | choline phosphate | 0.80 | 0.0096 | 0.0022 |
| Lipid | Fatty Acid, Dicarboxylate | dodecanedioate (C12-DC) | 0.80 | 0.1177 | 0.0198 |
| Carbohydrate | Nucleotide Sugar | cytidine 5'-monophospho-N-acetylneuram | 0.80 | 0.0006 | 0.0002 |
| Amino Acid | Tryptophan Metabolism | 5-hydroxytryptophol | 0.79 | 0.3025 | 0.0461 |
| Amino Acid | Leucine, Isoleucine and Valine Metabolism | 2-methylbutyrylcarnitine (C5) | 0.79 | 0.1608 | 0.0261 |
| Cofactors and Vitam | Thiamine Metabolism | thiamin monophosphate | 0.79 | 0.0116 | 0.0026 |
| Lipid | Dihydroceramides | N-stearoyl-phytosphingosine (t18:0/18:0)* | 0.79 | 0.0781 | 0.014 |
| Nucleotide | Dinucleotide | (3'-5')-adenylylguanosine* | 0.79 | 0.6412 | 0.0884 |
| Cofactors and Vitam | Thiamine Metabolism | thiamin (Vitamin B1) | 0.78 | 0.1103 | 0.0188 |
| Amino Acid | Alanine and Aspartate Metabolism | aspartate | 0.78 | 0.0109 | 0.0025 |
| Lipid | Monoacylglycerol | 1-docosahexaenoylglycerol (22:6) | 0.78 | 0.2158 | 0.034 |
| Amino Acid | Methionine, Cysteine, SAM and Taurine Met | N-acetylcysteine | 0.78 | 0.0398 | 0.0077 |
| Lipid | Fatty Acid, Monohydroxy | 13-HODE + 9-HODE | 0.78 | 0.1036 | 0.0178 |
| Lipid | Ceramides | N-palmitoyl-sphingadienine (d18:2/16:0)* | 0.77 | 0.0004 | 0.0002 |
| Lipid | Sphingomyelins | sphingomyelin (d17:2/16:0, d18:2/15:0)* | 0.77 | 0.143 | 0.0237 |
| Lipid | Phosphatidylethanolamine (PE) | 1,2-dipalmitoyl-GPE (16:0/16:0)* | 0.77 | 0.1655 | 0.0268 |
| Lipid | Inositol Metabolism | myo-inositol | 0.76 | 0.0009 | 0.0003 |
| Energy | TCA Cycle | succinylcarnitine (C4-DC) | 0.76 | 0.0006 | 0.0002 |
| Lipid | Fatty Acid, Monohydroxy | 2-hydroxynervonate* | 0.76 | 0.1583 | 0.0258 |
| Nucleotide | Purine Metabolism, Guanine containing | guanosine | 0.76 | 0.014 | 0.0031 |
| Lipid | Lysophospholipid | 1-palmitoyl-GPI (16:0) | 0.75 | 0.1791 | 0.0288 |
| Cofactors and Vitam | Hemoglobin and Porphyrin Metabolism | protoporphyrin IX | 0.75 | 0.2086 | 0.0329 |
| Carbohydrate | Glycolysis, Gluconeogenesis, and Pyruvate M | glucose | 0.75 | 0.0657 | 0.012 |
| Lipid | Long Chain Polyunsaturated Fatty Acid (n3 a | heptadecatrienoate (17:3)* | 0.75 | 0.1489 | 0.0244 |
| Lipid | Medium Chain Fatty Acid | caproate (6:0) | 0.75 | 0.1789 | 0.0288 |
| Amino Acid | Lysine Metabolism | cadaverine | 0.74 | 0.0478 | 0.009 |
| Nucleotide | Purine Metabolism, Adenine containing | N1-methyladenosine | 0.74 | 0.0006 | 0.0002 |
| Amino Acid | Leucine, Isoleucine and Valine Metabolism | valine | 0.74 | 0.0204 | 0.0043 |
| Amino Acid | Creatine Metabolism | creatinine | 0.74 | 0.0683 | 0.0124 |
| Nucleotide | Dinucleotide | (3'-5')-adenylyladenosine* | 0.74 | 0.1931 | 0.0309 |
| Amino Acid | Urea cycle; Arginine and Proline Metabolism | carboxy-methyl-arginine | 0.74 | 0.0898 | 0.0158 |
| Carbohydrate | Fructose, Mannose and Galactose Metabolis | galactitol (dulcitol) | 0.73 | 0.8254 | 0.1101 |
| Lipid | Fatty Acid Synthesis | malonylcarnitine | 0.73 | 0.0027 | 0.0007 |
| Lipid | Endocannabinoid | N-palmitoyltaurine | 0.73 | 0.1309 | 0.0219 |
| Carbohydrate | Aminosugar Metabolism | N-acetylglucosamine 6-phosphate | 0.73 | 0.0453 | 0.0086 |
| Nucleotide | Pyrimidine Metabolism, Thymine containing | 5,6-dihydrothymine | 0.73 | 0.5609 | 0.079 |
| Cofactors and Vitam | Tocopherol Metabolism | delta-tocopherol | 0.72 | 0.4122 | 0.0605 |
| Amino Acid | Tyrosine Metabolism | vanillactate | 0.72 | 0.2561 | 0.0397 |
| Cofactors and Vitam | Nicotinate and Nicotinamide Metabolism | nicotinamide riboside | 0.72 | 0.3672 | 0.0546 |
| Amino Acid | Glutathione Metabolism | cysteinylglycine | 0.72 | 0.0112 | 0.0025 |
| Amino Acid | Urea cycle; Arginine and Proline Metabolism | N-acetylcitrulline | 0.72 | 0.0004 | 0.0001 |
| Amino Acid | Tryptophan Metabolism | indolepropionate | 0.72 | 0.0779 | 0.014 |
| Lipid | Fatty Acid, Dihydroxy | 9,10-DiHOME | 0.72 | 0.1165 | 0.0197 |
| Lipid | Sphingomyelins | sphingomyelin (d17:1/16:0, d18:1/15:0, d1 | 0.71 | 0.0073 | 0.0018 |
| Nucleotide | Purine Metabolism, Adenine containing | adenine | 0.71 | 0.0203 | 0.0043 |
| Lipid | Eicosanoid | prostaglandin F2alpha | 0.71 | 0.0241 | 0.005 |
| Nucleotide | Purine Metabolism, Adenine containing | 1-methyladenine | 0.71 | 0.3067 | 0.0467 |
| Lipid | Fatty Acid, Monohydroxy | 2-hydroxyarachidate* | 0.71 | 0.1693 | 0.0273 |
| Lipid | Phosphatidylinositol (PI) | 1-stearoyl-2-arachidonoyl-GPI (18:0/20:4) | 0.71 | 0.0006 | 0.0002 |
| Lipid | Phosphatidylcholine (PC) | 1-palmitoyl-2-docosahexaenoyl-GPC (16:0/ | 0.71 | 2.38E-05 | 1.19E-05 |
| Lipid | Long Chain Polyunsaturated Fatty Acid (n3 a | hexadecadienoate (16:2n6) | 0.71 | 0.1466 | 0.0241 |
| Nucleotide | Purine Metabolism, Guanine containing | 7-methylguanine | 0.70 | 0.0201 | 0.0042 |
| Lipid | Endocannabinoid | oleoyl ethanolamide | 0.70 | 0.0722 | 0.0131 |
| Amino Acid | Glutathione Metabolism | S-(1,2-dicarboxyethyl)glutathione | 0.70 | 0.0084 | 0.002 |
| Xenobiotics | Benzoate Metabolism | 4-methylcatechol sulfate | 0.70 | 0.5272 | 0.0748 |

| | | | | | |
|---|---|---|---|---|---|
| Amino Acid | Tyrosine Metabolism | o-Tyrosine | 0.69 | 0.0929 | 0.0163 |
| Amino Acid | Tryptophan Metabolism | tryptophan | 0.69 | 0.029 | 0.0059 |
| Xenobiotics | Food Component/Plant | glycitin (glycitein 7-O-glucoside) | 0.68 | 0.5203 | 0.0741 |
| Lipid | Endocannabinoid | stearoyl ethanolamide | 0.68 | 0.0055 | 0.0014 |
| Amino Acid | Urea cycle; Arginine and Proline Metabolism | N-alpha-acetylornithine | 0.68 | 0.0157 | 0.0034 |
| Lipid | Plasmalogen | 1-(1-enyl-palmitoyl)-2-oleoyl-GPE (P-16:0/1 | 0.68 | 0.0013 | 0.0004 |
| Lipid | Sphingomyelins | sphingomyelin (d18:1/17:0, d17:1/18:0, d1 | 0.68 | 0.0013 | 0.0004 |
| Carbohydrate | Fructose, Mannose and Galactose Metabolis | fructose | 0.68 | 0.1523 | 0.0249 |
| Amino Acid | Polyamine Metabolism | N-carbamoylputrescine | 0.67 | 0.4954 | 0.0709 |
| Amino Acid | Tryptophan Metabolism | picolinate | 0.67 | 0.0261 | 0.0053 |
| Lipid | Lysophospholipid | 1-palmitoyl-GPG (16:0)* | 0.67 | 0.1683 | 0.0272 |
| Lipid | Secondary Bile Acid Metabolism | glycohyocholate | 0.67 | 0.1584 | 0.0258 |
| Cofactors and Vitam | Vitamin B6 Metabolism | pyridoxate | 0.66 | 0.0317 | 0.0063 |
| Amino Acid | Tryptophan Metabolism | oxindolylalanine | 0.66 | 0.0424 | 0.0081 |
| Nucleotide | Purine Metabolism, Adenine containing | methylthioadenosine sulfoxide | 0.66 | 0.099 | 0.0171 |
| Lipid | Fatty Acid, Dicarboxylate | pimelate (C7-DC) | 0.66 | 0.2882 | 0.0442 |
| Lipid | Long Chain Monounsaturated Fatty Acid | palmitoleate (16:1n7) | 0.66 | 0.0269 | 0.0055 |
| Amino Acid | Glutamate Metabolism | 4-hydroxyglutamate | 0.66 | 0.034 | 0.0067 |
| Amino Acid | Leucine, Isoleucine and Valine Metabolism | N-acetylisoleucine | 0.65 | 0.0616 | 0.0113 |
| Lipid | Fatty Acid, Dicarboxylate | dodecenedioate (C12:1-DC)* | 0.65 | 0.0795 | 0.0142 |
| Lipid | Sterol | 7-alpha-hydroxy-3-oxo-4-cholestenoate (7- | 0.65 | 0.0043 | 0.0011 |
| Amino Acid | Urea cycle; Arginine and Proline Metabolism | urea | 0.65 | 0.0072 | 0.0017 |
| Lipid | Sphingosines | hexadecasphingosine (d16:1)* | 0.64 | 0.0421 | 0.0081 |
| Xenobiotics | Chemical | thioproline | 0.64 | 0.0019 | 0.0005 |
| Nucleotide | Dinucleotide | (3'-5')-cytidylylcytidine* | 0.64 | 0.0725 | 0.0131 |
| Amino Acid | Lysine Metabolism | N6-carboxyethyllysine | 0.64 | 0.0064 | 0.0016 |
| Lipid | Fatty Acid, Monohydroxy | 4-hydroxybutyrate (GHB) | 0.63 | 0.0445 | 0.0085 |
| Lipid | Long Chain Polyunsaturated Fatty Acid (n3 a | hexadecatrienoate (16:3n3) | 0.63 | 0.0936 | 0.0163 |
| Xenobiotics | Benzoate Metabolism | catechol sulfate | 0.63 | 0.33 | 0.0497 |
| Lipid | Diacylglycerol | palmitoyl-arachidonoyl-glycerol (16:0/20:4 | 0.63 | 0.0275 | 0.0056 |
| Xenobiotics | Food Component/Plant | 4-hydroxycinnamate | 0.63 | 0.162 | 0.0263 |
| Amino Acid | Leucine, Isoleucine and Valine Metabolism | N-acetylleucine | 0.62 | 0.0475 | 0.0089 |
| Carbohydrate | Disaccharides and Oligosaccharides | lactose | 0.62 | 0.4455 | 0.0647 |
| Lipid | Carnitine Metabolism | deoxycarnitine | 0.62 | 0.0005 | 0.0002 |
| Carbohydrate | Pentose Metabolism | ribulose/xylulose | 0.62 | 0.0455 | 0.0086 |
| Lipid | Fatty Acid, Branched | (16 or 17)-methylstearate (a19:0 or i19:0) | 0.61 | 0.0973 | 0.0169 |
| Amino Acid | Tyrosine Metabolism | N-acetyltyrosine | 0.61 | 0.0178 | 0.0038 |
| Cofactors and Vitam | Nicotinate and Nicotinamide Metabolism | nicotinamide ribonucleotide (NMN) | 0.61 | 0.1421 | 0.0236 |
| Amino Acid | Tyrosine Metabolism | N-formylphenylalanine | 0.60 | 0.0335 | 0.0066 |
| Lipid | Monoacylglycerol | 1-palmitoylglycerol (16:0) | 0.60 | 0.0292 | 0.0059 |
| Lipid | Phosphatidylcholine (PC) | 1-stearoyl-2-docosahexaenoyl-GPC (18:0/2 | 0.60 | 2.88E-07 | 3.04E-07 |
| Carbohydrate | Pentose Metabolism | arabitol/xylitol | 0.59 | 0.0071 | 0.0017 |
| Cofactors and Vitam | Nicotinate and Nicotinamide Metabolism | nicotinate ribonucleoside | 0.59 | 0.0839 | 0.0149 |
| Amino Acid | Histidine Metabolism | N-acetylhistidine | 0.58 | 0.0367 | 0.0072 |
| Lipid | Fatty Acid, Dicarboxylate | glutarate (C5-DC) | 0.58 | 0.0025 | 0.0007 |
| Lipid | Galactosyl Glycerolipids | galactosylglycerol | 0.58 | 0.0518 | 0.0097 |
| Xenobiotics | Food Component/Plant | daidzein | 0.58 | 0.075 | 0.0135 |
| Lipid | Glycerolipid Metabolism | glycerol | 0.58 | 0.0007 | 0.0002 |
| Lipid | Primary Bile Acid Metabolism | taurocholate | 0.58 | 0.0899 | 0.0158 |
| Nucleotide | Purine Metabolism, Guanine containing | N2,N2-dimethylguanosine | 0.57 | 0.0742 | 0.0134 |
| Amino Acid | Phenylalanine Metabolism | N-acetylphenylalanine | 0.57 | 0.0105 | 0.0024 |
| Xenobiotics | Food Component/Plant | equol | 0.56 | 0.0494 | 0.0092 |
| Amino Acid | Glutamate Metabolism | N-acetyl-aspartyl-glutamate (NAAG) | 0.56 | 0.0324 | 0.0064 |
| Amino Acid | Tryptophan Metabolism | 5-hydroxyindoleacetate | 0.56 | 0.0025 | 0.0007 |
| Lipid | Sterol | cholesterol sulfate | 0.56 | 1.02E-07 | 1.24E-07 |
| Lipid | Sphingomyelins | sphingomyelin (d18:1/20:0, d16:1/22:0)* | 0.56 | 0.0003 | 9.68E-05 |
| Lipid | Endocannabinoid | N-linoleoyltaurine* | 0.56 | 0.1066 | 0.0183 |
| Amino Acid | Methionine, Cysteine, SAM and Taurine Met | cysteine sulfinic acid | 0.56 | 0.0017 | 0.0005 |

| | | | | | |
|---|---|---|---|---|---|
| Amino Acid | Histidine Metabolism | trans-urocanate | 0.55 | 0.0993 | 0.0171 |
| Carbohydrate | Advanced Glycation End-product | N6-carboxymethyllysine | 0.55 | 0.0008 | 0.0002 |
| Lipid | Fatty Acid, Dicarboxylate | hexadecanedioate (C16-DC) | 0.55 | 0.0002 | 6.34E-05 |
| Lipid | Lysophospholipid | 1-arachidonoyl-GPE (20:4n6)* | 0.54 | 0.0004 | 0.0002 |
| Nucleotide | Purine Metabolism, (Hypo)Xanthine/Inosine | hypoxanthine | 0.54 | 0.0001 | 4.43E-05 |
| Carbohydrate | Pentose Metabolism | lyxonate | 0.54 | 0.0052 | 0.0013 |
| Cofactors and Vitam | Ascorbate and Aldarate Metabolism | gulonate* | 0.54 | 0.0002 | 8.52E-05 |
| Lipid | Dihydroceramides | N-palmitoyl-phytosphingosine (t18:0/16:0) | 0.54 | 0.0003 | 0.0001 |
| Lipid | Sphingomyelins | behenoyl sphingomyelin (d18:1/22:0)* | 0.54 | 0.0003 | 0.0001 |
| Xenobiotics | Food Component/Plant | N1,N10-dicoumaroylspermidine | 0.54 | 0.1372 | 0.0229 |
| Xenobiotics | Food Component/Plant | 2-oxindole-3-acetate | 0.53 | 0.0556 | 0.0103 |
| Nucleotide | Pyrimidine Metabolism, Cytidine containing | 3-methylcytidine | 0.53 | 0.003 | 0.0008 |
| Xenobiotics | Food Component/Plant | daidzin (daidzein 7-O-glucoside) | 0.53 | 0.1448 | 0.0239 |
| Lipid | Phosphatidylcholine (PC) | 1-stearoyl-2-arachidonoyl-GPC (18:0/20:4) | 0.53 | 2.28E-07 | 2.53E-07 |
| Peptide | Dipeptide | prolylglycine | 0.53 | 0.04 | 0.0077 |
| Lipid | Secondary Bile Acid Metabolism | deoxycholate | 0.52 | 0.2681 | 0.0414 |
| Carbohydrate | Glycolysis, Gluconeogenesis, and Pyruvate M | 1,5-anhydroglucitol (1,5-AG) | 0.52 | 3.99E-06 | 2.7E-06 |
| Carbohydrate | Glycolysis, Gluconeogenesis, and Pyruvate M | glycerate | 0.52 | 0.0021 | 0.0006 |
| Lipid | Monoacylglycerol | 1-pentadecanoylglycerol (15:0) | 0.52 | 0.0006 | 0.0002 |
| Amino Acid | Lysine Metabolism | N2-acetyllysine | 0.52 | 3.35E-05 | 1.6E-05 |
| Lipid | Phosphatidylethanolamine (PE) | 1-palmitoyl-2-docosahexaenoyl-GPE (16:0/ | 0.52 | 0.0002 | 9.18E-05 |
| Xenobiotics | Drug - Other | S-carboxymethyl-L-cysteine | 0.52 | 0.0116 | 0.0026 |
| Carbohydrate | Aminosugar Metabolism | diacetylchitobiose | 0.51 | 0.0014 | 0.0004 |
| Lipid | Sphingomyelins | sphingomyelin (d18:2/23:0, d18:1/23:1, d1 | 0.51 | 2.97E-05 | 1.45E-05 |
| Lipid | Diacylglycerol | palmitoyl-docosahexaenoyl-glycerol (16:0/ | 0.51 | 0.0531 | 0.0099 |
| Energy | TCA Cycle | malate | 0.51 | 2.64E-07 | 2.84E-07 |
| Xenobiotics | Food Component/Plant | ferulate | 0.51 | 0.0386 | 0.0075 |
| Lipid | Secondary Bile Acid Metabolism | hyocholate | 0.50 | 0.1195 | 0.0201 |
| Lipid | Fatty Acid, Dicarboxylate | hexadecenedioate (C16:1-DC)* | 0.50 | 0.0014 | 0.0004 |
| Amino Acid | Methionine, Cysteine, SAM and Taurine Met | succinoyltaurine | 0.50 | 3.27E-05 | 1.58E-05 |
| Lipid | Inositol Metabolism | inositol 1-phosphate (I1P) | 0.50 | 0.0936 | 0.0163 |
| Lipid | Fatty Acid, Dicarboxylate | octadecenedioate (C18:1-DC) | 0.49 | 0.0069 | 0.0017 |
| Lipid | Fatty Acid Metabolism (also BCAA Metabolis | butyrylglycine | 0.49 | 0.0291 | 0.0059 |
| Amino Acid | Phenylalanine Metabolism | phenylpyruvate | 0.49 | 0.0025 | 0.0007 |
| Lipid | Lysophospholipid | 1-palmitoyl-GPE (16:0) | 0.49 | 1.88E-05 | 9.78E-06 |
| Lipid | Fatty Acid, Dicarboxylate | 3-hydroxydodecanedioate* | 0.49 | 0.004 | 0.001 |
| Lipid | Fatty Acid, Amino | N-acetyl-2-aminooctanoate* | 0.49 | 0.0004 | 0.0001 |
| Nucleotide | Purine Metabolism, Adenine containing | N6-succinyladenosine | 0.49 | 0.002 | 0.0006 |
| Lipid | Fatty Acid, Dihydroxy | 12,13-DiHOME | 0.49 | 0.0129 | 0.0029 |
| Xenobiotics | Chemical | 1,2,3-benzenetriol sulfate (2) | 0.48 | 0.0931 | 0.0163 |
| Amino Acid | Tyrosine Metabolism | 1-carboxyethyltyrosine | 0.48 | 0.0031 | 0.0008 |
| Lipid | Phosphatidylcholine (PC) | 1-palmitoyl-2-arachidonoyl-GPC (16:0/20:4 | 0.48 | 7.59E-10 | 2.91E-09 |
| Amino Acid | Urea cycle; Arginine and Proline Metabolism | N,N,N-trimethyl-alanylproline betaine (TM | 0.48 | 0.0003 | 0.0001 |
| Lipid | Phosphatidylserine (PS) | 1-stearoyl-2-arachidonoyl-GPS (18:0/20:4) | 0.48 | 1.54E-05 | 8.29E-06 |
| Lipid | Phosphatidylcholine (PC) | 1-myristoyl-2-palmitoyl-GPC (14:0/16:0) | 0.48 | 5.39E-08 | 7.76E-08 |
| Amino Acid | Urea cycle; Arginine and Proline Metabolism | N-acetylarginine | 0.47 | 4.47E-06 | 2.93E-06 |
| Lipid | Phosphatidylinositol (PI) | 1-oleoyl-2-arachidonoyl-GPI (18:1/20:4)* | 0.47 | 9.37E-05 | 3.97E-05 |
| Lipid | Endocannabinoid | N-linolenoyltaurine* | 0.47 | 0.0107 | 0.0025 |
| Lipid | Lysophospholipid | 1-palmitoyl-GPS (16:0)* | 0.46 | 0.0266 | 0.0054 |
| Xenobiotics | Food Component/Plant | mannonate* | 0.46 | 0.0038 | 0.001 |
| Lipid | Secondary Bile Acid Metabolism | taurohyocholate* | 0.46 | 0.017 | 0.0037 |
| Amino Acid | Leucine, Isoleucine and Valine Metabolism | isovalerylglycine | 0.46 | 0.2026 | 0.0322 |
| Lipid | Primary Bile Acid Metabolism | cholate sulfate | 0.46 | 0.0256 | 0.0052 |
| Lipid | Long Chain Saturated Fatty Acid | myristate (14:0) | 0.46 | 0.0014 | 0.0004 |
| Amino Acid | Methionine, Cysteine, SAM and Taurine Met | cysteine | 0.46 | 0.0005 | 0.0002 |
| Peptide | Gamma-glutamyl Amino Acid | gamma-glutamylglycine | 0.45 | 0.0004 | 0.0001 |
| Amino Acid | Guanidino and Acetamido Metabolism | 4-guanidinobutanoate | 0.45 | 0.0004 | 0.0001 |
| Xenobiotics | Food Component/Plant | genistein | 0.45 | 0.0054 | 0.0014 |

| | | | | | |
|---|---|---|---|---|---|
| Lipid | Secondary Bile Acid Metabolism | glycohyodeoxycholate | 0.45 | 0.047 | 0.0088 |
| Amino Acid | Histidine Metabolism | N-acetyl-1-methylhistidine* | 0.44 | 0.0206 | 0.0043 |
| Energy | TCA Cycle | fumarate | 0.44 | 2.16E-06 | 1.64E-06 |
| Lipid | Plasmalogen | 1-(1-enyl-stearoyl)-2-oleoyl-GPE (P-18:0/18 | 0.44 | 0.0001 | 4.29E-05 |
| Amino Acid | Phenylalanine Metabolism | 1-carboxyethylphenylalanine | 0.43 | 0.0004 | 0.0002 |
| Xenobiotics | Food Component/Plant | apigenin glucuronide (1) | 0.43 | 0.0119 | 0.0027 |
| Lipid | Phosphatidylcholine (PC) | 1-palmitoyl-2-oleoyl-GPC (16:0/18:1) | 0.43 | 1.67E-10 | 8.44E-10 |
| Lipid | Fatty Acid, Dicarboxylate | maleate | 0.43 | 2.36E-05 | 1.19E-05 |
| Xenobiotics | Chemical | 3-hydroxypyridine sulfate | 0.43 | 0.0268 | 0.0055 |
| Lipid | Endocannabinoid | N-arachidonoyltaurine | 0.42 | 0.0524 | 0.0098 |
| Amino Acid | Leucine, Isoleucine and Valine Metabolism | 1-carboxyethylleucine | 0.42 | 9.41E-05 | 3.97E-05 |
| Lipid | Fatty Acid, Dicarboxylate | dodecadienoate (12:2)* | 0.42 | 0.0391 | 0.0076 |
| Peptide | Gamma-glutamyl Amino Acid | gamma-glutamylglutamine | 0.42 | 7.81E-05 | 3.39E-05 |
| Carbohydrate | Pentose Metabolism | ribose | 0.42 | 0.0031 | 0.0008 |
| Xenobiotics | Food Component/Plant | glycitein sulfate (2) | 0.42 | 0.0071 | 0.0017 |
| Lipid | Lysophospholipid | 1-stearoyl-GPG (18:0) | 0.42 | 0.0011 | 0.0003 |
| Lipid | Diacylglycerol | linoleoyl-linoleoyl-glycerol (18:2/18:2) [1]* | 0.42 | 0.0078 | 0.0019 |
| Xenobiotics | Food Component/Plant | soyasaponin I | 0.41 | 0.0017 | 0.0005 |
| Carbohydrate | Pentose Metabolism | sedoheptulose | 0.41 | 0.0094 | 0.0022 |
| Xenobiotics | Chemical | 2,4-di-tert-butylphenol | 0.41 | 9.01E-05 | 3.84E-05 |
| Lipid | Diacylglycerol | palmitoyl-oleoyl-glycerol (16:0/18:1) [1]* | 0.41 | 0.0025 | 0.0007 |
| Cofactors and Vitam | Riboflavin Metabolism | riboflavin (Vitamin B2) | 0.41 | 0.0215 | 0.0045 |
| Lipid | Lysophospholipid | 1-oleoyl-GPS (18:1) | 0.40 | 0.0113 | 0.0026 |
| Amino Acid | Urea cycle; Arginine and Proline Metabolism | argininosuccinate | 0.40 | 1.10E-05 | 6.19E-06 |
| Amino Acid | Urea cycle; Arginine and Proline Metabolism | argininate* | 0.40 | 0.0006 | 0.0002 |
| Xenobiotics | Food Component/Plant | deoxymugineic acid | 0.40 | 0.0843 | 0.0149 |
| Peptide | Gamma-glutamyl Amino Acid | gamma-glutamylthreonine | 0.40 | 2.81E-05 | 1.39E-05 |
| Carbohydrate | Fructose, Mannose and Galactose Metabolis | mannose | 0.40 | 0.0013 | 0.0004 |
| Amino Acid | Tyrosine Metabolism | 4-hydroxyphenylpyruvate | 0.40 | 0.0054 | 0.0014 |
| Xenobiotics | Food Component/Plant | N-acetylpyrraline | 0.40 | 0.0109 | 0.0025 |
| Lipid | Medium Chain Fatty Acid | 5-dodecenoate (12:1n7) | 0.40 | 0.0003 | 0.0001 |
| Nucleotide | Purine Metabolism, Guanine containing | guanine | 0.40 | 8.40E-07 | 7.24E-07 |
| Amino Acid | Creatine Metabolism | creatine phosphate | 0.40 | 0.0007 | 0.0002 |
| Lipid | Fatty Acid, Dicarboxylate | suberate (C8-DC) | 0.40 | 0.0054 | 0.0014 |
| Xenobiotics | Benzoate Metabolism | 3-(3-hydroxyphenyl)propionate sulfate | 0.40 | 0.0059 | 0.0014 |
| Nucleotide | Pyrimidine Metabolism, Uracil containing | 5,6-dihydrouracil | 0.39 | 2.47E-06 | 1.82E-06 |
| Lipid | Lysophospholipid | 1-palmitoyl-GPC (16:0) | 0.39 | 6.15E-05 | 2.71E-05 |
| Cofactors and Vitam | Ascorbate and Aldarate Metabolism | ascorbic acid 3-sulfate* | 0.39 | 1.45E-05 | 7.9E-06 |
| Lipid | Lysophospholipid | 1-stearoyl-GPE (18:0) | 0.39 | 1.08E-07 | 1.29E-07 |
| Lipid | Dihydrosphingomyelins | sphingomyelin (d18:0/18:0, d19:0/17:0)* | 0.39 | 3.23E-05 | 1.57E-05 |
| Lipid | Plasmalogen | 1-(1-enyl-palmitoyl)-2-linoleoyl-GPE (P-16: | 0.39 | 1.45E-06 | 1.15E-06 |
| Xenobiotics | Food Component/Plant | genistein sulfate* | 0.39 | 0.0031 | 0.0008 |
| Lipid | Primary Bile Acid Metabolism | glycochenodeoxycholate | 0.39 | 0.033 | 0.0065 |
| Lipid | Secondary Bile Acid Metabolism | taurodeoxycholate | 0.39 | 0.1036 | 0.0178 |
| Lipid | Lysophospholipid | 1-stearoyl-GPS (18:0)* | 0.38 | 0.001 | 0.0003 |
| Xenobiotics | Food Component/Plant | quinate | 0.38 | 0.0598 | 0.011 |
| Lipid | Plasmalogen | 1-(1-enyl-palmitoyl)-2-linoleoyl-GPC (P-16: | 0.38 | 1.92E-06 | 1.47E-06 |
| Lipid | Secondary Bile Acid Metabolism | 7-ketolithocholate | 0.38 | 0.0253 | 0.0052 |
| Carbohydrate | Glycolysis, Gluconeogenesis, and Pyruvate M | pyruvate | 0.38 | 0.0041 | 0.0011 |
| Lipid | Primary Bile Acid Metabolism | glyco-beta-muricholate** | 0.38 | 0.0145 | 0.0032 |
| Lipid | Primary Bile Acid Metabolism | cholate | 0.37 | 0.1117 | 0.019 |
| Lipid | Secondary Bile Acid Metabolism | 6-beta-hydroxylithocholate | 0.37 | 0.0359 | 0.007 |
| Lipid | Eicosanoid | prostaglandin E2 | 0.37 | 0.0003 | 0.0001 |
| Lipid | Endocannabinoid | palmitoyl ethanolamide | 0.37 | 0.0006 | 0.0002 |
| Xenobiotics | Food Component/Plant | 2,8-quinolinediol sulfate | 0.37 | 0.004 | 0.001 |
| Cofactors and Vitam | Riboflavin Metabolism | flavin mononucleotide (FMN) | 0.37 | 0.0003 | 9.76E-05 |
| Amino Acid | Leucine, Isoleucine and Valine Metabolism | N-succinyl-leucine | 0.36 | 0.0044 | 0.0011 |
| Xenobiotics | Food Component/Plant | ergothioneine | 0.36 | 7.44E-07 | 6.63E-07 |

| | | | | | |
|---|---|---|---|---|---|
| Lipid | Lysophospholipid | 1-stearoyl-GPC (18:0) | 0.36 | 1.26E-05 | 6.92E-06 |
| Xenobiotics | Food Component/Plant | soyasaponin II | 0.36 | 0.0005 | 0.0002 |
| Lipid | Monoacylglycerol | 1-myristoylglycerol (14:0) | 0.36 | 0.0021 | 0.0006 |
| Lipid | Monoacylglycerol | 1-linoleoylglycerol (18:2) | 0.36 | 0.0002 | 8.68E-05 |
| Peptide | Gamma-glutamyl Amino Acid | gamma-glutamylglutamate | 0.36 | 0.0002 | 7.59E-05 |
| Cofactors and Vitam | Nicotinate and Nicotinamide Metabolism | nicotinamide N-oxide | 0.36 | 0.0004 | 0.0001 |
| Amino Acid | Polyamine Metabolism | N-acetyl-isoputreanine | 0.35 | 1.38E-08 | 2.42E-08 |
| Amino Acid | Glutamate Metabolism | beta-citrylglutamate | 0.35 | 0.0003 | 0.0001 |
| Partially Characteriz | Partially Characterized Molecules | Flavone derivative C26H28O14 (2)* | 0.35 | 0.0085 | 0.002 |
| Partially Characteriz | Partially Characterized Molecules | Flavone derivative C26H28O14 (4)* | 0.35 | 0.024 | 0.005 |
| Amino Acid | Leucine, Isoleucine and Valine Metabolism | 1-carboxyethylvaline | 0.35 | 4.44E-06 | 2.93E-06 |
| Lipid | Endocannabinoid | linoleoyl ethanolamide | 0.35 | 0.0171 | 0.0037 |
| Amino Acid | Glutathione Metabolism | 5-oxoproline | 0.35 | 0.0002 | 8.69E-05 |
| Lipid | Diacylglycerol | palmitoyl-arachidonoyl-glycerol (16:0/20:4 | 0.35 | 0.0009 | 0.0003 |
| Lipid | Diacylglycerol | linoleoyl-linolenoyl-glycerol (18:2/18:3) [1] | 0.35 | 0.0017 | 0.0005 |
| Amino Acid | Tyrosine Metabolism | 4-hydroxycinnamate sulfate | 0.35 | 0.0109 | 0.0025 |
| Carbohydrate | Fructose, Mannose and Galactose Metabolis | galactonate | 0.34 | 0.0598 | 0.011 |
| Partially Characteriz | Partially Characterized Molecules | Flavone derivative C26H28O14 (3)* | 0.34 | 0.0223 | 0.0046 |
| Carbohydrate | Fructose, Mannose and Galactose Metabolis | mannitol/sorbitol | 0.34 | 0.0106 | 0.0024 |
| Lipid | Ceramides | N-(2-hydroxypalmitoyl)-sphingosine (d18:1 | 0.34 | 0.0001 | 4.5E-05 |
| Lipid | Fatty Acid Metabolism (Acyl Carnitine, Dicar | octadecenedioylcarnitine (C18:1-DC)* | 0.34 | 0.0022 | 0.0006 |
| Xenobiotics | Chemical | S-(3-hydroxypropyl)mercapturic acid (HPM | 0.34 | 0.0041 | 0.0011 |
| Lipid | Ceramides | N-stearoyl-sphingosine (d18:1/18:0)* | 0.34 | 8.14E-06 | 4.82E-06 |
| Carbohydrate | Glycolysis, Gluconeogenesis, and Pyruvate M | dihydroxyacetone phosphate (DHAP) | 0.34 | 7.10E-06 | 4.34E-06 |
| Lipid | Lysophospholipid | 1-linoleoyl-GPS (18:2)* | 0.33 | 0.0022 | 0.0006 |
| Carbohydrate | Pentose Metabolism | arabonate/xylonate | 0.33 | 7.97E-05 | 3.43E-05 |
| Amino Acid | Leucine, Isoleucine and Valine Metabolism | 1-carboxyethylisoleucine | 0.33 | 9.25E-06 | 5.4E-06 |
| Lipid | Primary Bile Acid Metabolism | tauro-beta-muricholate | 0.33 | 0.0089 | 0.0021 |
| Energy | TCA Cycle | alpha-ketoglutarate | 0.33 | 0.002 | 0.0006 |
| Lipid | Phosphatidylserine (PS) | 1-stearoyl-2-oleoyl-GPS (18:0/18:1) | 0.33 | 1.10E-06 | 9.01E-07 |
| Partially Characteriz | Partially Characterized Molecules | Flavone derivative C26H28O14 (1)* | 0.32 | 0.0071 | 0.0017 |
| Cofactors and Vitam | Hemoglobin and Porphyrin Metabolism | biliverdin | 0.32 | 0.001 | 0.0003 |
| Lipid | Ceramides | ceramide (d18:1/20:0, d16:1/22:0, d20:1/1 | 0.32 | 4.82E-06 | 3.1E-06 |
| Xenobiotics | Chemical | 3-hydroxy-2-methylpyridine sulfate | 0.32 | 0.035 | 0.0069 |
| Cofactors and Vitam | Pantothenate and CoA Metabolism | phosphopantetheine | 0.32 | 0.0004 | 0.0002 |
| Lipid | Phosphatidylethanolamine (PE) | 1-stearoyl-2-docosahexaenoyl-GPE (18:0/2 | 0.32 | 3.96E-05 | 1.87E-05 |
| Xenobiotics | Food Component/Plant | gluconate | 0.31 | 0.0006 | 0.0002 |
| Lipid | Lysophospholipid | 1-arachidonoyl-GPI (20:4)* | 0.31 | 1.74E-05 | 9.14E-06 |
| Lipid | Phosphatidylinositol (PI) | 1-palmitoyl-2-arachidonoyl-GPI (16:0/20:4 | 0.31 | 1.01E-08 | 1.96E-08 |
| Xenobiotics | Food Component/Plant | glycitein | 0.31 | 0.0017 | 0.0005 |
| Lipid | Diacylglycerol | oleoyl-oleoyl-glycerol (18:1/18:1) [1]* | 0.31 | 6.04E-05 | 2.68E-05 |
| Lipid | Primary Bile Acid Metabolism | glyco-alpha-muricholate** | 0.31 | 0.0191 | 0.0041 |
| Amino Acid | Leucine, Isoleucine and Valine Metabolism | N-succinyl-isoleucine | 0.30 | 0.0077 | 0.0018 |
| Lipid | Monoacylglycerol | 1-arachidonylglycerol (20:4) | 0.30 | 5.44E-05 | 2.47E-05 |
| Amino Acid | Tryptophan Metabolism | xanthurenate | 0.30 | 0.0894 | 0.0158 |
| Lipid | Lysophospholipid | 2-palmitoyl-GPC (16:0)* | 0.30 | 3.34E-05 | 1.6E-05 |
| Amino Acid | Tryptophan Metabolism | N-acetyltryptophan | 0.30 | 9.73E-06 | 5.57E-06 |
| Xenobiotics | Food Component/Plant | daidzein sulfate (2) | 0.30 | 0.0004 | 0.0001 |
| Xenobiotics | Food Component/Plant | vanillic acid glycine | 0.30 | 0.0038 | 0.001 |
| Lipid | Fatty Acid Metabolism (Acyl Glycine) | N-linoleoylglycine | 0.30 | 0.0063 | 0.0015 |
| Nucleotide | Purine Metabolism, (Hypo)Xanthine/Inosine | allantoic acid | 0.30 | 3.85E-05 | 1.83E-05 |
| Peptide | Gamma-glutamyl Amino Acid | gamma-glutamylcysteine | 0.30 | 0.0226 | 0.0047 |
| Lipid | Fatty Acid Metabolism (Acyl Glycine) | hexanoylglycine | 0.29 | 0.0015 | 0.0004 |
| Xenobiotics | Food Component/Plant | apigenin 7-O(6-malonyl-beta-D-glucoside) | 0.29 | 0.0135 | 0.003 |
| Carbohydrate | Disaccharides and Oligosaccharides | stachyose | 0.29 | 0.0237 | 0.0049 |
| Xenobiotics | Food Component/Plant | apigenin glucuronide (2) | 0.29 | 0.0015 | 0.0004 |
| Lipid | Medium Chain Fatty Acid | caprate (10:0) | 0.29 | 1.91E-06 | 1.47E-06 |
| Xenobiotics | Benzoate Metabolism | 2,4,6-trihydroxybenzoate | 0.29 | 0.0002 | 7.92E-05 |

| | | | | | |
|---|---|---|---|---|---|
| Lipid | Phosphatidylcholine (PC) | 1-palmitoyl-2-palmitoleoyl-GPC (16:0/16:1 | 0.29 | 5.28E-09 | 1.12E-08 |
| Carbohydrate | Disaccharides and Oligosaccharides | sucrose | 0.29 | 0.0002 | 9.26E-05 |
| Amino Acid | Leucine, Isoleucine and Valine Metabolism | 3-methyl-2-oxobutyrate | 0.29 | 0.0002 | 6.49E-05 |
| Lipid | Sphingosines | heptadecasphingosine (d17:1) | 0.28 | 0.0007 | 0.0002 |
| Amino Acid | Polyamine Metabolism | isoputreanine | 0.28 | 5.08E-09 | 1.11E-08 |
| Amino Acid | Lysine Metabolism | fructosyllysine | 0.28 | 0.0008 | 0.0003 |
| Xenobiotics | Food Component/Plant | daidzein sulfate (1) | 0.28 | 0.0023 | 0.0006 |
| Xenobiotics | Food Component/Plant | equol sulfate | 0.28 | 0.0033 | 0.0009 |
| Lipid | Ceramides | ceramide (d18:1/14:0, d16:1/16:0)* | 0.28 | 1.80E-08 | 3.04E-08 |
| Lipid | Medium Chain Fatty Acid | cis-4-decenoate (10:1n6)* | 0.28 | 0.0003 | 0.0001 |
| Amino Acid | Polyamine Metabolism | spermine | 0.28 | 0.0463 | 0.0087 |
| Partially Characteriz | Partially Characterized Molecules | pentose acid* | 0.27 | 0.0181 | 0.0039 |
| Xenobiotics | Food Component/Plant | soyasaponin III | 0.27 | 1.11E-05 | 6.23E-06 |
| Peptide | Gamma-glutamyl Amino Acid | gamma-glutamylalanine | 0.27 | 2.16E-06 | 1.64E-06 |
| Lipid | Glycosyl PE | 1-stearoyl-2-linoleoyl-glycosyl-GPE (18:0/1 | 0.27 | 0.0012 | 0.0004 |
| Amino Acid | Glycine, Serine and Threonine Metabolism | betaine aldehyde | 0.26 | 0.0002 | 7.08E-05 |
| Lipid | Galactosyl Glycerolipids | digalactosylglycerol* | 0.26 | 0.0007 | 0.0002 |
| Carbohydrate | Disaccharides and Oligosaccharides | raffinose | 0.26 | 0.0169 | 0.0037 |
| Cofactors and Vitam | Ascorbate and Aldarate Metabolism | ascorbic acid 2-sulfate | 0.26 | 3.82E-07 | 3.97E-07 |
| Lipid | Phosphatidylethanolamine (PE) | 1-stearoyl-2-arachidonoyl-GPE (18:0/20:4) | 0.26 | 2.88E-07 | 3.04E-07 |
| Lipid | Ceramides | N-palmitoyl-sphingosine (d18:1/16:0) | 0.26 | 4.86E-08 | 7.16E-08 |
| Xenobiotics | Food Component/Plant | 2-ketogluconate | 0.26 | 0.0152 | 0.0033 |
| Xenobiotics | Food Component/Plant | caffeic acid sulfate | 0.26 | 0.0063 | 0.0015 |
| Lipid | Secondary Bile Acid Metabolism | 6-oxolithocholate | 0.25 | 0.0032 | 0.0009 |
| Lipid | Eicosanoid | 13,14-dihydro-15-keto-prostaglandin A2 | 0.25 | 5.29E-06 | 3.36E-06 |
| Lipid | Inositol Metabolism | pinitol | 0.25 | 0.0069 | 0.0017 |
| Lipid | Sphingomyelins | sphingomyelin (d18:2/21:0, d16:2/23:0)* | 0.25 | 7.29E-09 | 1.46E-08 |
| Lipid | Monoacylglycerol | 1-dihomo-linolenylglycerol (20:3) | 0.25 | 1.59E-05 | 8.5E-06 |
| Peptide | Gamma-glutamyl Amino Acid | gamma-glutamylvaline | 0.25 | 2.45E-06 | 1.82E-06 |
| Amino Acid | Urea cycle; Arginine and Proline Metabolism | 2-oxoarginine* | 0.25 | 0.0012 | 0.0004 |
| Lipid | Phosphatidylcholine (PC) | 1-oleoyl-2-docosahexaenoyl-GPC (18:1/22 | 0.25 | 3.92E-07 | 4.01E-07 |
| Lipid | Medium Chain Fatty Acid | laurate (12:0) | 0.25 | 3.12E-06 | 2.17E-06 |
| Xenobiotics | Food Component/Plant | ferulic acid 4-sulfate | 0.25 | 0.0002 | 9.23E-05 |
| Amino Acid | Glutathione Metabolism | cysteine-glutathione disulfide | 0.25 | 0.0002 | 7.36E-05 |
| Lipid | Phosphatidylethanolamine (PE) | 1-palmitoyl-2-arachidonoyl-GPE (16:0/20:4 | 0.25 | 8.19E-10 | 3.05E-09 |
| Lipid | Lysophospholipid | 1-palmitoleoyl-GPC (16:1)* | 0.24 | 6.79E-06 | 4.19E-06 |
| Lipid | Monoacylglycerol | 1-linolenoylglycerol (18:3) | 0.24 | 0.0007 | 0.0002 |
| Lipid | Monoacylglycerol | 1-dihomo-linoleoylglycerol (20:2) | 0.24 | 0.0003 | 0.0001 |
| Lipid | Phosphatidylcholine (PC) | 1-stearoyl-2-oleoyl-GPC (18:0/18:1) | 0.24 | 5.71E-12 | 6.02E-11 |
| Lipid | Sphingomyelins | tricosanoyl sphingomyelin (d18:1/23:0)* | 0.24 | 9.46E-08 | 1.19E-07 |
| Lipid | Monoacylglycerol | 1-eicosapentaenoylglycerol (20:5)* | 0.24 | 2.11E-05 | 1.08E-05 |
| Lipid | Primary Bile Acid Metabolism | beta-muricholate | 0.23 | 0.0457 | 0.0086 |
| Lipid | Monoacylglycerol | 1-oleoylglycerol (18:1) | 0.23 | 5.18E-05 | 2.38E-05 |
| Peptide | Gamma-glutamyl Amino Acid | gamma-glutamylmethionine | 0.23 | 9.06E-07 | 7.65E-07 |
| Lipid | Monoacylglycerol | 2-palmitoylglycerol (16:0) | 0.23 | 5.34E-05 | 2.43E-05 |
| Lipid | Fatty Acid, Dicarboxylate | 2-hydroxysebacate | 0.23 | 0.0004 | 0.0001 |
| Peptide | Gamma-glutamyl Amino Acid | gamma-glutamylserine | 0.23 | 2.32E-05 | 1.18E-05 |
| Lipid | Secondary Bile Acid Metabolism | taurolithocholate | 0.23 | 0.0029 | 0.0008 |
| Lipid | Lysophospholipid | 1-arachidonoyl-GPC (20:4n6)* | 0.22 | 1.80E-10 | 8.75E-10 |
| Peptide | Gamma-glutamyl Amino Acid | gamma-glutamylhistidine | 0.22 | 1.78E-05 | 9.31E-06 |
| Cofactors and Vitam | Nicotinate and Nicotinamide Metabolism | nicotinate | 0.22 | 0.0151 | 0.0033 |
| Carbohydrate | Fructose, Mannose and Galactose Metabolis | 2-ketogulonate | 0.22 | 0.0185 | 0.004 |
| Lipid | Secondary Bile Acid Metabolism | taurohyodeoxycholic acid | 0.22 | 0.0035 | 0.0009 |
| Xenobiotics | Food Component/Plant | nicotianamine | 0.21 | 0.0793 | 0.0142 |
| Lipid | Fatty Acid, Monohydroxy | 2-hydroxybehenate | 0.21 | 3.49E-06 | 2.4E-06 |
| Lipid | Lysophospholipid | 2-stearoyl-GPE (18:0)* | 0.21 | 5.30E-07 | 4.97E-07 |
| Lipid | Monoacylglycerol | 2-docosahexaenoylglycerol (22:6)* | 0.21 | 0.0002 | 6.34E-05 |
| Carbohydrate | Aminosugar Metabolism | glucuronate | 0.21 | 1.92E-07 | 2.19E-07 |

| | | | | | |
|---|---|---|---|---|---|
| Lipid | Primary Bile Acid Metabolism | taurochenodeoxycholate | 0.21 | 0.0048 | 0.0012 |
| Lipid | Lysophospholipid | 1-oleoyl-GPC (18:1) | 0.20 | 1.14E-05 | 6.34E-06 |
| Peptide | Gamma-glutamyl Amino Acid | gamma-glutamylphenylalanine | 0.20 | 2.65E-06 | 1.9E-06 |
| Peptide | Gamma-glutamyl Amino Acid | gamma-glutamyltyrosine | 0.20 | 2.45E-06 | 1.82E-06 |
| Amino Acid | Leucine, Isoleucine and Valine Metabolism | 3-methyl-2-oxovalerate | 0.20 | 2.10E-05 | 1.08E-05 |
| Xenobiotics | Food Component/Plant | genistein glucuronide* | 0.20 | 0.0004 | 0.0001 |
| Lipid | Diacylglycerol | stearoyl-docosahexaenoyl-glycerol (18:0/2 | 0.20 | 5.77E-05 | 2.59E-05 |
| Lipid | Secondary Bile Acid Metabolism | tauroursodeoxycholic acid sulfate (2) | 0.19 | 0.0002 | 8.74E-05 |
| Lipid | Phosphatidylcholine (PC) | 1-palmitoyl-2-linoleoyl-GPC (16:0/18:2) | 0.19 | 2.21E-12 | 3.12E-11 |
| Amino Acid | Glutamate Metabolism | S-1-pyrroline-5-carboxylate | 0.19 | 0.0017 | 0.0005 |
| Lipid | Monoacylglycerol | 1-heptadecenoylglycerol (17:1)* | 0.18 | 8.70E-05 | 3.72E-05 |
| Lipid | Fatty Acid, Monohydroxy | 16-hydroxypalmitate | 0.18 | 8.97E-08 | 1.15E-07 |
| Peptide | Gamma-glutamyl Amino Acid | gamma-glutamylisoleucine* | 0.18 | 6.72E-06 | 4.17E-06 |
| Lipid | Phosphatidylethanolamine (PE) | 1-oleoyl-2-docosahexaenoyl-GPE (18:1/22: | 0.18 | 6.06E-08 | 8.33E-08 |
| Lipid | Phosphatidylcholine (PC) | 1-myristoyl-2-arachidonoyl-GPC (14:0/20:4 | 0.18 | 4.58E-07 | 4.53E-07 |
| Lipid | Phosphatidylserine (PS) | 1-palmitoyl-2-oleoyl-GPS (16:0/18:1) | 0.17 | 4.72E-07 | 4.6E-07 |
| Peptide | Gamma-glutamyl Amino Acid | gamma-glutamylleucine | 0.17 | 2.58E-06 | 1.86E-06 |
| Lipid | Phosphatidylcholine (PC) | 1-palmitoyl-2-dihomo-linolenoyl-GPC (16:0 | 0.17 | 1.95E-12 | 3.12E-11 |
| Xenobiotics | Food Component/Plant | daidzein 7-O-glucuronide | 0.17 | 0.0001 | 5.92E-05 |
| Lipid | Phosphatidylcholine (PC) | 1-stearoyl-2-linoleoyl-GPC (18:0/18:2)* | 0.17 | 2.12E-12 | 3.12E-11 |
| Lipid | Secondary Bile Acid Metabolism | tauroursodeoxycholate | 0.17 | 0.0043 | 0.0011 |
| Lipid | Primary Bile Acid Metabolism | chenodeoxycholate | 0.16 | 0.0091 | 0.0022 |
| Xenobiotics | Food Component/Plant | glycitein glucuronide (2)* | 0.16 | 0.0002 | 7.81E-05 |
| Lipid | Phosphatidylethanolamine (PE) | 1-oleoyl-2-arachidonoyl-GPE (18:1/20:4)* | 0.16 | 1.35E-11 | 9.49E-11 |
| Lipid | Secondary Bile Acid Metabolism | 12-dehydrocholate | 0.16 | 0.0012 | 0.0004 |
| Lipid | Lysophospholipid | 1-lignoceroyl-GPC (24:0) | 0.15 | 3.93E-07 | 4.01E-07 |
| Lipid | Lysophospholipid | 1-oleoyl-GPE (18:1) | 0.15 | 1.29E-09 | 3.68E-09 |
| Amino Acid | Leucine, Isoleucine and Valine Metabolism | 4-methyl-2-oxopentanoate | 0.15 | 0.0025 | 0.0007 |
| Lipid | Diacylglycerol | stearoyl-arachidonoyl-glycerol (18:0/20:4) | 0.15 | 5.90E-05 | 2.63E-05 |
| Lipid | Diacylglycerol | oleoyl-linoleoyl-glycerol (18:1/18:2) [1] | 0.15 | 0.0002 | 7.89E-05 |
| Lipid | Phosphatidylcholine (PC) | 1-palmitoyl-2-gamma-linolenoyl-GPC (16:0 | 0.14 | 0.0003 | 0.0001 |
| Xenobiotics | Food Component/Plant | histidine betaine (hercynine)* | 0.14 | 0.0006 | 0.0002 |
| Amino Acid | Glutamate Metabolism | carboxyethyl-GABA | 0.14 | 6.02E-07 | 5.54E-07 |
| Lipid | Sphingomyelins | sphingomyelin (d18:1/21:0, d17:1/22:0, d1 | 0.14 | 9.71E-09 | 1.92E-08 |
| Lipid | Lysophospholipid | 1-linoleoyl-GPE (18:2)* | 0.14 | 5.60E-08 | 7.97E-08 |
| Xenobiotics | Food Component/Plant | naringenin | 0.14 | 0.3438 | 0.0514 |
| Xenobiotics | Food Component/Plant | dihydrokaempferol | 0.14 | 0.2564 | 0.0397 |
| Xenobiotics | Food Component/Plant | enterolactone sulfate | 0.13 | 1.20E-05 | 6.68E-06 |
| Peptide | Gamma-glutamyl Amino Acid | gamma-glutamyl-alpha-lysine | 0.13 | 9.70E-06 | 5.57E-06 |
| Peptide | Gamma-glutamyl Amino Acid | gamma-glutamyltryptophan | 0.13 | 6.40E-07 | 5.79E-07 |
| Lipid | Ceramides | ceramide (d18:1/17:0, d17:1/18:0)* | 0.13 | 5.14E-08 | 7.47E-08 |
| Cofactors and Vitam | Pantothenate and CoA Metabolism | pantetheine | 0.13 | 0.0005 | 0.0002 |
| Lipid | Diacylglycerol | linoleoyl-linoleoyl-glycerol (18:2/18:2) [2]* | 0.13 | 0.0001 | 5.99E-05 |
| Lipid | Diacylglycerol | linoleoyl-linolenoyl-glycerol (18:2/18:3) [2] | 0.13 | 0.0002 | 7.08E-05 |
| Lipid | Secondary Bile Acid Metabolism | 7-ketodeoxycholate | 0.12 | 0.0058 | 0.0014 |
| Lipid | Long Chain Monounsaturated Fatty Acid | myristoleate (14:1n5) | 0.12 | 9.52E-10 | 3.32E-09 |
| Lipid | Fatty Acid Metabolism (Acyl Carnitine, Dicar | adipoylcarnitine (C6-DC) | 0.12 | 8.26E-06 | 4.85E-06 |
| Lipid | Sphingomyelins | sphingomyelin (d18:1/19:0, d19:1/18:0)* | 0.12 | 6.20E-10 | 2.53E-09 |
| Xenobiotics | Food Component/Plant | equol glucuronide | 0.12 | 0.0004 | 0.0001 |
| Lipid | Monoacylglycerol | 2-linoleoylglycerol (18:2) | 0.12 | 5.03E-07 | 4.79E-07 |
| Lipid | Phosphatidylcholine (PC) | 1-oleoyl-2-linoleoyl-GPC (18:1/18:2)* | 0.12 | 1.24E-12 | 3.12E-11 |
| Lipid | Secondary Bile Acid Metabolism | ursocholate | 0.11 | 0.0009 | 0.0003 |
| Lipid | Lysophospholipid | 1-linoleoyl-GPI (18:2)* | 0.11 | 6.05E-06 | 3.79E-06 |
| Lipid | Monoacylglycerol | 2-oleoylglycerol (18:1) | 0.11 | 2.88E-06 | 2.02E-06 |
| Cofactors and Vitam | Vitamin A Metabolism | carotene diol (1) | 0.11 | 4.75E-05 | 2.19E-05 |
| Lipid | Lysophospholipid | 1-linolenoyl-GPC (18:3)* | 0.11 | 1.34E-09 | 3.68E-09 |
| Lipid | Lysophospholipid | 1-linoleoyl-GPC (18:2) | 0.11 | 5.85E-08 | 8.14E-08 |
| Lipid | Phosphatidylethanolamine (PE) | 1-palmitoyl-2-oleoyl-GPE (16:0/18:1) | 0.11 | 1.05E-08 | 1.98E-08 |

| | | | | | |
|---|---|---|---|---|---|
| Lipid | Monoacylglycerol | 2-myristoylglycerol (14:0) | 0.11 | 4.12E-07 | 4.17E-07 |
| Lipid | Primary Bile Acid Metabolism | alpha-muricholate | 0.10 | 0.0008 | 0.0002 |
| Lipid | Monoacylglycerol | 2-arachidonoylglycerol (20:4) | 0.10 | 7.62E-07 | 6.75E-07 |
| Lipid | Monoacylglycerol | 2-eicosapentaenoylglycerol (20:5)* | 0.10 | 2.88E-06 | 2.02E-06 |
| Lipid | Sterol | campesterol | 0.10 | 1.13E-09 | 3.66E-09 |
| Lipid | Phosphatidylethanolamine (PE) | 1-stearoyl-2-oleoyl-GPE (18:0/18:1) | 0.10 | 3.43E-09 | 8E-09 |
| Lipid | Fatty Acid, Branched | 18-methylnonadecanoate (i20:0) | 0.09 | 1.11E-07 | 1.32E-07 |
| Lipid | Monoacylglycerol | 1-palmitoleoylglycerol (16:1)* | 0.09 | 1.69E-06 | 1.31E-06 |
| Xenobiotics | Food Component/Plant | ferulylglycine (1) | 0.09 | 1.07E-06 | 8.76E-07 |
| Lipid | Sterol | beta-sitosterol | 0.09 | 1.34E-09 | 3.68E-09 |
| Cofactors and Vitam | Thiamine Metabolism | 5-(2-Hydroxyethyl)-4-methylthiazole | 0.09 | 0.0054 | 0.0014 |
| Lipid | Diacylglycerol | palmitoyl-linoleoyl-glycerol (16:0/18:2) [1] | 0.08 | 3.69E-06 | 2.53E-06 |
| Amino Acid | Histidine Metabolism | formiminoglutamate | 0.08 | 6.28E-07 | 5.72E-07 |
| Lipid | Ceramides | N-palmitoyl-heptadecasphingosine (d17:1/ | 0.08 | 1.20E-09 | 3.68E-09 |
| Lipid | Diacylglycerol | stearoyl-docosahexaenoyl-glycerol (18:0/2 | 0.08 | 1.28E-06 | 1.04E-06 |
| Xenobiotics | Food Component/Plant | ferulylglycine (2) | 0.08 | 1.52E-06 | 1.2E-06 |
| Lipid | Secondary Bile Acid Metabolism | ursodeoxycholate | 0.07 | 0.0013 | 0.0004 |
| Lipid | Diacylglycerol | diacylglycerol (14:0/18:1, 16:0/16:1) [2]* | 0.07 | 1.03E-08 | 1.98E-08 |
| Lipid | Diacylglycerol | palmitoyl-linolenoyl-glycerol (16:0/18:3) [2 | 0.07 | 7.79E-12 | 7.23E-11 |
| Lipid | Phosphatidylcholine (PC) | 1-linoleoyl-2-arachidonoyl-GPC (18:2/20:4 | 0.07 | 2.22E-09 | 5.62E-09 |
| Xenobiotics | Food Component/Plant | naringenin 7-glucuronide | 0.07 | 0.0003 | 0.0001 |
| Lipid | Diacylglycerol | diacylglycerol (12:0/18:1, 14:0/16:1, 16:0/ | 0.06 | 7.20E-08 | 9.6E-08 |
| Lipid | Monoacylglycerol | 2-heptadecenoylglycerol (17:1)* | 0.06 | 7.22E-07 | 6.48E-07 |
| Carbohydrate | Pentose Metabolism | arabinose | 0.06 | 0.0003 | 0.0001 |
| Lipid | Diacylglycerol | diacylglycerol (16:1/18:2 [2], 16:0/18:3 [1] | 0.06 | 4.70E-05 | 2.18E-05 |
| Lipid | Phosphatidylethanolamine (PE) | 1-linoleoyl-2-arachidonoyl-GPE (18:2/20:4) | 0.06 | 2.57E-08 | 4.12E-08 |
| Lipid | Phosphatidylinositol (PI) | 1-palmitoyl-2-linoleoyl-GPI (16:0/18:2) | 0.06 | 7.50E-10 | 2.91E-09 |
| Lipid | Phosphatidylserine (PS) | 1-stearoyl-2-linoleoyl-GPS (18:0/18:2) | 0.06 | 2.51E-12 | 3.17E-11 |
| Lipid | Diacylglycerol | oleoyl-linoleoyl-glycerol (18:1/18:2) [2] | 0.06 | 1.61E-05 | 8.55E-06 |
| Lipid | Monoacylglycerol | 1-myristoleoylglycerol (14:1) | 0.05 | 1.03E-05 | 5.85E-06 |
| Lipid | Phosphatidylglycerol (PG) | 1-palmitoyl-2-oleoyl-GPG (16:0/18:1) | 0.05 | 1.74E-09 | 4.58E-09 |
| Lipid | Diacylglycerol | stearoyl-arachidonoyl-glycerol (18:0/20:4) | 0.05 | 2.20E-05 | 1.12E-05 |
| Lipid | Secondary Bile Acid Metabolism | taurochenodeoxycholic acid (7 or 27)-sulfa | 0.05 | 5.62E-07 | 5.23E-07 |
| Partially Characteriz | Partially Characterized Molecules | branched-chain, straight-chain, or cyclopro | 0.04 | 1.53E-10 | 8.1E-10 |
| Lipid | Diacylglycerol | palmitoyl-linoleoyl-glycerol (16:0/18:2) [2] | 0.04 | 8.07E-06 | 4.8E-06 |
| Lipid | Phosphatidylethanolamine (PE) | 1-palmitoyl-2-linoleoyl-GPE (16:0/18:2) | 0.04 | 2.22E-12 | 3.12E-11 |
| Lipid | Diacylglycerol | stearoyl-linoleoyl-glycerol (18:0/18:2) [2]* | 0.04 | 9.96E-10 | 3.32E-09 |
| Lipid | Phosphatidylethanolamine (PE) | 1,2-dilinoleoyl-GPE (18:2/18:2)* | 0.04 | 4.22E-08 | 6.51E-08 |
| Lipid | Monoacylglycerol | 2-palmitoleoylglycerol (16:1)* | 0.03 | 7.13E-08 | 9.6E-08 |
| Lipid | Diacylglycerol | palmitoyl-oleoyl-glycerol (16:0/18:1) [2]* | 0.03 | 2.99E-13 | 9.47E-12 |
| Lipid | Phosphatidylinositol (PI) | 1-stearoyl-2-linoleoyl-GPI (18:0/18:2) | 0.03 | 3.55E-10 | 1.6E-09 |
| Lipid | Phosphatidylethanolamine (PE) | 1,2-dioleoyl-GPE (18:1/18:1) | 0.03 | 9.90E-10 | 3.32E-09 |
| Lipid | Phosphatidylcholine (PC) | 1,2-dilinoleoyl-GPC (18:2/18:2) | 0.03 | 6.02E-10 | 2.53E-09 |
| Lipid | Diacylglycerol | oleoyl-oleoyl-glycerol (18:1/18:1) [2]* | 0.03 | 1.65E-08 | 2.87E-08 |
| Lipid | Phosphatidylethanolamine (PE) | 1-oleoyl-2-linoleoyl-GPE (18:1/18:2)* | 0.02 | 1.78E-11 | 1.13E-10 |
| Lipid | Phosphatidylcholine (PC) | 1-palmitoleoyl-2-linolenoyl-GPC (16:1/18:3 | 0.02 | 4.28E-06 | 2.84E-06 |
| Lipid | Phosphatidylethanolamine (PE) | 1-stearoyl-2-linoleoyl-GPE (18:0/18:2)* | 0.02 | 5.13E-11 | 2.95E-10 |
| Lipid | Phosphatidylcholine (PC) | 1-linoleoyl-2-linolenoyl-GPC (18:2/18:3)* | 0.01 | 1.95E-13 | 8.22E-12 |
| Amino Acid | Methionine, Cysteine, SAM and Taurine Met | cysteine s-sulfate | 0.01 | 7.79E-07 | 6.8E-07 |

**SUPPLEMENTAL TABLE 2: FLOW CYTOMETRY ANTIBODIES AND OTHER REAGENTS**

| Target | Clone number | Fluorochrome(s) | Vendor |
|---|---|---|---|
| CD11b | M1/70 | e450, APC, APCCy7 | eBioscience |
| CD4 | RM4-5 | e450 | BD Pharmingen |
| CD45.1 | A20 | PE | BD Pharmingen |
| Gr-1 | RB6-8C5 | APC | BD Pharmingen |
| Hematopoietic lineage cocktail | N/A | e450 | Invitrogen |
| KLRG1 | 2F1 | APCe780 | Invitrogen |
| Live/Dead fixable aqua | N/A | Pacific Orange | Invitrogen |
| Sca-1 (Ly-6A/E) | D7 | PerCP-Cy5 | Invitrogen |
| TCRβ | H57-597 | APC, FITC | BD Pharmingen |
| Thy1.1 | OX-7 | FITC | BD Pharmingen |
| Dnk anti-mouse | N/A | AF488 | Abcam |
| Goat anti-mouse | N/A | AF594 | Abcam |
| Goat anti-rabbit | N/A | AF594, AF488, FITC | Abcam |
| Goat anti-rat | N/A | Cy5 | Abcam |
| Mouse anti-rat | N/A | biotin | Invitrogen |
| Goat anti-rabbit | N/A | biotin | Thermo Scientific |
| Mouse IgG1 | MOPC-21 | N/A | BioXCell |
| Mouse IgG2a | C140SF9 | N/A | N/A |
| Mouse IgG2a | C1.18.4 | N/A | BioXCell |
| Rat IgG | N/A | N/A | Jackson Immuno Res |
| Purified anti-CD4 | GK1.5 | N/A | BioXCell |
| Purified anti-IL9 | MM9C1 | N/A | N/A |
| Purified anti- IL13 | | N/A | UCB Pharma |
| Purified anti-GATA3 | L50-823 | N/A | BD Pharmingen |
| Purified anti-KLRG1 | 2F1 | N/A | BD Pharmingen |
| Purified anti-ST2 | 245707 | N/A | R&D |
| Rabbit anti-mouse DCAMKL-1 | N/A | N/A | Abcam |
| Rabbit anti-mouse E-Cadherin | N/A | N/A | Proteintech |
| Rabbit anti-mouse IL9 | EPR23484-151 | N/A | Abcam |
| Rat anti-mouse MCP-1 | RF6.1 | N/A | Invitrogen |
| Rat anti-mouse CD3 | CD3-12 | N/A | Abcam |
| Rat anti-mouse IL33 | 396118 | N/A | R&D |
| Rat anti-mouse IL33 | Polyclonal | N/A | R&D |
| Rat anti-mouse MCP-1/Mcpt1 | Polyclonal | N/A | R&D |
| Mouse anti-ß-actin | E4D9Z | N/A | Cell Signaling |

* APC=Allophycocyanin, AF=AlexaFluor, Cy=Cyanine, PE= Phycoerythrin, FITC= Fluorescein Isothiocyanate, PerCP= Peridinin-Chlorophyll-protein

**SUPPLEMENTAL TABLE 3: QRT-PCR PRIMERS (LISTED 5'-3')**

| Target | Forward | Reverse |
|---|---|---|
| *Areg* | GCAGATACATCGAGAACCTGG | CTGCAATCTTGGATAGGTCCTTG |
| *Crypt1* | TCAAGAGGCTGCAAAGGAAGA GAAC | TGGTCTCCATGTTCAGCGACAGC |
| *DCLK* | CAAGCCAGCCATGTCGTTC | TTCCTTTGAAGTAGCGGTCAC |
| *Fcer1a* | GCCCCGTCTCCATTAG | CAATAACCCCGTGTCC |
| *GMCSF* | TTTACTTTTCCTGGGCAT | TAGCTGGCTGTCATGTTCAA |
| *Gob-5* | ACTAAAGGTGGCCTACCTCCAA | GGAGGTGACAGTCAAGGTGAG |
| *IL13* | GCTTATTGAGGAGCTGAGCAAC A | GGCCAGGTCCACACTCCATA |
| *IL17rb* | CCATCCCTCCAGATGACAAC | TGCTCCTTCCTTGCCTCCAAGTTA |
| *IL22* | TCTGAGAAATGCTTGCGTCTGA | ACTGAGCCAGGTTTCATGTGAA |
| *IL25* | ACAGGGACTTGAATCGGGTC | TGGTAAAGTGGGACGGAGTTG |
| *IL33* | GGTGTGGATGGGAAGAAGCTG | GAGGACTTTTTGTGAAGGACG |
| *IL4* | ATCATCGGCATTTTGAACGAGG TC | ACCTTGGAAGCCCTACAGACG |
| *IL5* | GATGAGGCTTCCTGTCCCTACT | TGACAGGTTTTGGAATAGCATTTC C |
| *IL6* | GTTXCTCTGGGAAATCGTGGA | TCCAGTTTGGTAGCATCCATC |
| *IL9* | CATCAGTGTCTCTCCGTCCCAA CTGATG | GATTTCTGTGTGGCATTGGTCAG |
| *IL9R* | ATGGGACAGGAACAGGTCAG | AGGTCACTCCAACGATACGG |
| *IFNγ* | TCAAGTGGCATAGATGTGGAAG AA | TGGCTCTGCAGGATTTTCATG |
| *mMCPs* (1,2,4chymase) | GCTGGAGCTGAGGAGATT | GGTGAAGACTGCAGGGG |
| *mMCP-7* (Tryptase) | CCTCACTGTGTCCAAATGCTA | CCTCCTGCCTCAGAGACC |
| *Sucnr1* | GGGGACCTATGGAGATGTTCT | GCCAGCGAGATTAAAATGGCAA |
| *sPLA2* | AGGATTCCCCCAAGATGCCAC | CAGCCGTTTCTGACAGGAGTTCTG G |
| *ST2* | TCTCTTCTGGACCCTACCTCAG | TACTGCCCTCCGTAACTGTCA |
| *TNFα* | CTTCTGTCTACTGAACTTCGGG | CAGGCTTGTCACTCGAATTTTG |
| *TSLP* | AGCTTGTCTCCTGAAAATCGAG | AGGTTTGATTCAGGCAGATGTT |
